# Colonial ground nesting by *Archaeopteryx* suggests wing evolution in primal association with nesting and the ground up evolution of flight

**DOI:** 10.1101/2023.05.30.542892

**Authors:** M. Jorge Guimarães

## Abstract

Following an investigation into the hypothesis that the iconic Berlin specimen of *Archaeopteryx* fossilized in nesting position, which led to the discovery not only of its association with soft eggs and several hatchlings, but also similar findings in a second *Archaeopteryx* specimen, an attempt to characterize the entire Berlin specimen nest and estimate its number of eggs is reported here.

The Berlin specimen arranged and brooded its eggs on the ground. Its clutch size appears to have exceeded one hundred eggs. Egg littering found not only in its fossil bed but also in the sediment layer immediately above it, inclusively with evidence that a subsequent generation nested over the specimen, is consistent with repeated usage of a ground nesting site.

All *Archaeopteryx* specimens fossilized in different views of a similar pose that is compatible with a nesting posture, and evidence of eggs of consistent size with the 2D outlines of 3D flattened eggs is present not only in the Berlin, Teylers, Thermopolis and Maxberg specimens, but also in the isolated *Archaeopteryx* feather fossil.

In addition, egg and hatchling littering are present in the Berlin, Teylers and isolated *Archaeopteryx* feather fossils.

Taken together, these findings are indicative of colonial ground nesting behavior by *Archaeopteryx* in Solnhofen.

Egg littering, eggs dorsal to the Berlin specimen torso and limb rotations in the London and Thermopolis *Archaeopteryx* specimens can all be explained by nesting in reentrances located at the margins or in sand banks of marine lagoons in Solnhofen, which would have been flooded, causing the subsequent collapse of the nest and the still-life preservation of its content.

The discovery of colonial ground nesting in a winged Jurassic bird relative favors the evolution of birds from the ground up and suggests that wings and their elongated feathers were primarily associated with ground nest protection and only secondarily with flight.

## Introduction

The Berlin *Archaeopteryx*, being the best preserved specimen of a Jurassic winged and feathered species with a bony tail, clawed wings and toothed beak, features that do not coexist in extant animals, became the Mona Lisa of fossils and the embodiment of the theory of evolution for seeming to represent a transitional stage into today’s birds (*1,2*).

The main slab of this fossil is on public display at Naturkundemuseum Berlin, the Berlin Museum of Natural History, encased in its exhibition mount, whereas its much lesser known counterslab is kept at the collection room of the same museum. The dorsal surface of the main slab, where the Berlin specimen skeleton is observable, has been extensively studied in regards to anatomical details of the specimen and their implications for the evolution of birds and flight. However, the ventral surface of the said slab and the counterslab are rarely observed.

All *Archaeopteryx* fossils were found in the Solnhofen Formation, Germany (*1,3-5*), widely considered as a purely marine environment despite the fact that it contains, in addition to a dozen specimens of the winged *Archaeopteryx* species, numerous pterosaurs, animals and plants that would have lived on dry land, such as lizards, turtles without aquatic adaptations, small dinosaurs (*1,5-7*), inclusively in association with eggs (*7*), and dinosaur tracks (*8*). To explain these findings in a marine environment, the often fully articulated skeletons of these land animals were presumed to have been transported from the Solnhofen archipelago, after their death, by rain waters and to have sunk into shallow lagoons despite the absence of settling marks and the speculative nature of this explanation (*1,9*).

Based on the resemblance of the fossilized pose of the Berlin specimen’s fully articulated skeleton with that of a nesting bird, the possibility that it died nesting near Solnhofen’s lagoons and fossilized with minor disturbances to its posture and original location was investigated, resulting in the identification of soft eggs and hatchlings in this fossil, findings that were corroborated in the Teylers specimen fossil (*10–12*).

An analysis of the entirety of the two fossil slabs that encase Berlin specimen’s body using macrophotography, transillumination, stereomicroscopy, CT scan analysis, including three-dimensional software analysis of computed tomography data, in addition to 3D software modeling, is reported here in an attempt to characterize the entire Berlin specimen nest and estimate its number of eggs. Additional studies of the Teylers, Thermopolis and isolated *Archaeopteryx* feather fossils are also described in this report.

A new interpretation of the fossilized pose of all *Archaeopteryx* specimens as that of nesting animals will be presented here, together with a nest architecture and plausible death mechanisms that can explain all data that is now available on these fossils.

## Materials and Methods

### Experimental approach to the study of the Berlin specimen of A*rchaeopteryx* based on the nesting hypothesis

This specimen’s reinterpretation as the dorso-lateral view of a nesting animal requires an alternative explanation for the previously accepted relative position of its fossil slabs in the Solnhofen Plattenkalk that was based solely on the finding that most, but not all, Solnhofen fossilized skeletons reside in the upper slab after separation because the original position of the slabs was not documented (*1, 10–12*).

According to the hypothesis explored here, the specimen’s skeleton, dorsally exposed in the main slab, reposed with its ventral side – not with its back –, against the sediment floor, whereas the counterslab would have resided above the specimen (Figure 1).

**Figure 1.**
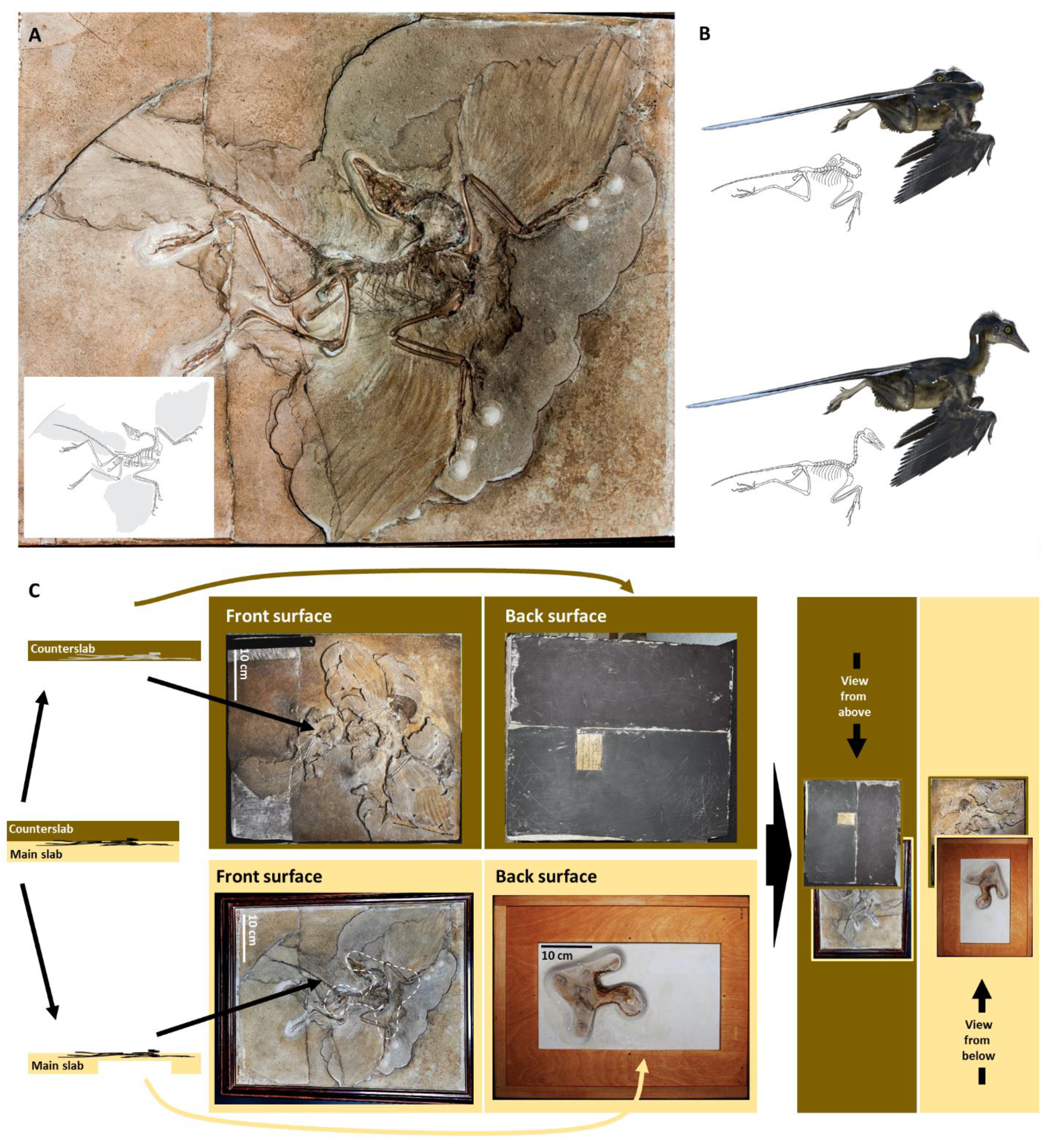
Hypothesized nesting posture of the fossilized Berlin specimen and required reinterpretation of the original relative position of its slabs. The Berlin specimen’s fossil main slab was turned clockwise from its usual upright display position to reflect the nesting posture hypothesis (**A**). Both forelimbs are abducted and flexed at the elbows and wrist, but slightly rotated counterclockwise. Postmortem overextension of the neck would have caused the thoracic vertebral column to detach from the thoracic grid. The initial position of the neck is uncertain as represented in 3D computer models of the hypothesized living nesting animal holding a retrocurved resting position (**B**, top) and an upstanding neck (**B**, bottom), each with corresponding skeletal illustrations. Reinterpretation of the original position of Berlin specimen’s fossil slabs (**C**). Contrary to the previous interpretation of the position each of the Berlin specimen’s slabs would have had on the Plattenkalk, which is based on the general finding that most Solhnofen fossils have their skeleton atttached to the underside of the upper slab following slab separation (*1*), the main slab of the Berlin specimen’s fossil is shown here in the lower position of the Solnhoffen Plattenkalk. The back surface of the counterslab is covered by a protective layer (center top) and the back surface of the main slab is also almost entirely covered by a protective mount (center bottom), except for a small central area that might provide for a ventral view of the hypothesized nest environment (right).

Despite a transversal fracture at the level of its lower third, the main slab of the Berlin specimen fossil is a single unit where most of the animal’s skeleton resides. In turn, the counterslab is composed of counterslab fragments and a slab proper.

The animal’s body is fully represented in the main slab and in the counterslab fragments (Figure 2; *1, 10*). The surface of the latter shows the ventral side of the specimen’s wings and tail feathers, which they fully encase, in addition to a few ribs. Following detachment from the main slab, these counterslab fragments fell apart from the slab proper and were believed to have been glued back to a non-original Solnhofen slab (*1*).

**Figure 2.**
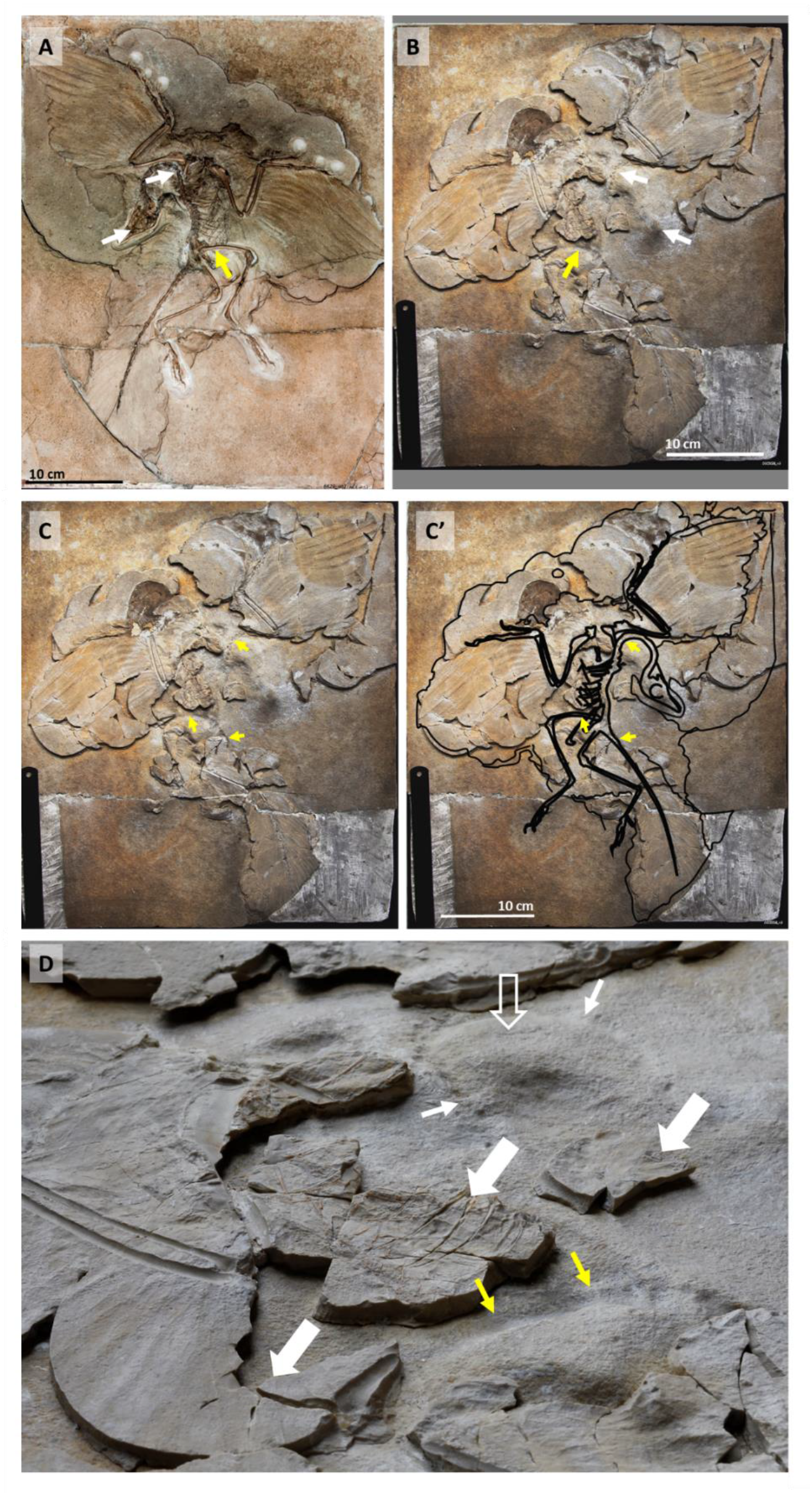
The slab proper of the counterslab is original. Arrows point at matching 3D correspondences between the main slab (**A**) and the slab proper of the counterslab (**B**). A projection of the Berlin specimen skeleton (**C**) onto the counterslab is shown on the right (**C’**) to denote matching impressions of the hindlimbs, retrocurved neck and tail in the slab proper of the counterslab (yellow arrows). 3D impressions of the Berlin specimen skeleton on the slab proper of the counterslab (**D**) demonstrate its originality. Two thin white arrows indicate an impression of the retrocurved neck onto the slab proper of the counterslab, one open arrow points at a protruding area that matches a depression in the retrocurved neck area of the main slab, and two yellow arrows point at the impression of the right femur onto the slab proper of the counterslab. Additionally, three large full white arrows point at counterslab fragments that constitute a separate layer atop the slab proper of the counterslab.

However, contrary to past opinion, reanalysis shows matching 3D impressions, a matching fracture and even matching stains between the slab proper of the counterslab and the main slab, indicating that it is original and represents a sediment layer immediately adjacent to the animal (Figure 2 and Movie 1).

The authenticity of the slab proper of the counterslab implies availability for study of the entire animal fully encased in its original fossil bed. In order to try to determine the total number of eggs, egg remnants and egg vestiges in the fossil and whether these could have belonged to the animal’s clutch, fossil areas were delineated in each slab in relation to matching areas in the opposing slab to represent different potential nest spaces, each in a particular relation to the body of the animal, as follows (Figure 3 and Movie 1):

- **Area 1** – contributed by the main slab, corresponds to the torso of the animal and structures ventral to it. Eggs and egg remnants found in this location would have been underneath the torso of the animal and therefore would be more likely to have been laid or brooded if no significant displacement of structures occurred;
- **Area D1** – represented by counterslab fragments dorsal to Area 1. It includes the dorsal surface of the Berlin specimen’s torso and sediment lying immediately over it. Eggs found in this location (over the animal’s torso) would have been probably dislodged from other locations before fossilization occurred;
- **Area 2** – is directly anterior to the body of the animal, sided by the anterior margin of its folded wings in both slabs and delimited anteriorly by the main slab depression that spans from wing to wing and matching counterslab fragments. This area was given special consideration for including distinctly dark brown areas investigated for the possibility of representing exceptional fossil preservation material;
- **Area 3** – contributed by both slabs like Area 2, includes the specimen’s wing feathers, body filaments and tail feathers and the sediment regions ventral to these. It was considered for investigation because filaments and feathers might delimit nest components;
- **Area 4** – corresponds to the main slab external to the animal’s body outline. Eggs and egg remnants in this location would be less likely to have belonged to the specimen’s clutch if displacement was not significant, and;
- **Area 5** – corresponds to the counterslab slab proper, which did not belong to the fossil’s nesting bed (*1, 10*). Eggs, egg remnants and egg vestiges found in this location would not have belonged to the Berlin specimen’s nest.

**Figure 3.**
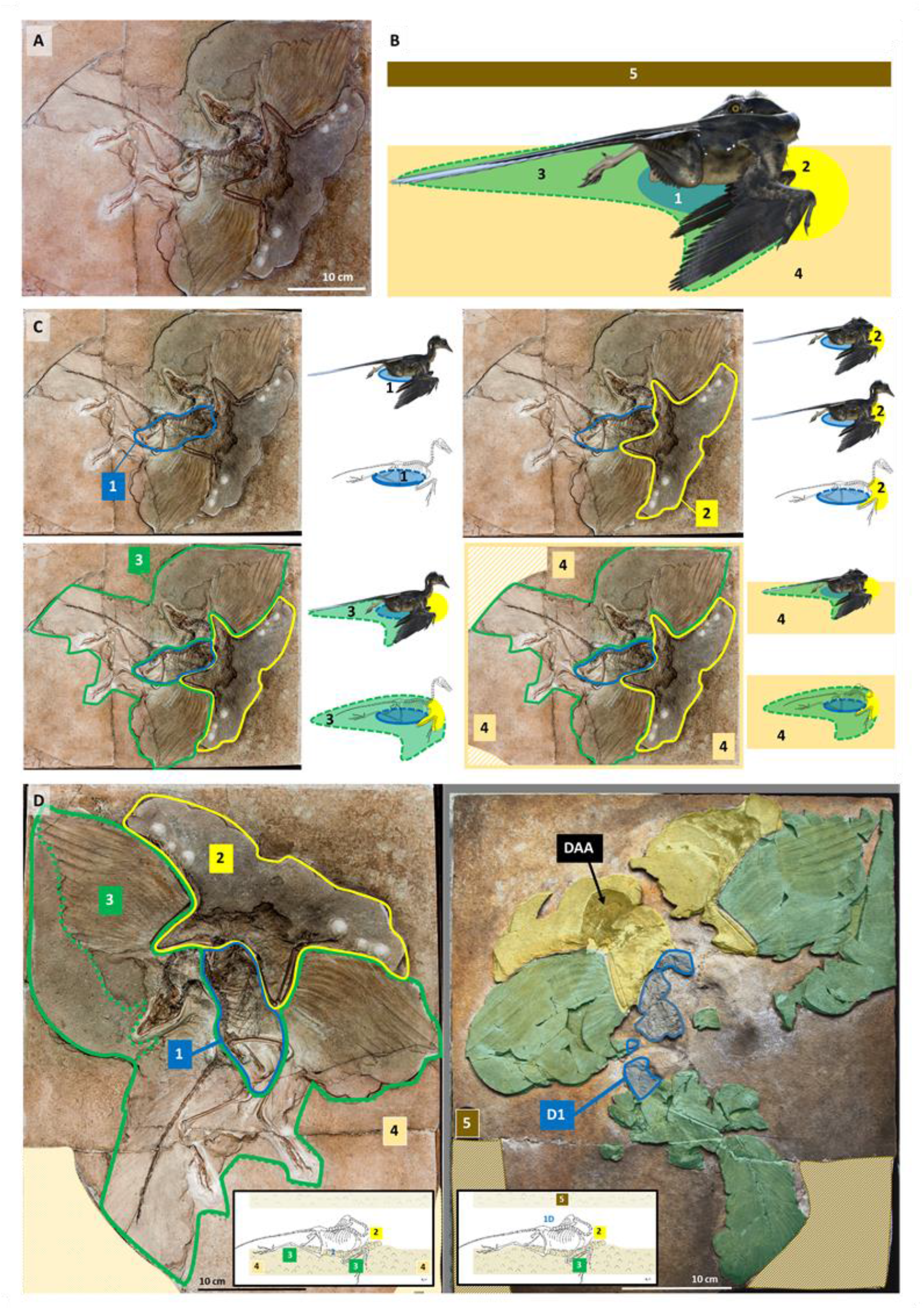
Encasement of the entire Berlin specimen in its fossil bed and definition of study areas for investigation of the nesting hypothesis. View of the Berlin specimen to reflect the nesting hypothesis (**A**) and computer model of the hypothesized nesting animal (**B**) with representation of 4 areas in the main slab and a fifth area (area 5) in the counterslab. **C**: Representation in the front surface of the main slab of the areas defined for this study of Berlin specimen’s fossil. The outlines of Areas 1, 2, 3 and 4 on the front surface of the main slab are shown, together with their hypothesized relation to the nesting animal (shown on the right of each figure section) if minimal displacement before fossilization ocurred. The computer model of the nesting *Archaeopteryx* is shown with upright or retrocurved resting neck to reflect both possibilities. Non-original areas of the main slab not considered in this study are shown filled with diagonal lines (10). The correspondance between main slab and counterslab counterparts is shown in **D**. Areas 1 and 4 reside in the main slab only. Areas 2 (in yellow) and 3 (in green) are contributed by both slabs. Areas D1 (dorsal to the specimen’s torso) and 5 (above the specimen’s fossil bed) reside in the counterslab only (Area D1 being contributed by the counterslab fragments and Area 5 by the slab proper). Non-original areas of each slab are shown filled in beige and were not analyzed in this study. Dark Area A (DAA) from the counterslab is indicated (see below).

In addition, eggs were categorized by type of evidence from Groups I – IV as follows:

- **Group I** – includes egg-enclosed fetuses and fetal remnants directly visible on the surface of the fossil slabs or detectable via transillumination;
- **Group II** – egg-enclosed fetuses identified via 3D software analysis of CT data that are not surface exposed;
- **Group III** – eggs with radiologic definition of their outline, mostly via 3D software analysis of CT data, for which no fetal preservation was identified and which are undetectable via direct observation of the fossil slabs, and;

**Group IV** – other eggs, egg remnants and egg vestiges, including directly visible egg outlines, concave or convex egg fragments, 2D egg impressions, heavily compressed eggs and 3D egg impressions that conform to the size and shape of the egg outlines observed in Groups I – III.

### Materials used in the study of the Berlin specimen

The Berlin specimen fossil is kept at Naturkundemuseum Berlin, Germany. The main slab (MB.Av.101) was observed while in its exhibition mount, which allows for vertical rotation, in the dedicated compartment of the Naturkundemuseum Berlin where it is on display. Except for the first two observations, all others were made on Mondays, when the museum is closed to the general public and the main slab could be accessed in closer proximity. Its front surface was observed on 9 occasions starting September 2014. The counterslab was observed in the collection room of the same museum in 14 different occasions starting September 2015.

The first two observations and captures of photographs of the front surface of the main slab, in September 2014 (using a handheld iPhone 4 camera and a NIKON D5200 camera with a AF-S DX NIKKOR 18-105mm f/3.5-5.6 GD VR lens without a tripod) and July 2015 (using the same NIKON D5200 camera and a Canon 5D Mark II camera associated with either a Canon EF 100 mm f/2.8 USM Macro or a Canon EF 24-105mm f/4 L IS USM lens with a tripod) were done while the fossil was still behind its bullet proof exhibition glass. On 21 September 2015 and all subsequent occasions the fossil was observed and photographed with a tripod after the protecting glass was removed from the fossil’s display case. Various angles of exposure and light conditions were used.

The Nikon and Canon cameras were used as described above, but the Canon camera was at times changed to a Canon EOS 5D Mark II, and upgraded to a Canon EOS 5D Mark IV associated with either a Canon EW-88D 16-35mm ULTRASONIC lens, a Canon Macro Lens EF 100mm 1:2.8 L IS USM or a Canon Macro Lens EF 180 mm 1:3.5 L ULTRASONIC in the 2017 sessions. The 180mm Macro had already been used with the previous Canon equipment.

Using a tripod, pictures in different angles and light conditions were obtained of the counterslab using the following cameras: (1) a NIKON D5200, (2) a NIKON D5500, (3) a Canon 5D Mark II, (4) a Canon EOS 5D Mark II or (5) a Canon EOS 5D Mark IV. Cameras 1 and 2 were associated with an AF-S DX NIKKOR 18-105mm f/3.5-5.6 GD VR lens and cameras 3-5 were associated with either a Canon EF 24-105mm f/4 L IS USM lens, a Canon EW-88D 16-35mm ULTRASONIC lens, a Canon Macro Lens EF 100 mm f/2.8 USM, a Canon Macro Lens EF 100mm 1:2.8 L IS USM or a Canon Macro Lens EF 180 mm 1:3,5 L ULTRASONIC.

Transillumination, a technique that consists in placing a strong light source in the front or back surface of the slab followed by observation, under macrophotography or stereomicroscopy, on the opposite side of the slab to that of the light source, was carried out in the Berlin specimen fossil by using a 150W or a 400W, 8850 lm, ELRO® halogen construction light. This technique had not been reported in this fossil previously and its application is made possible due to extensive preparation work done by other researchers to expose anatomical details of the wings.

Stereomicroscopy of the main slab and counterslab were done under a Nikon SMZ1270 stereomicroscope associated with a Photonic Optics LED control unit as a light source and mounted on a Motic horizontally articulated arm. The stereomicroscope was attached to a Nikon D5500 digital camera, which captured JPEG and RAW files under manual setting using a 20 second delay for picture shooting. Stereomicroscopic observation of the main slab took place on 4 occasions between April 2016 and October 2017. A preliminary observation of the counterslab under a stereomicroscope was done using a Zeiss stereomicroscope attached to a digital camera. All subsequent studies of that slab were done under a Nikon SMZ1270 stereomicroscope as described above.

Image processing was as follows: Photographs with the Canon and Nikon cameras were taken in RAW mode and later analyzed and corrected in Camera Raw Plug-in using Adobe Photoshop^®^. Adjustments were made in the exposure and contrast levels in order to get the best shadows and highlights and saved into JPEG format. All photographs were further studied and used in this manuscript in the latter format after being copied into Microsoft PowerPoint and labelled in their lower right corner for identification purposes.

A CT scan of the main slab had been obtained in 2011, previous to this study. A counterslab CT scan was obtained in 2016. Both scans were performed by Thomas Hildebrandt and Guido Fritsch from the Institute of Zoo and Wildlife Research, Berlin, Germany, using a Toshiba Aquilion CT Scanner at 0.5 mm resolution, 135 kV and 300 mA, and the results were made available for this study in the form of digital files in DICOM format.

CT scan data was explored with 3D Slicer^®^ version 4.6.2 r25516, an open source software program developed by the National Alliance for Medical Image Computing (NA-MIC). An ASUSTek^®^ (GL502VS model, with Intel^®^ Core i7-6700HQ, CPU@2.60 GHz, 32.0 Gb RAM, GPU NVIDIA Geforce GTX 1070 8 GB DDR5, and 64-bit Operating System on Windows^®^ 10) computer was used for 3D Slicer^®^ exploration regarding the data reported here. Different presets of volume rendering viewing were used.

Captures of 3D Slicer^®^ images were obtained and worked in Microsoft PowerPoint for alignment with the corresponding areas photographed as described above. The following references were used for alignments: (1) bone structures; (2) crosshairs and slice intersections provided by 3D Slicer^®^ in coronal, sagittal and transversal views obtained in different orientations, and (3) measurements obtained during direct observation and photography of the fossil.

The image of the Berlin *Archaeoptery*x with surface representation of each egg, egg remnant and egg vestige was produced in Microsoft PowerPoint using scaled models of a complete egg and detailed photographs for guidance.

Illustrations of *Archaeopteryx* at its hypothetical nest in dorsal and lateral view were produced using Adobe Illustrator CS5 software. SketchUp 8^®^ software program was used to produce a 3D model of a single *Archaeopteryx* egg. Next, the same program was used to produce a model of how all discovered eggs are positioned relative to each other in the fossil and in relation to the 2D plane of the dorsal surface of the Berlin specimen.

A 3D model of the nesting Berlin *Archaeopteryx* was produced using the ArchaeopteryxDR – Extended License 3D model developed by the 3D model author Raul Lunia and licensed from DAZ Studio 4.9 software.

### Teylers Archaeopteryx

The Teylers specimen fossil (collection item with references 6928 and 6929 for its two slabs, respectively main or bigger and counter or smaller slabs) is kept at the Teylers Museum, in Haarlem, Netherlands. It was observed outside its display case on 23, 24, 30 and 31 January 2017, 20 and 22 March 2017, 28 August 2017, 18 September 2017, and 16 and 17 April 2018.

Both slabs were photographed using a Canon EOS 5D Mark IV and either a Canon Macro Lens EF 100 mm 1:2.8 L IS USM or a Canon Macro Lens EF 180 mm 1:3.5 L ULTRASONIC. Approximately 1 000 photographs were taken. Different angles and light conditions were used with the help of a tripod to change the angle of exposure of the freely movable fossil slabs. Careful examination and serial pictures of the side surfaces of both slabs were also obtained.

Stereomicroscopy was as described for the Berlin specimen, and about 1 600 stereomicroscope photographs of the Teylers specimen were obtained.

Image processing was as described for the Berlin specimen.

Candidate egg-like structures were named using the acronyms TSS (Teylers Small Slab) when identified in the smaller slab (6929 or counterslab), TBS (Teylers Big Slab) when found in the bigger slab (6928 or main slab), TS/BS to designate the counterpart located in the smaller slab of a structure split among both slabs that had initially been identified in the bigger slab, and TB/SS in the opposite situation.

Photographs were explored using as search tool the computer generated line models of MSB2, which are identical to the outline of TBS1 from Teylers, and also by using the line model of TSS1 (TBS1 and TSS1 were the most noticeable egg-like structures upon initial observation of the Teylers specimen and were immediately identified).

Areas of egg-like structures identified via photography were further studied using macro lenses and the stereomicroscope, and detailed 3D software analysis of CT scan data was performed to further investigate their existence (see below).

The Teylers specimen’s fossil was subjected to a medical quality CT scan at 0,5 mm resolution using the same parameters described above for the Berlin specimen’s fossil. In addition, it was also subjected to a micro CT scan on a Zeiss Xradia 520 versa CT scanner at the Naturalis Museum, in Leiden, Netherlands, at resolutions ranging from 35 to 46 micrometer. Different areas of each slab, each measuring approximately 4.7 x 4.7 cm, were individually scanned at a time. Scanned data from each slab was stitched together using Vertical Stitch – 11.1.5707.17179 from Zeiss. 160KV with 10W were used to scan both slabs. Amperage was 50 mA in the 6928 (bigger or main) slab and 63 mA in the 6929 (smaller or counter) slab. Stitched data from the Teylers CT scans at nearly 45 μm resolution was either used in DICOM format or saved in TIFF files and explored using 3D Slicer^®^ as described for the Berlin specimen’s fossil. This software was used to screen for 3D structures that would correspond to the size and shape defined by the outline of putative eggs TSS1 and TBS1. Bi-dimensional CT scan images were also screened using the same approach.

### Maxberg *Archaeopteryx* X-rays

Maxberg specimen X-rays by Wilhelm Stuermer from Wellnhofer’s “*Archaeopteryx* The Icon of Evolution”, 2009, were obtained in digital TIFF format from Verlag Dr. Friedrich Pfeil.

### Thermopolis Archaeopteryx

This fossil’s main slab was observed at the Dinosaur Center, in Thermopolis, Wyoming, in its display case on 8 March 2016, and after it was removed from its display case on 9 March 2016 and from 28 February until 2 March 2017. Photography, stereomicroscopy and image processing were conducted as described for the Berlin specimen’s fossil. Over 500 macrophotographs and over 600 stereomicroscope photographs were obtained. Photographs of this specimen captured under UV light were kindly provided by Dr. Burkhard Pohl.

### Illustrations of the skeletons of *Archaeopteryx* specimens

The illustrations of the skeletons of the Berlin, London, Eichstatt, Teylers, Solnhofen, Maxberg, Thermopolis and Munich specimens were modified from those by Wellnhofer, P. (Ed.) *Archaeopteryx* The Icon of Evolution. Ed Verlag Dr. Friedrich Pfeil (2009). The illustrations of the 11^th^ and 12^th^ specimens were produced using Adobe Illustrator CS5 software.

### Isolated *Archaeopteryx* feather fossil

The counterslab of the isolated *Archaeopteryx* feather fossil (MB.Av.100) kept in the Collection Room of the Naturkundemuseum Berlin, Germany, was studied in that room in multiple occasions under macrophotography and stereomicroscopy using the same materials and methods described above for the Berlin specimen fossil counterslab during some of the visits to the Collection Room of the said museum described above to study the latter slab.

## Results

### Skeletal findings in Berlin specimen’s torso and their relation with a ventral egg mass that lies ventral to its thorax

The lumbar-thoracic transition of the Berlin specimen’s vertebral column is curved. This curved vertebral segment is immediately caudal to a straight thoracic segment that detached from the costal grid (*10*), and overlays two globular prominences in the right hemithorax of the specimen. These bulbous protuberances pierced through the thoracic wall of the animal, causing sharp curvatures in its ribs (Figure 4). These protuberances are at the center of a mass that lies ventral to the Berlin specimen’s thorax, which is composed of delimited globular radiodense units with various levels of preservation (Figure 5).

**Figure 4.**
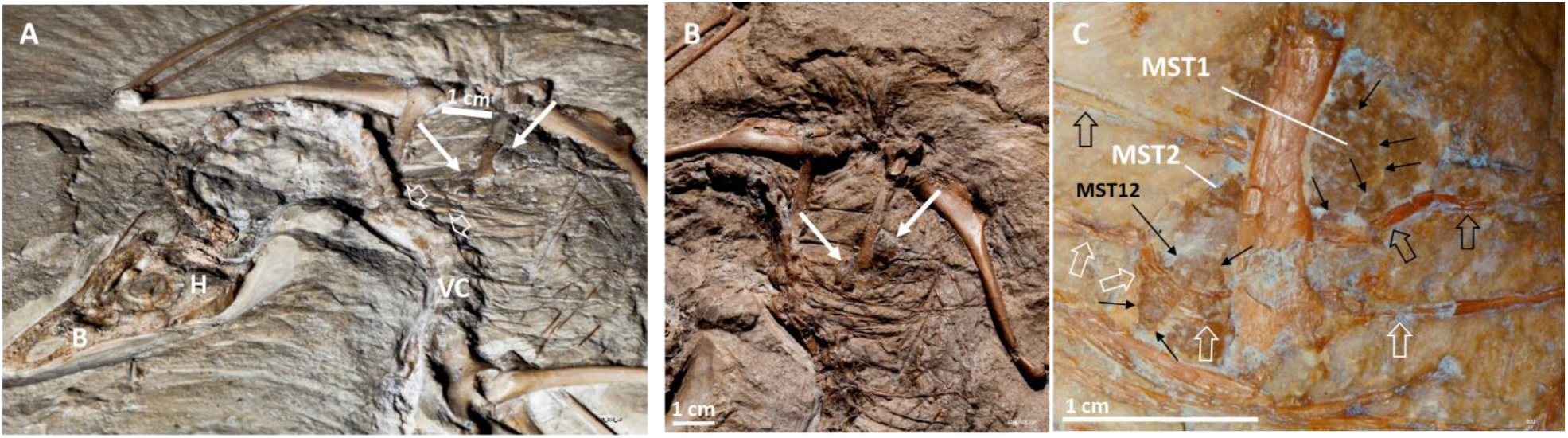
A curvature in the Berlin specimen’s vertebral column and two prominences in its thoracic wall. A caudo-rostral view of the Berlin specimen fossil (**A**) puts into evidence both a curvature in the vertebral column (open arrows) that is caudal to the detachment of this skeletal structure from the thoracic grid, and two prominences in the thoracic wall (thin arrows), shown in front view in **B** and magnified in **C**. The two thoracic wall prominences sharply distort overlying ribs (open white and open black arrows identify two different ribs). Surface exposure of the MST1, MST2 and MST12 structures is indicated. These and other irregularities noted on the surface of both thoracic wall prominences, indicated by smaller thin black arrows in C were investigated and are described below. B – Beak; H – Head; VC – Vertebral column.

**Figure 5.**
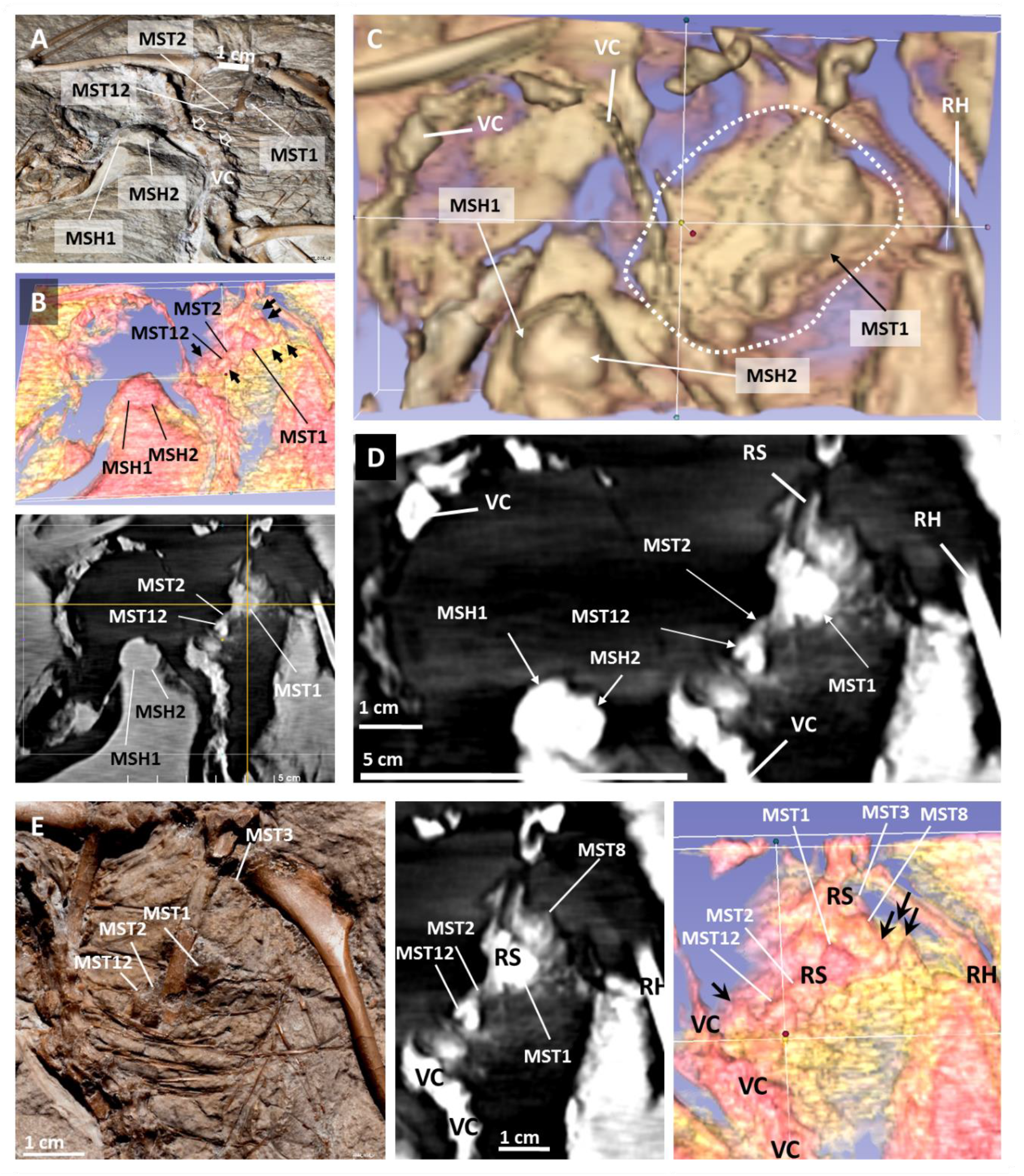
The Berlin specimen lies atop a mass of globular structures that is ventral to its thorax. A curvature in the vertebral column of the Berlin specimen precedes its detachment from the thorax (**A**, open arrows) and lies atop globular units better perceived via 3D software analysis of CT scan data (**B**, top), and conventional CT scan images (B, below), which integrate a radiodense mass observable under 3D software analysis of CT scan data (**C**) and in coronal 2D CT scan sections (**D**). This mass is composed of delimited globular units shown to correlate between macrophotography (**E**, left), 2D CT scan images (E, center) and 3D software analysis of CT scan data (E, right). MST1, MST2, MST3, MST8, MST12, MSH1 and MSH2 indicated here are some of these globular structures that will be described below. RH – Right Humerus; RS – Right Scapula; VC – Vertebral column.

Some of these globular structures are apparent on the fossil surface under normal lighting conditions or transillumination, whereas others are only detectable under CT scan analysis (see below). Despite their different 3D shapes and appearing to have been distorted by neighboring structures, including the adjacent skeleton, all measure 1.0 to 1.4 cm in their longer axis, averaging 1.3 cm in length. Some show an outline suggestive of egg-enclosed fetal preservation. For example, MST11, MST12, MST38 and MST40 are adjacent to each other and observable under macrophotography in the right hemithorax of the Berlin specimen in the front surface of the main slab (Figure 6). In turn, MSB5 and MSH27 are observable in the specimen’s left scapula area under transillumination (Figure 7). Examples of 3D fetal morphology preservation detectable under CT scan analysis will be described below.

**Figure 6.**
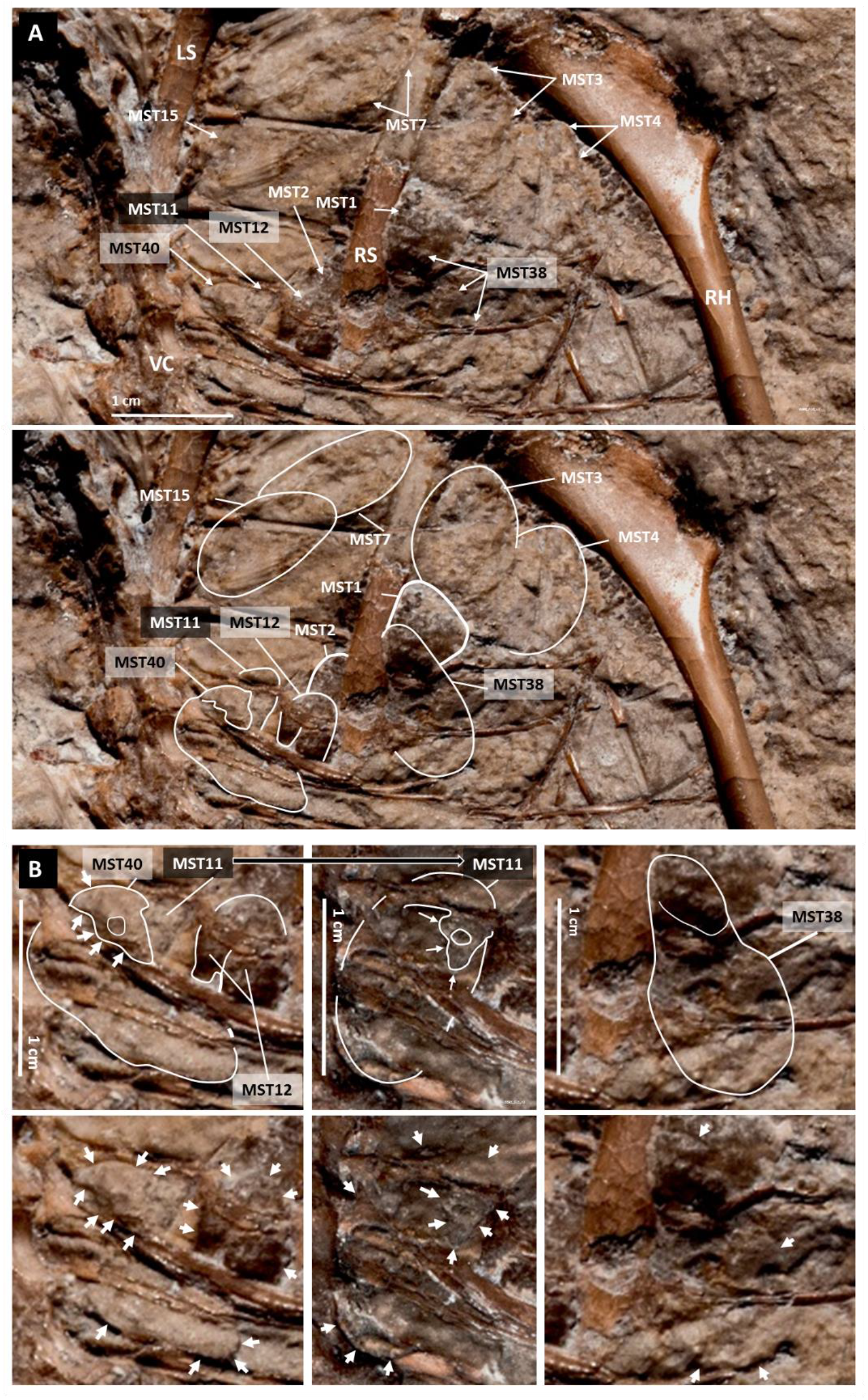
Outline of globular structures on the surface of the thorax of the Berlin specimen, some of which seem to encase a fetus with surface morphology preservation. Macrophotography of the surface of the thorax (**A**) allows to outline several structures, some of which are also detectable via CT scan analysis (see Figure 8 below and data not shown), both within and around the protuberances that pierced through the thoracic wall and distorted adjacent ribs. Some of these structures seem to encase a fetus with surface morphology preservation (**B**).

**Figure 7.**
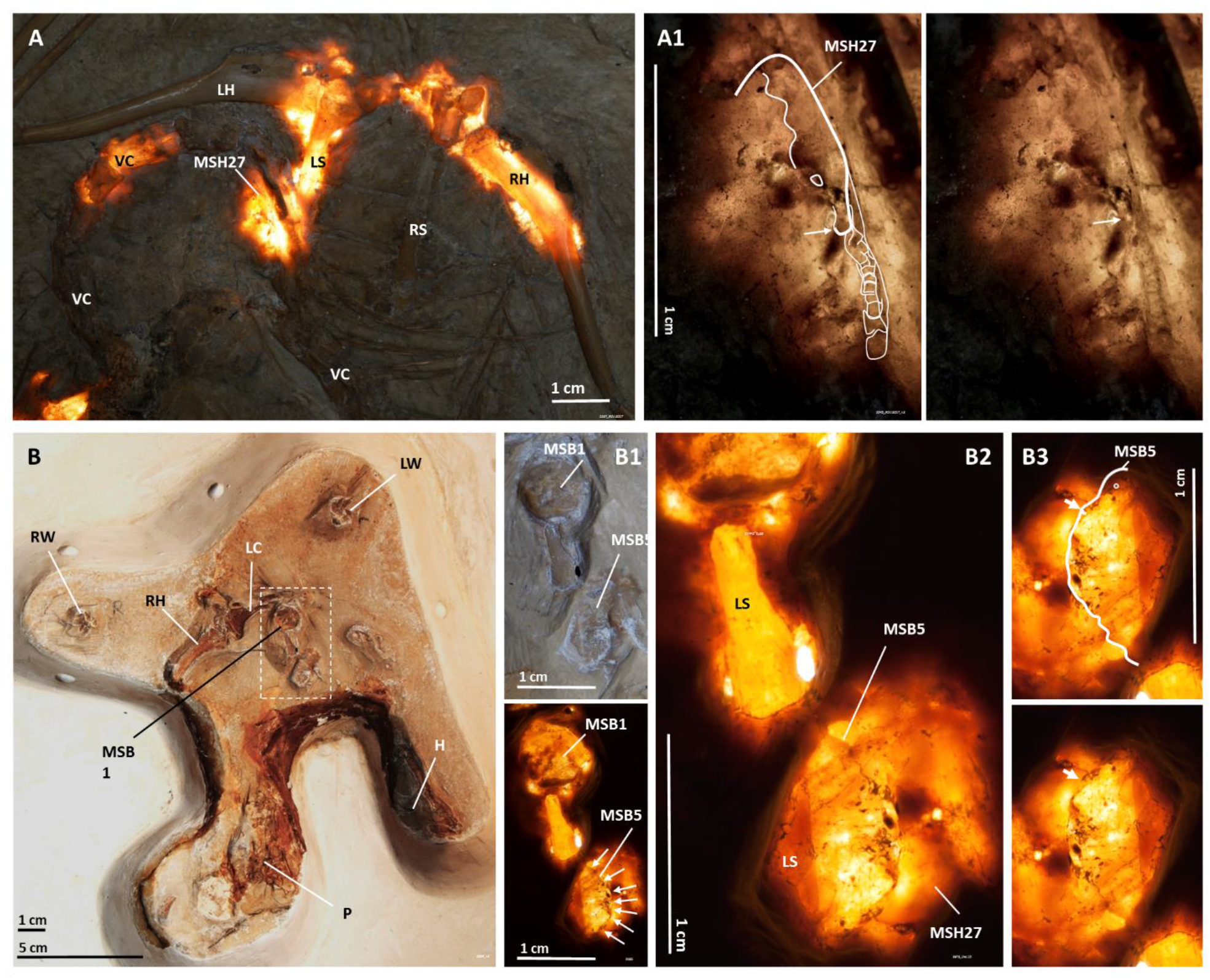
Transluminescent areas of the main slab of the Berlin specimen conceded egg discoveries. MSH27 is a globular structure that seems to encase a fetus with surface morphology observable via transillumination on the upper body area of the front surface of the main slab (**A** and **A1**). In turn, MSB5 is detectable on the back (ventral) surface of the main slab (**B**, **B1**, **B2** and **B3**). The dashed rectangle in B is seen under natural light in B1-top and transillumination in B1-bottom. MSH27 and MSB5 are adjacent to each other in the left scapula area (B2 and data not shown). H – Head; LC – Left Coracoid; LH – Left Humerus; LS – Left Scapula; LW – Left Wrist; P – Pelvic area; RH – Right Humerus; RS – Right Scapula; RW – Right Wrist; VC – Vertebral Column.

A 3D computer model of the thoracic mass based on a unitary structure slightly flattened and measuring 1.3 x 1.0 x 0.8 cm correlates with observations of the thorax under macrophotography, transillumination, stereomicroscopy and CT scan analysis (Figure 8).

**Figure 8.**
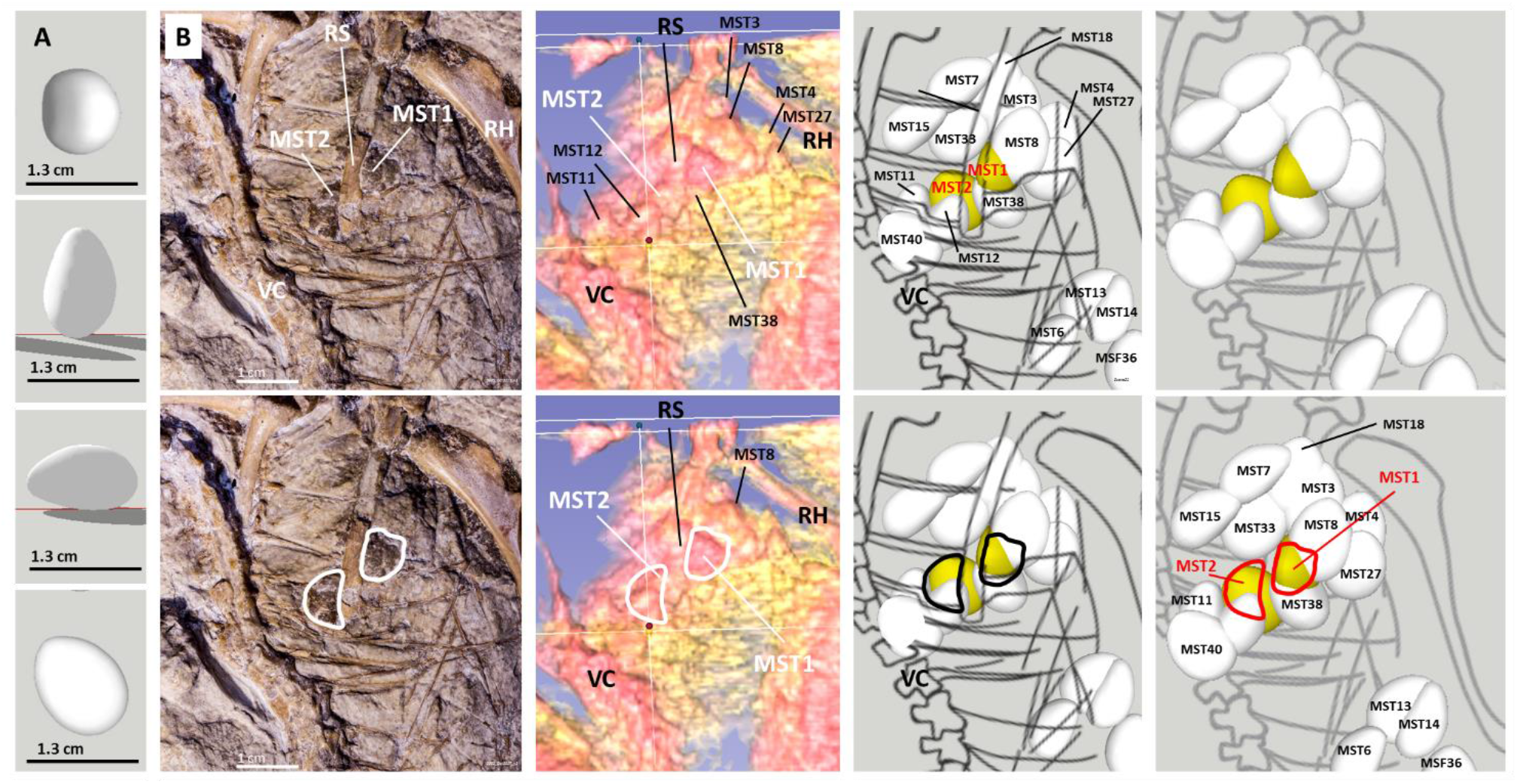
Prominences in the Berlin specimen’s thoracic wall produced by an egg mass. A 3D software model based on an egg measuring 1.3 x 1.0 x 0.8 cm (**A**) shows data concurrence with macrophotography (**B-left**) and 3D software analysis of CT scan data (**B-center left**). The prominences that pierced through the Berlin specimen thoracic wall are outlined in red on the dorsal surface of the Berlin specimen’s skeleton (**B-center right**) and in the projection of the skeleton onto the plane that the eggs lie against (**B-right**). All egg remnants are represented as complete eggs in the 3D model. RH – Right humerus; RS – Right scapula; VC – Vertebral column.

### A fossilized egg-enclosed fetus with a toothed beak is ventral to Berlin specimen’s thorax

Among other egg-like structures ventral to the animal’s torso, within Area 1 of the Berlin specimen fossil (see Figure 3 in Materials and Methods), MSB5, shown above in Figure 7, displays, under stereomicroscopy with transillumination, in one of its poles, the head morphology of a beaked fetus with teeth preservation, apparently located in the upper jaw (Figure 9).

**Figure 9.**
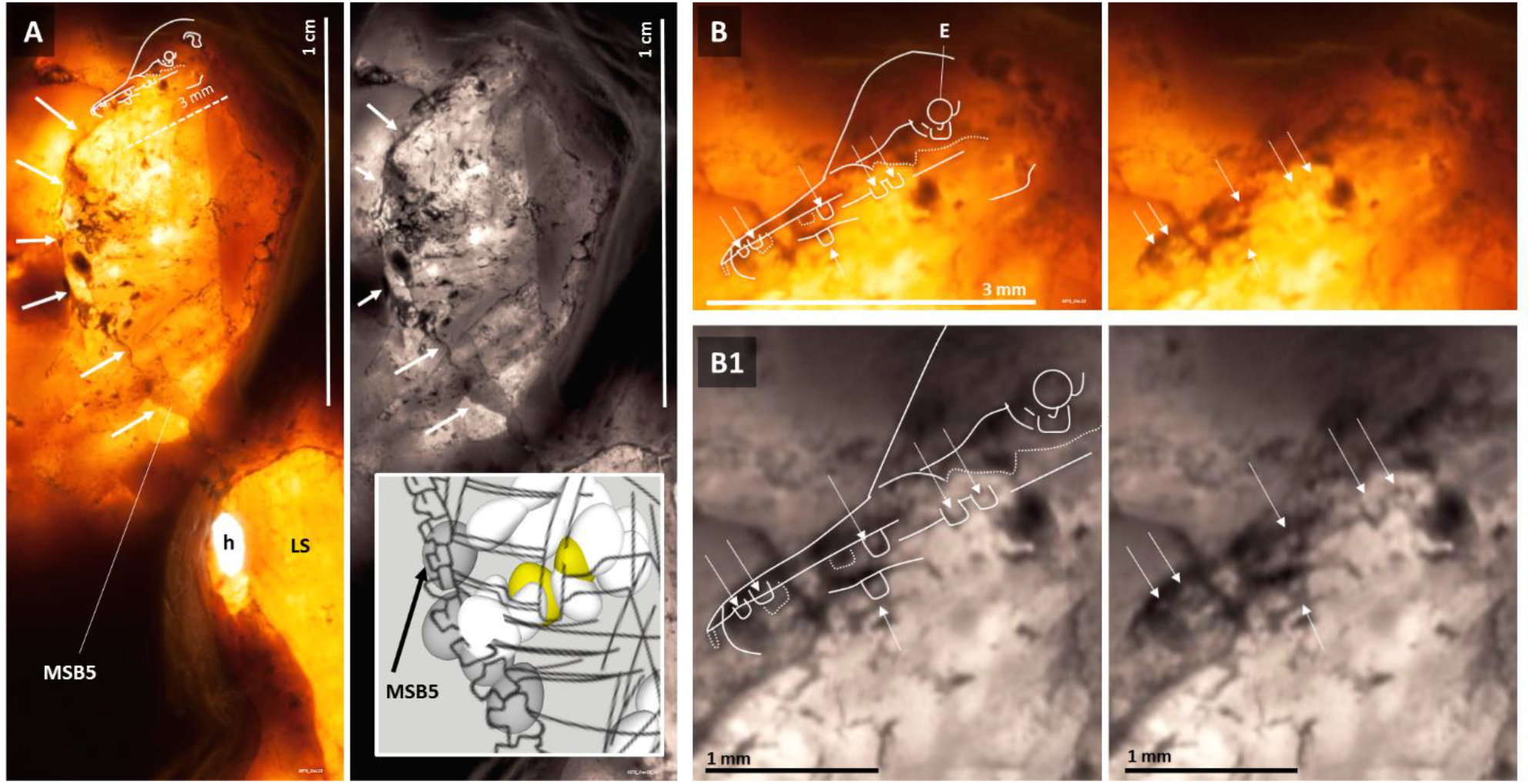
MSB5’s fetal head has a toothed beak. Contour of the head and beak of MSB5’s fetus under transillumination by putting a light source in the front and observing it from the back of the main slab of the Berlin specimen fossil. Inset: MSB5’s position in relation to the left scapula in a 3D computer model; MST1 and MST2, located in the protuberances of the right hemithorax (Figure 4) are shown in yellow for reference. Transillumination images were rotated by 180 degrees in relation to the longer axis of the fossilized adult animal. Magnified views of teeth in MSB5’s fetal head are shown in **B** and **B1**. Arrows are provided for positional guidance between B and B1. E – Eye; h – small hole between the front and back surfaces of the main slab caused by previous preparation work by other researchers; LS – Left Scapula.

### Eggs anterior to the Berlin specimen and its folded wings and their juxtaposition against the right wing

In addition to fetal morphology preservation in the main slab, it was also observed within one of two distinctly brown and radiodense areas of the counterslab suggestive of fossilized organic matter, named Dark Area A (DAA) and Dark Area B (DAB, *10*), located anteriorly to the Berlin specimen, within Area 2 (Figure 3). DAA, which is just anterior and medial to the wrist of the flexed right wing of the Berlin specimen, shows outlined eggs with fetal surface morphology preservation that can be observed under macrophotography (Figure 10).

**Figure 10.**
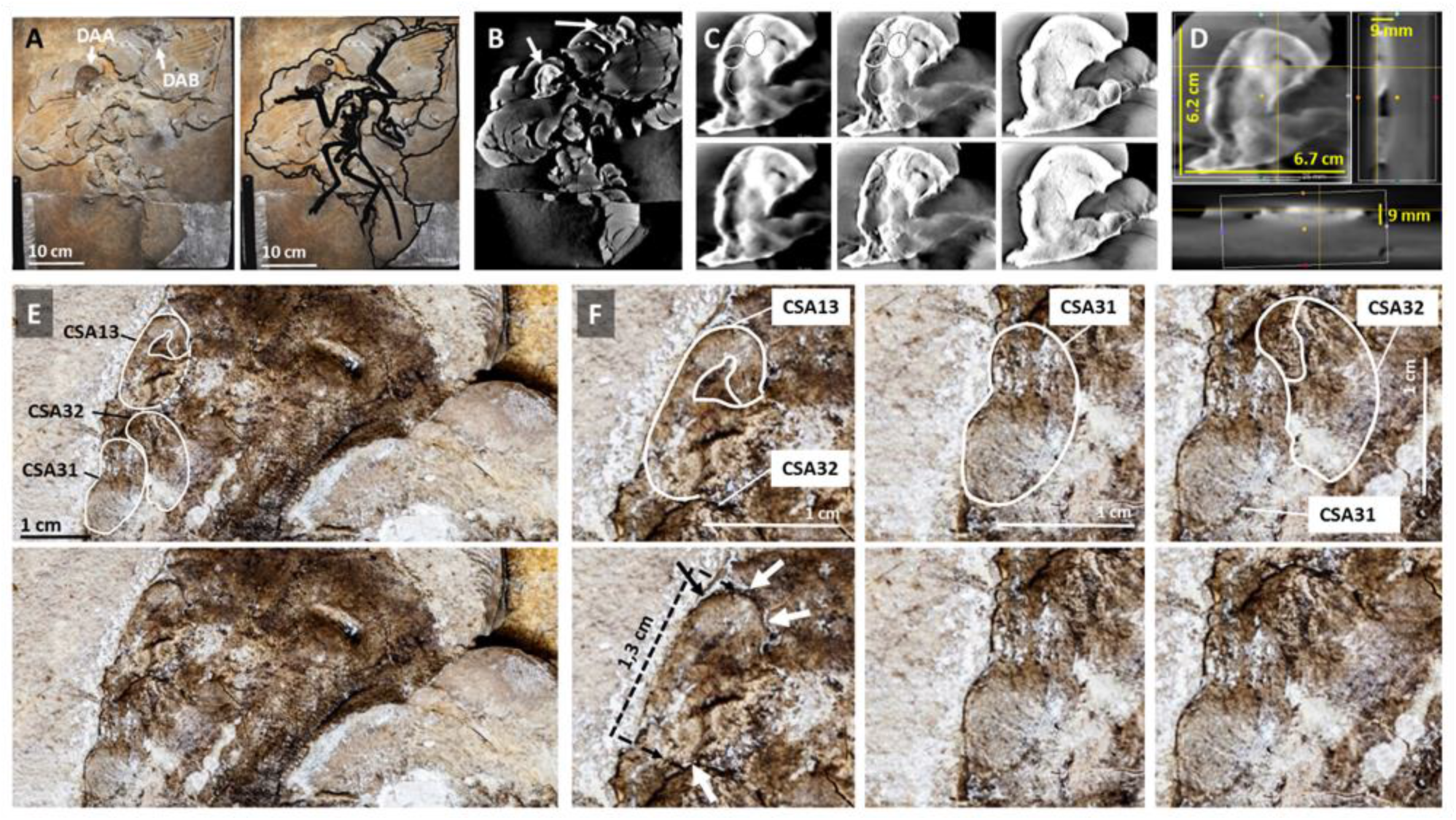
A newly described area of the counterslab with distinctly carbonaceous appearance contains distorted eggs with fetal surface morphology preservation. DAA and DAB (**A**) are radiodense under CT scanning (**B**). The former contains radiodense globular units of consistent size, approximately 1.3 cm long, outlined in different coronal CT views (**C** and **D**) and under macrophotography (**E**), some enclosing fetuses with surface morphology preservation (**F**).

Within a radiodense area that is also adjacent to the flexed right wing of the Berlin specimen, but located in the main slab, between the right humerus and the right antebrachium, are sub-vertically juxtaposed structures that cause crevices in the sediment (Figure 11). Some of these show the outline of a tucked fetal head with surface morphology preservation similar to that of DAA’s CSA13 and CSA32 – see MSF17 and MSF18 in Figure 11. Most structures from this area of the main slab are detectable only via 3D software analysis of CT scan data (data not shown).

**Figure 11.**
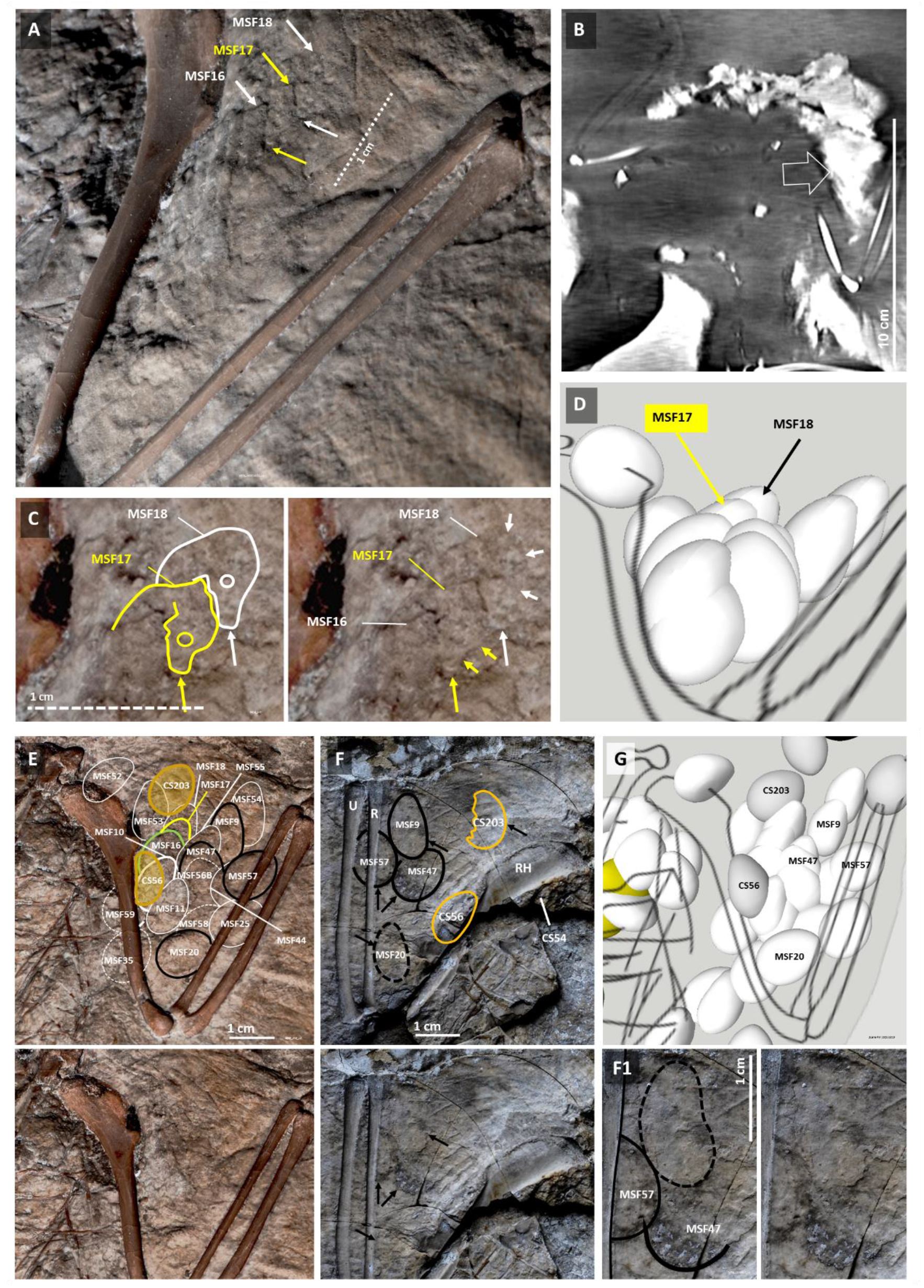
Juxtaposed fetal heads with surface morphology preservation near the right humerus of the Berlin specimen. Regularly spaced crevices in the sediment (arrows in **A**), anterior to the folded right wing of the Berlin specimen, where a mass of radiodense sediment is located (open arrow in **B**), led to the identification of two directly visible fetal beaked heads in tucked position with surface morphology preservation (arrows in **C**). A 3D model of these and other egg remnants identified in the same area via 3D software analysis of CT scan data is shown in **D** in an inclined caudo-rostral view of a computer model where are eggs are represented as complete eggs regardless of their status of preservation. The outlines of those eggs identified via 3D software analysis of CT scan data of the main slab are shown in **E** and the outlines of egg impressions detected via macrophotography and stereomicroscopy in the counterslab are shown in **F**. Outlines in black match between in E and F (main slab and counterslab, respectively). A front view of this right wing area as per the 3D computer model is shown in **G**.

When this area of the right wing anterior to the humerus and antebrachium is observed in the counterslab, several impressions of sagittal and coronal outlines of eggs can be observed via macrophotography and stereomicroscopy. Some of these match precisely, in their contour and position, the outlines of structures identified in the main slab (Figure 11F and F1).

Oval structures of similar size and shape are also found in the main slab anteriorly to the specimen’s torso (Figure 12A-B), and between the left wing humerus and antebrachium, just as seen in the right wing. However, the eggs in the left wing are mostly detectable via 3D software analysis of CT scan data (Figure 12C-C1’ and data not shown). Once identified via CT scan, the outlines of some of these structures become observable in the main slab surface and inclusively show some degree of surface fetal morphology preservation (Figure 12D). Accordingly, flattened oval remnants (CS205 and CS206) and an oval impression (CS204) are also observable in the corresponding left wing area of the counterslab (Figure 12E-E1), as modeled in Figure 12F (see Movie 1).

**Figure 12.**
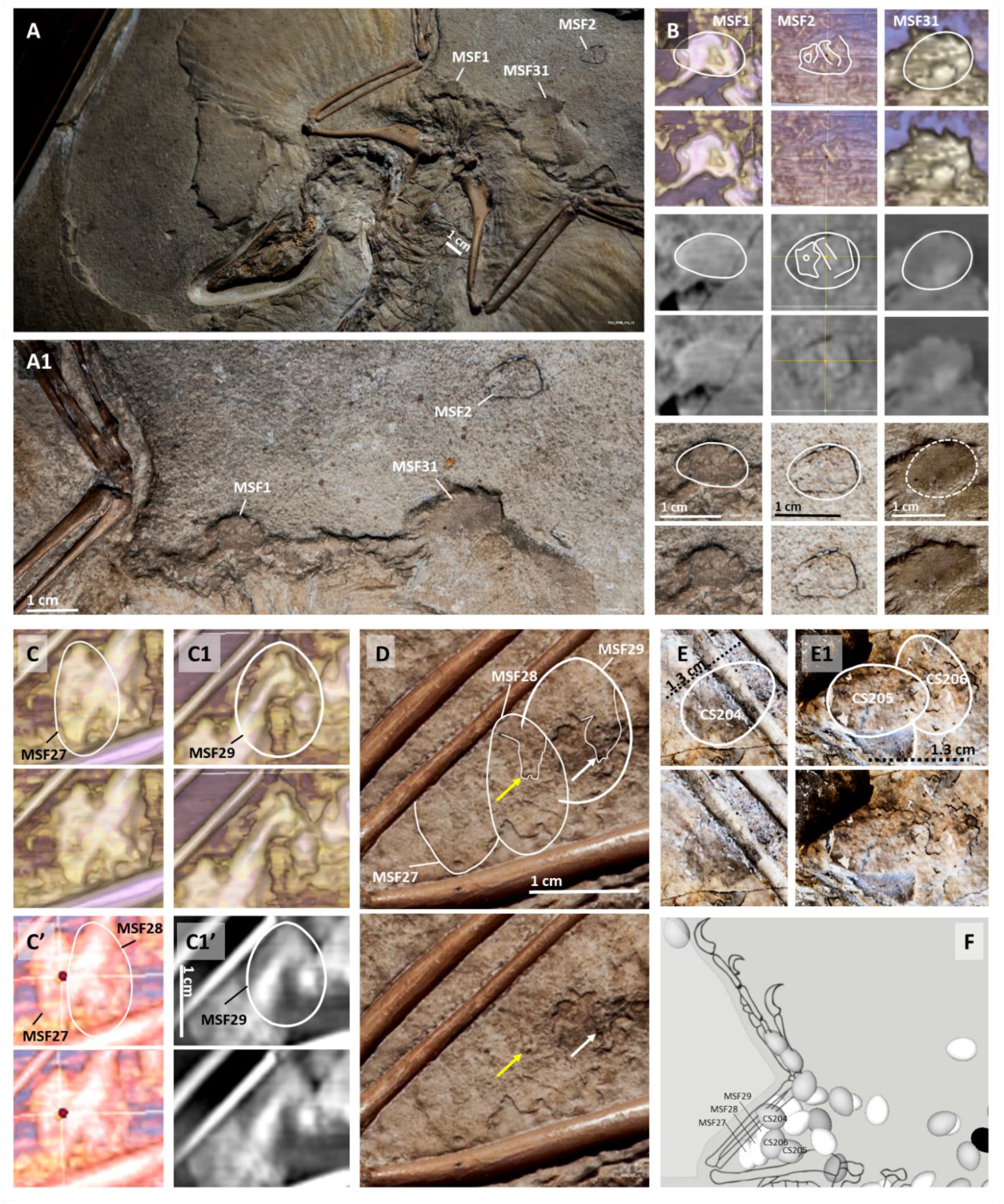
Egg remnants located anteriorly to the Berlin specimen and its left wing. MSF1, MSF2 and MSF31 located within Area 2 in the front surface of the main slab (**A** and **A1**), are shown in 3D images of CT scan data (**B**, top), 2D coronal CT sections (B, middle) and under macrophotography (B, bottom). Egg remnants detected via 3D software analysis of CT scan data in the space between the left humerus and antebrachium (MSF27 and MSF28, respectively outlined in **C** and **C’**, and MSF29 outlined in **C1**). MSF29 is also observable in 2D scan images (**C1’**). Outlines of MSF27, MSF28 and MSF29 detectable on the surface of the main slab at the location defined by CT scan data; the faint outline of the fetal heads within MSF28 and MSF29 are shown in **D**. Egg outlines in the same left wing area in the counterslab (**E** and **E1**). A 3D software model of the left wing area is shown in **F**.

Contrary to all these structures anterior to the flexed wings, only scarce egg-like impressions were observed within the perimeter of the adult animal’s wings in both slabs (Area 3 in Figure 3).

### Egg remnants in other locations around the Berlin specimen’s body

Eggs can also be found beyond the area ventral to Berlin specimen’s thorax and anteriorly to its flexed wings but in lesser numbers (see below). Examples of main slab eggs located in the retrocurved neck area of the Berlin specimen and in the posterior part of its body are respectively shown in Figures 13 and 14 and also in Movie 1 (the latter also shows the ventral surface of the main slab).

**Figure 13.**
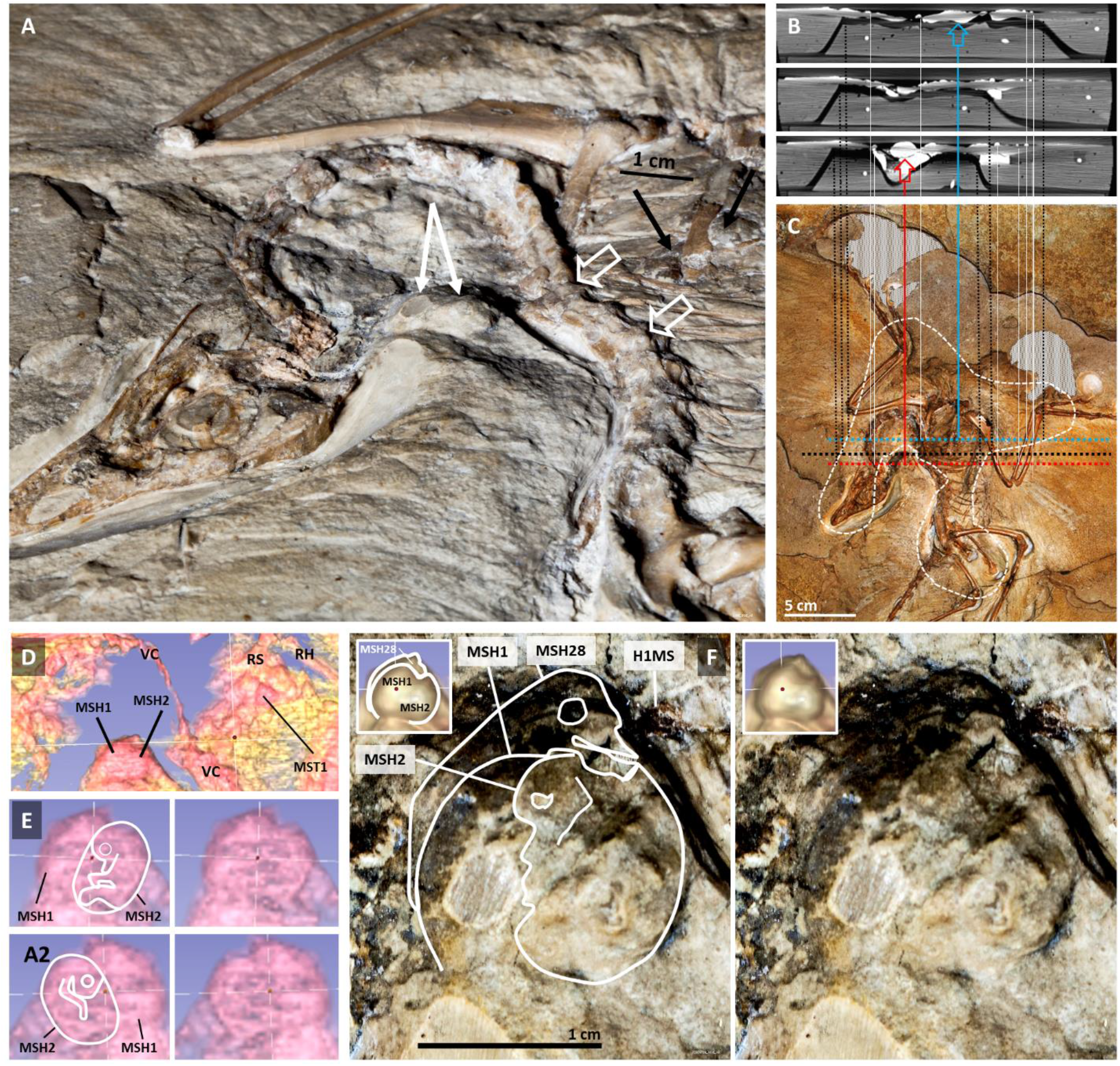
Main slab eggs in the retrocurved neck area of the Berlin specimen. An angled view of the main slab’s front surface (**A**) puts globular structures into evidence (white arrows) to the left of the vertebral column curvature (open arrows) dorsal to protrusions of the ventral egg mass (black arrows). The vertebral column rostral to said curvature detached from the thoracic grid due to postmortem neck overextension (*1*). Transversal CT scan sections (**B**) show two adjoined globules (red open arrow) at the level of the thoracic protuberance that contains MST1; the thoracic egg mass appears as a compressed and confluent radiodensity at this resolution (blue open arrow). **C**: Correspondence between 2D CT images and their main slab location. The globular structures shown in A and B are identified as MSH1 and MSH2 in 3D software analysis of CT scan data (**D**-**E**). The outline of fetal preservation within MSH2 detected via CT scan (E) matches observations under macrophotography and stereomicroscopy (**F)**. An additional remnant seems to lay over MSH1 and MSH2 (MSH28 in F).

**Figure 14.**
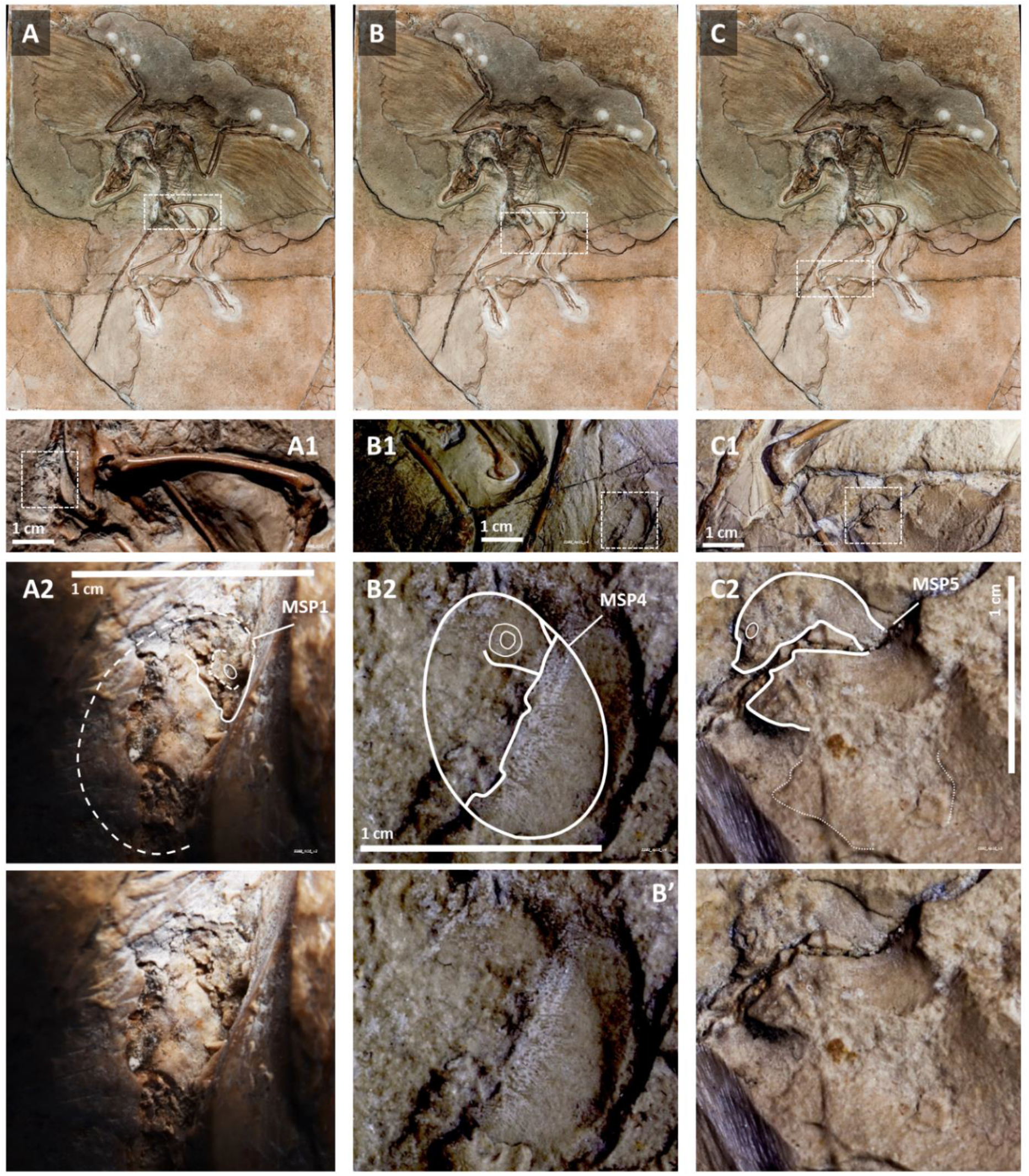
Main slab eggs in the posterior part of Berlin specimen’s body. MSP1 is an egg with an enclosed fetal remnant with surface morphology preservation located against the dorsal surface of the pelvic vertebral column of the Berlin specimen (**A-A2**). It is also detectable via 3D software analysis of CT scan data (data not shown). **MSP4 is a** compressed egg remnant located anteriorly to the right tibia on the front surface of the main slab of the Berlin specimen (inset in **B** and **B1**). The lower half of the egg surface was preserved together with an impression of its fetus in the upper half (**B2**). MSP5 is a fetal remnant in egg enclosed position exposed on the front surface of the main slab that is medial to the left foot of the Berlin specimen (**C-C2**).

Among others located in the posterior part of the Berlin specimen’s body, an egg designated CS1, with a fetal outline detected under 3D software analysis of CT scan data, was found in a counterslab fragment near the left foot of the specimen (Figure 15). Surface 3D impressions of a fetal folded wing and parallel linear structures compatible with feather remiges shafts can be seen in a position compatible with the general fetal outline derived from CT scan analysis of this egg (Figure 16).

**Figure 15.**
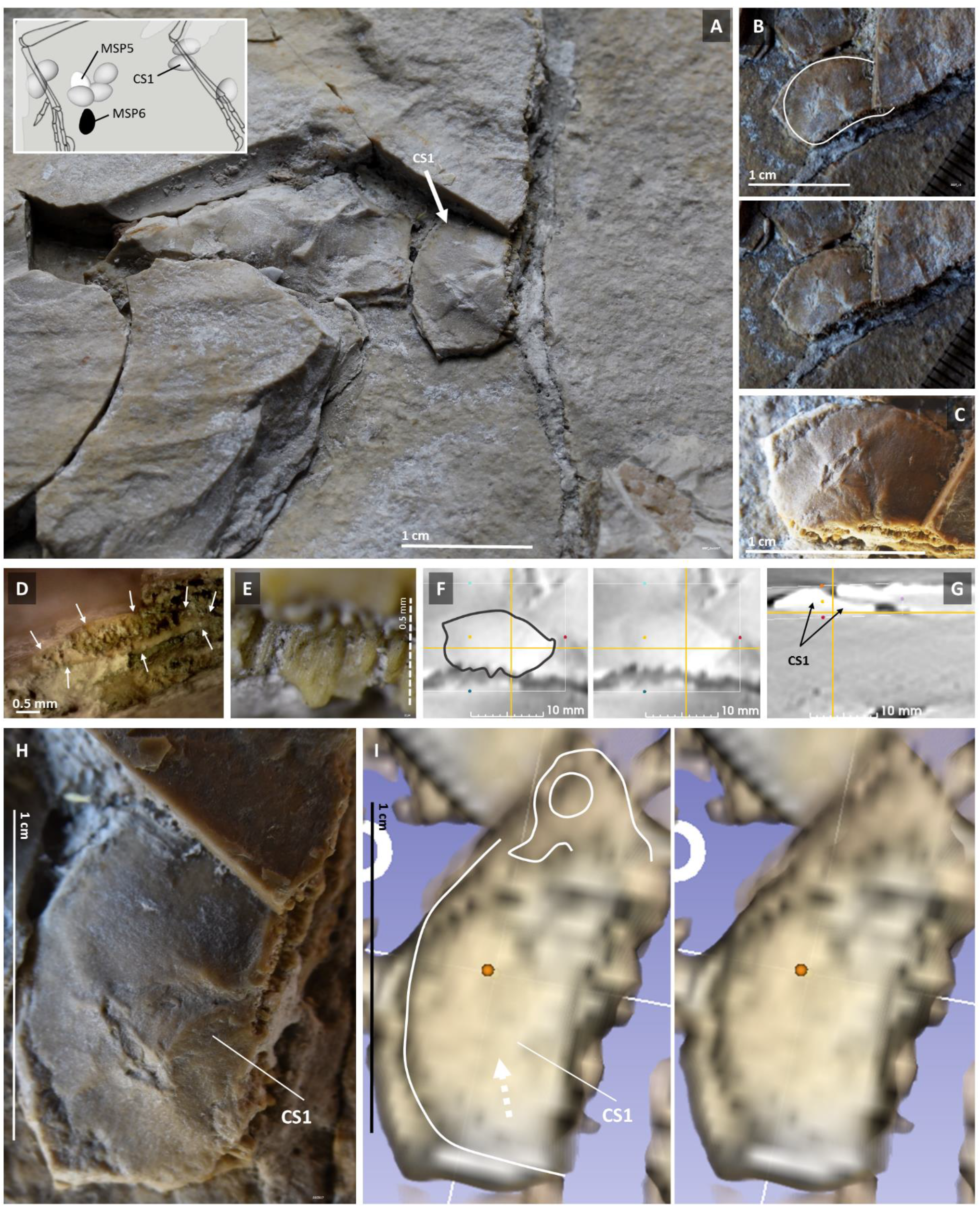
A counterslab egg in the posterior part of Berlin specimen’s body. A macrophotograph of the area of the right foot of the Berlin specimen in the counterslab (**A**; see inset) indicates the CS1 egg, outlined in **B** and shown under stereomicroscopy in **C**. It is broken just like the slab proper of the counterslab beneath it. Its broken edge is shown in **D** under a radial view and magnified in **E** in top view. The complete outline of CS1 can be appreciated under coronal (**F**) and transversal (**G**) 2D CT sections at the level where it extends underneath another counterslab fragment. An amplified view of CS1 under stereomicroscopy is provided (**H**) to correlate with the outline of its fetus detected via 3D software analysis of CT scan data (**I**).

**Figure 16.**
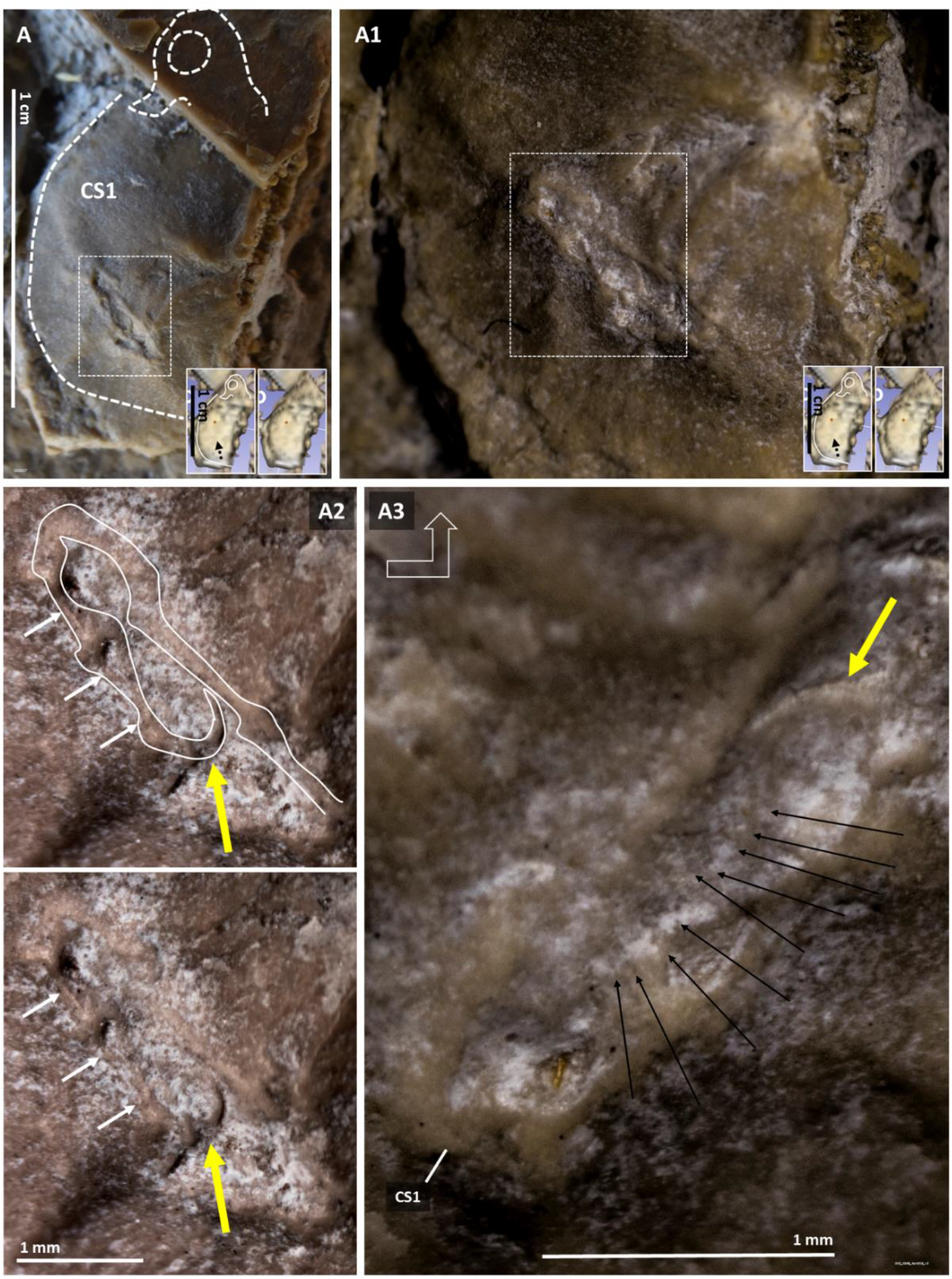
Observable 3D outline of a folded fetal wing with feather shafts at a location compatible with data obtained via CT scan. **A:** Outline of the CS1 egg and of the fetal head within it detected via 3D software analysis of CT data (inset) are shown. The dashed rectangle area was increasingly amplified in **A1**, **A2** and **A3** (the latter image was turned approximately 90 degrees counterclockwise). The yellow arrow in A2 and A3 points at the wing claw and the black arrows in A3 at linear impressions that, given their position in relation to the folded wing outlined in A2, may represent the shafts of feathers.

### Eggs lying on the dorsal surface of Berlin specimen’s torso

Two eggs with similar characteristics reside over the specimeńs right hemi-thorax, one, with fetal surface morphology preservation, at the surface of one counterslab fragment (CS54 in Figure 17) and the other egg, with fetal morphology detected via CT scan analysis, within another of such fragments (CS83 in Figure 17B and data not shown). The counterslab fragments that contain these eggs present rib fragments of the adult animal on their surface (Figure 17A and data not shown).

**Figure 17.**
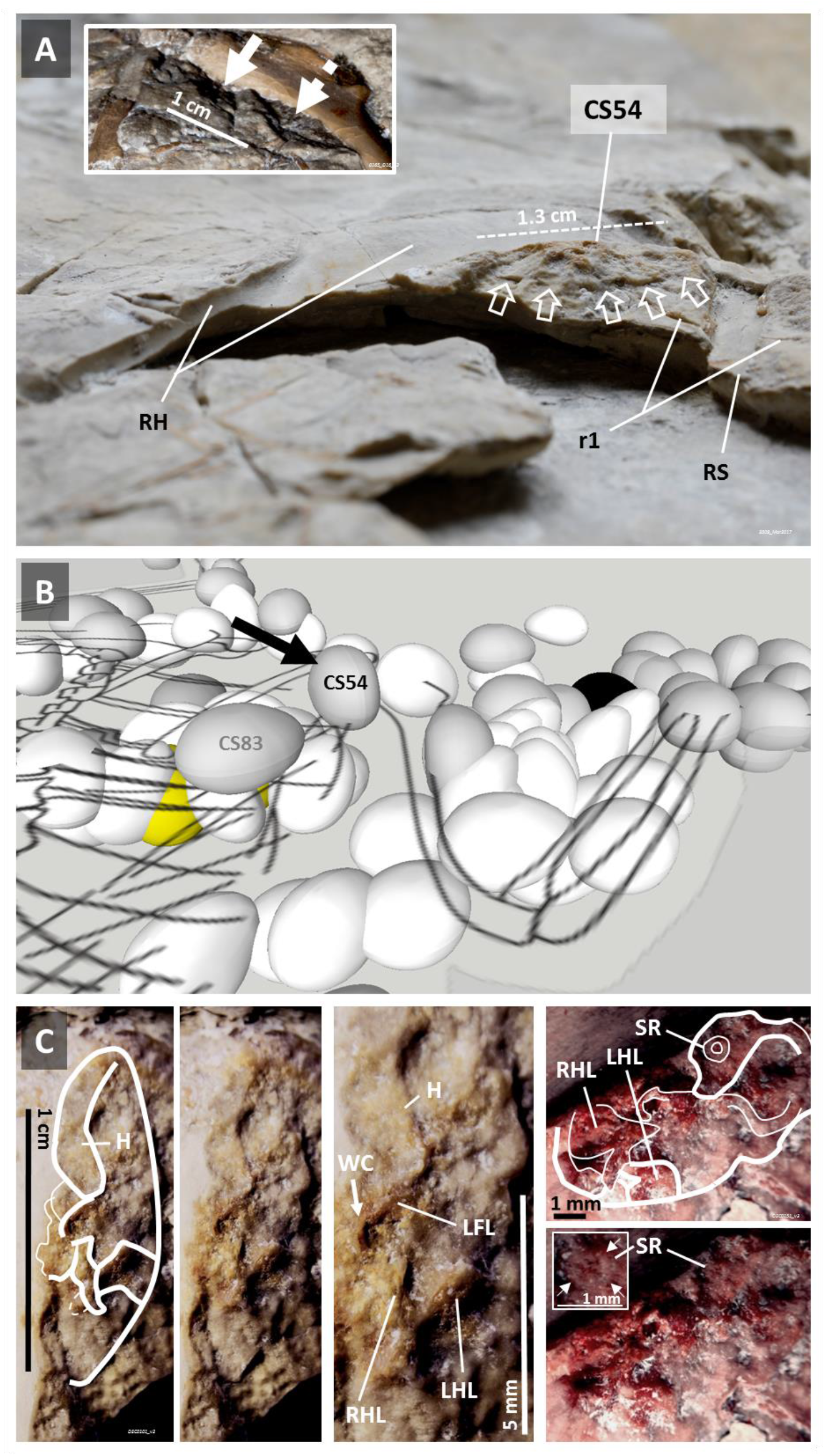
Two eggs lie dorsal to Berlin specimen’s torso. Periphery of CS54 on the surface of a counterslab fragment, photographed here in angled view under natural light and without contrast enhancement, against the posterior rim of the head of the right humerus (RH), near a thoracic rib (r1) and the right scapula (RS; **A**). Inset shows the 3D impression of the CS54 egg in the main slab (full arrow), medially to a depression compatible to that from another egg lost during slab separation (dashed arrow). An angled view of a 3D model indicating the position of eggs CS54 (arrow) and CS83 over the dorsal surface of the specimen’s right hemi-thorax is shown in **B**. CS54 encloses a directly visible fetus with surface morphology preservation (**C**). Aspects of the fetus encased within the outline of CS54 in microphotographs taken under stereomicroscopy using different light incidences are shown. The amplified sclerotic ring (SR) of this fetus is shown in the inset and indicated by white arrows. WC – Wing claw from a left wing finger; H – Head; LFL – Left forelimb; LHL – Left hindlimb; RHL – Right hindlimb; SR – Sclerotic ring.

No calcified eggshells could be detected in Berlin specimen’s fossil slabs, only a peripheral membrane-line structure 0.1 mm or less in thickness that will be presented below together with similar evidence from other *Archaeopteryx* fossils (see below).

### Beaked hatchlings located in the Berlin specimen’s retrocurved neck region

In addition to the MSH1 and MSH2 eggs, and what appears to be a small beaked cranium half a centimeter in size that overlays MSH1 (MSH28 in Figure 13), a hatchling with dislocated mandible was identified in the retrocurved neck area of the Berlin specimen (H1MS in Figure 18).

**Figure 18.**
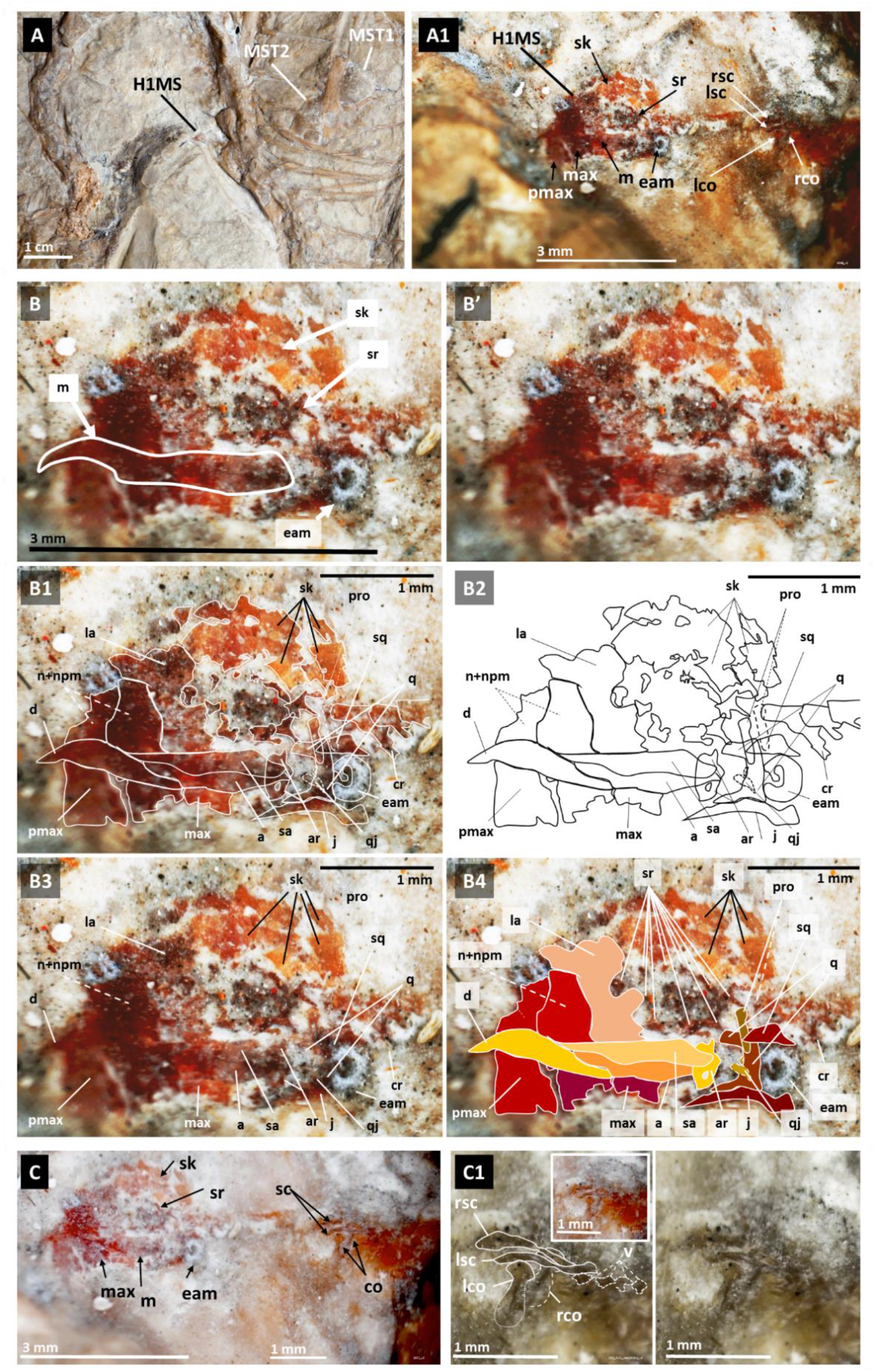
Berlin specimen’s H1MS and its bone preservation. H1MS is located in a depression between the MSH1 and MSH2 eggs on one side and the vertebral column of the Berlin specimen on the other (**A**). The head, neck and upper thorax of H1MS are shown and individual bones from these body segments are indicated (**A1**). The outline of H1MS’s mandible shows how it is dislocated and superimposes on the rest of its head (**B** and **B’**). A tentative identification of H1MS’s cranial bones is shown in **B1**-**B4**. The head, neck and upper thorax of H1MS (**C**) and a magnification of the latter (**C1**) are shown. The distal ends of the scapulae are difficult to discern. a – angular; ar – articular; co – coracoid; cr – cervical rib; d – dentary; eam – external acoustic meatus; j – jugal; la – putative lachrymal; lco – left coracoid; lsc – left scapula; m – mandible; max – maxillary; n + npm: nasal process of the maxilla and nasal bones; pmax – premaxillary; pro – prootic (continuous line left sided; dashed line right sided); q – putative quadrate; qj – uncertain contour of the quadratojugal; rco – right coracoid; rsc – right scapula; sa – surangular; sc – scapula; sk – skull; sq – putative squamosal, sr – sclerotic ring.

H1MS’s head morphology is consistent with that of the egg-enclosed fetus with toothed beak of MSB5 that lies next to it (see Figure 9), approximately 3 mm in size each, but smaller than MSH28’s.

H1MS shows bone preservation of the cranium (Figure 18B), part of the cervical vertebral column and shoulder girdle (Figure 18C-C1).

Nearby, a beaked head was observed in slab embedded MSH15, detected via 3D software analysis of CT scan data and shown in Figure 19.

**Figure 19.**
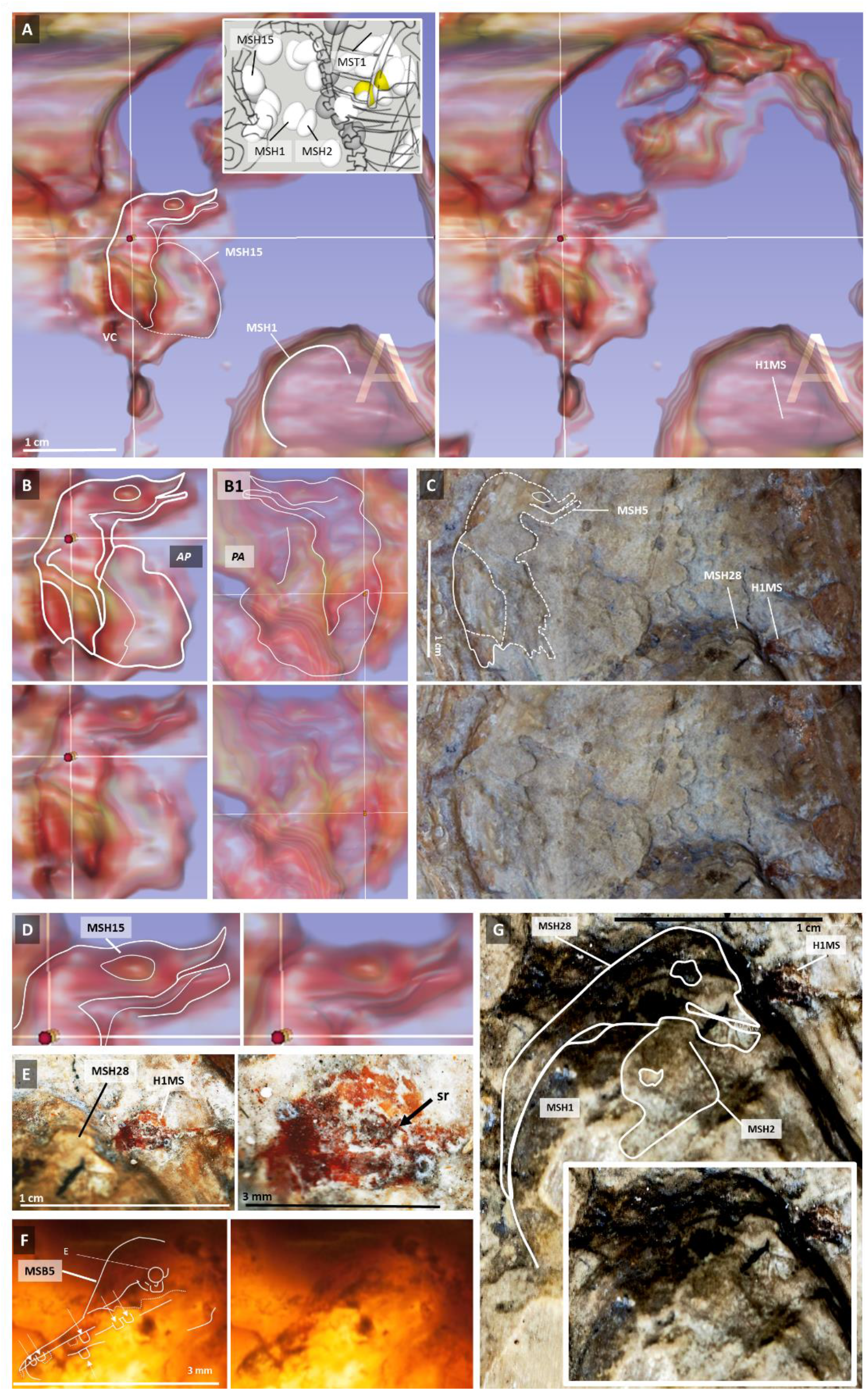
MSH15 is a beaked hatchling embedded in the main slab that appears to have died hatching. MSH15 identified via 3D software analysis of CT scan data (**A**), in antero-posterior (**B**) and posterior-anterior views (**B1**). Once identified via CT scan analysis, its outline was found and represented on the surface of the main slab (**C**). Head morphology of MSH15 (**D**) and hatchling H1MS (**E**), both located in the retrocurved neck area of the Berlin specimen, and that of egg-enclosed MSB5 located in the adjacent thoracic region (**F**). One centimeter away from MSH5, MSH28 lies atop MSH1 and MSH2, shown here for comparison (**G**).

Just one centimeter away from MSH15, and two from H1MS, exposed on the main surface but with body orientation inverted in relation to that of the Berlin specimen, lying against the back of the adult animal’s skull, is another beaked hatchling, designated H2MS, with head to tail outer body preservation (Figure 20). In addition to its beaked head, H2MS shows parallel stumps compatible with those of remiges shafts at a location compatible with that of its fossilized wing, which appears to have fossilized in clinging position.

**Figure 20.**
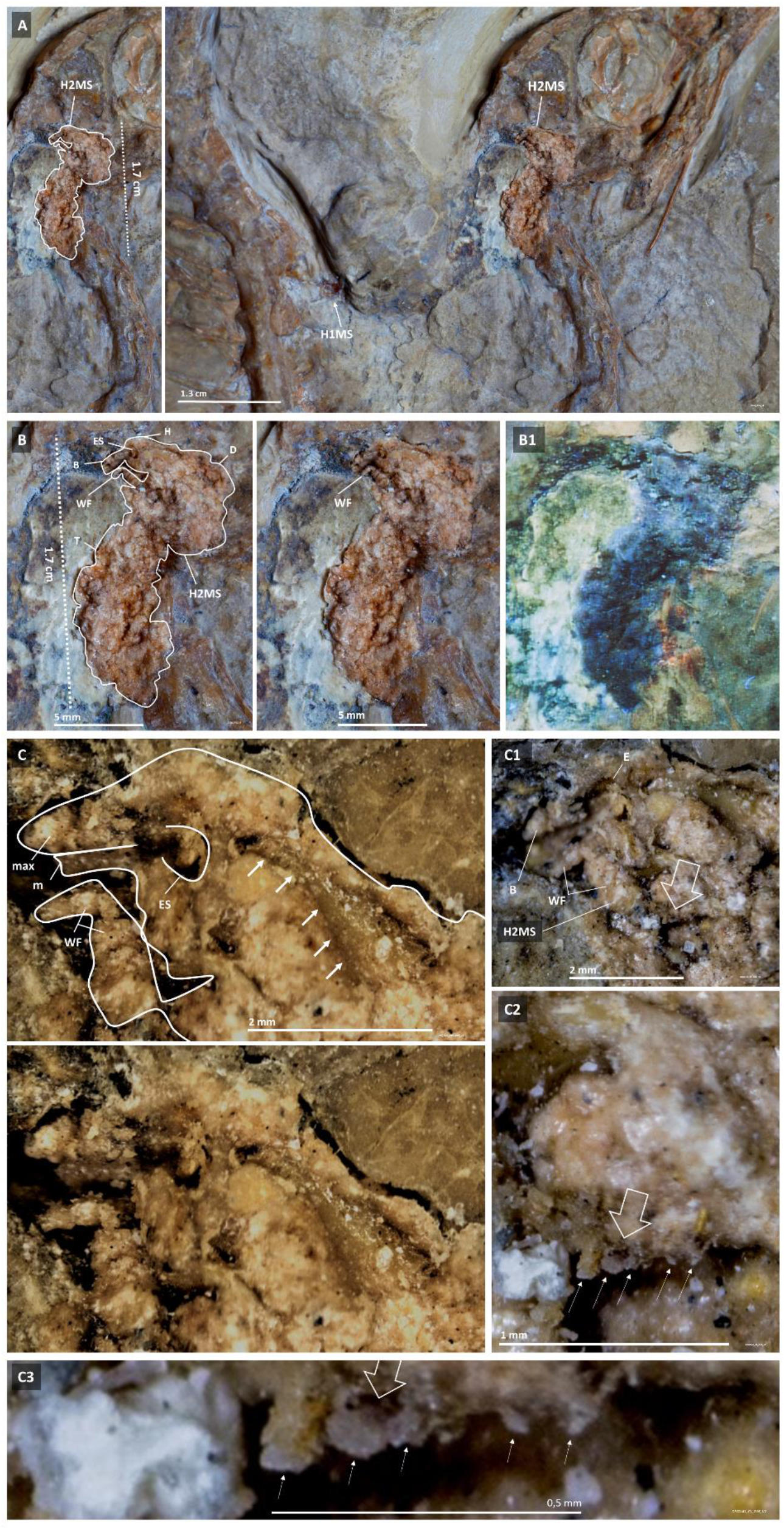
H2MS is surface exposed against the skull of the Berlin specimen within its retrocurved neck region. H2MS is directly visible (**A**) from head to tail under natural (**B**) and UV light (**B1**). The back and left side of its head (**C**) is shown under stereomicroscopy. The non-segmented structure in the cervical area (white arrows in C) may represent its neural tube. At a location compatible with that of its right wing (open arrow), parallel stumps compatible with remiges shafts are shown (**C1**-**C3**). B – beak; D - Dorsum; E – Eye; ES – Eye socket; H – head; m – mandible; max – maxilla;– skull; sr – sclerotic ring; T-tail; v-vertebrae; WF – wing finger.

The retrocurved neck region of the Berlin specimen shows several fossil findings within 3.5 cm of each other: hatchlings H1MS, H2MS, MSH15, egg-enclosed MSB5, MSH27, MSH2 and MSH28 (Figure 21A). The outline of another hatchling, designated H3MS, apparently also fossilized in clinging position, is outlined in Figure 21A. All these structures plus the thoracic egg mass THAT lies ventral to the adult animal are encompassed within a 7 cm in diameter area (Figure 21B).

**Figure 21.**
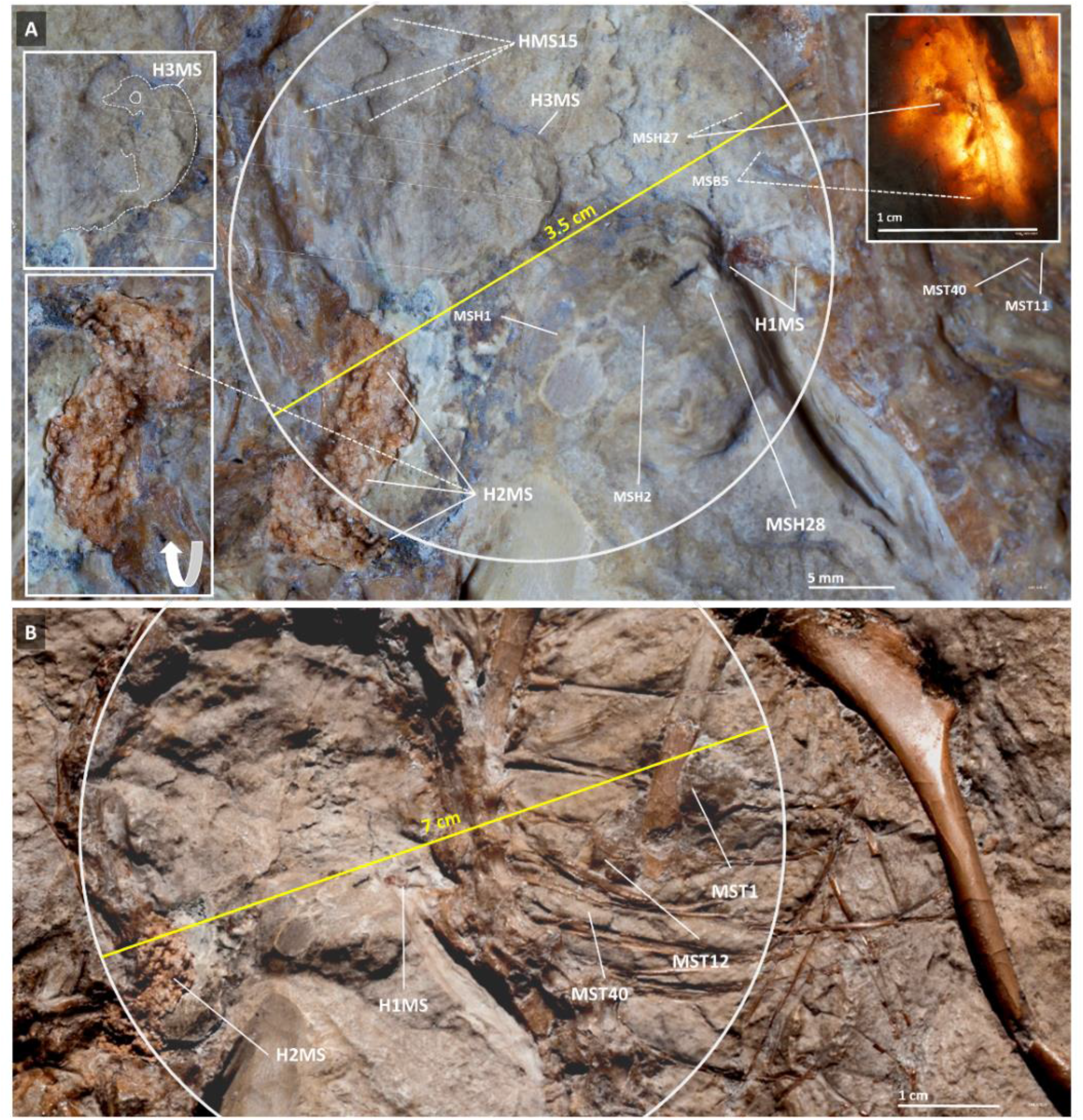
Fossil findings within the retrocurved neck region of the Berlin specimen and their proximity to the ventral egg mass. A circle 3.5 cm in diameter encompasses several hatchlings and an egg-enclosed fetus (**A**); a circle 7 cm in diameter also encloses the thoracic egg mass that lies ventral to the Berlin specimen (**B**).

### Egg littering in both slabs of the Berlin specimen

Egg vestiges were also found outside the Berlin specimen’s feathered body perimeter in the main slab (Area 4 in Figure 3): MSS14’s beaked head shown in Figure 22 is slightly better outlined than that of MSS27. These are embedded in the sediment in opposite sides of the main slab and are among 12 egg vestiges singled out in several locations of Area 4 (see Movie 1).

**Figure 22.**
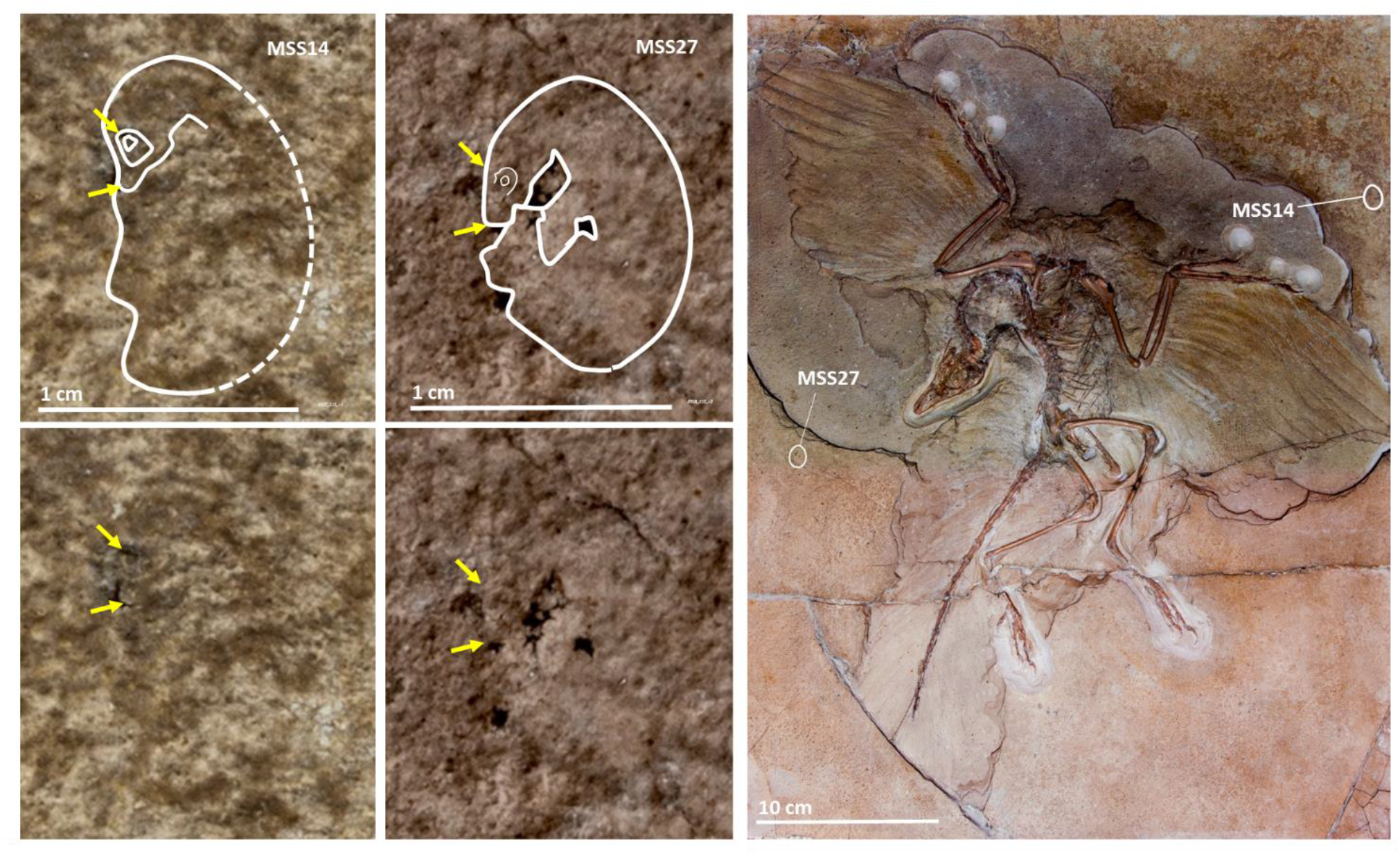
Main slab egg remnants outside the body perimeter of the Berlin specimen of *Archaeopteryx*. MSS14 and MSS27, seen here under stereomicroscopy, are embedded in opposite sides of the main slab. Arrows are provided for guidance.

Similarly, numerous vestiges of egg-enclosed fetuses with surface morphology preservation were identified via direct observation in different locations of the surface of the slab proper of the counterslab (Area 5; Figure 23 and Movie 1). In addition, 4 eggs with some degree of 3D preservation were identified via 3D software analysis of CT scan data embedded in the slab proper of the counterslab. One of these, designated CS53, caused a matching 3D impression in the cranium of the Berlin specimen in the main slab (Figure 24 and Movie 1).

**Figure 23.**
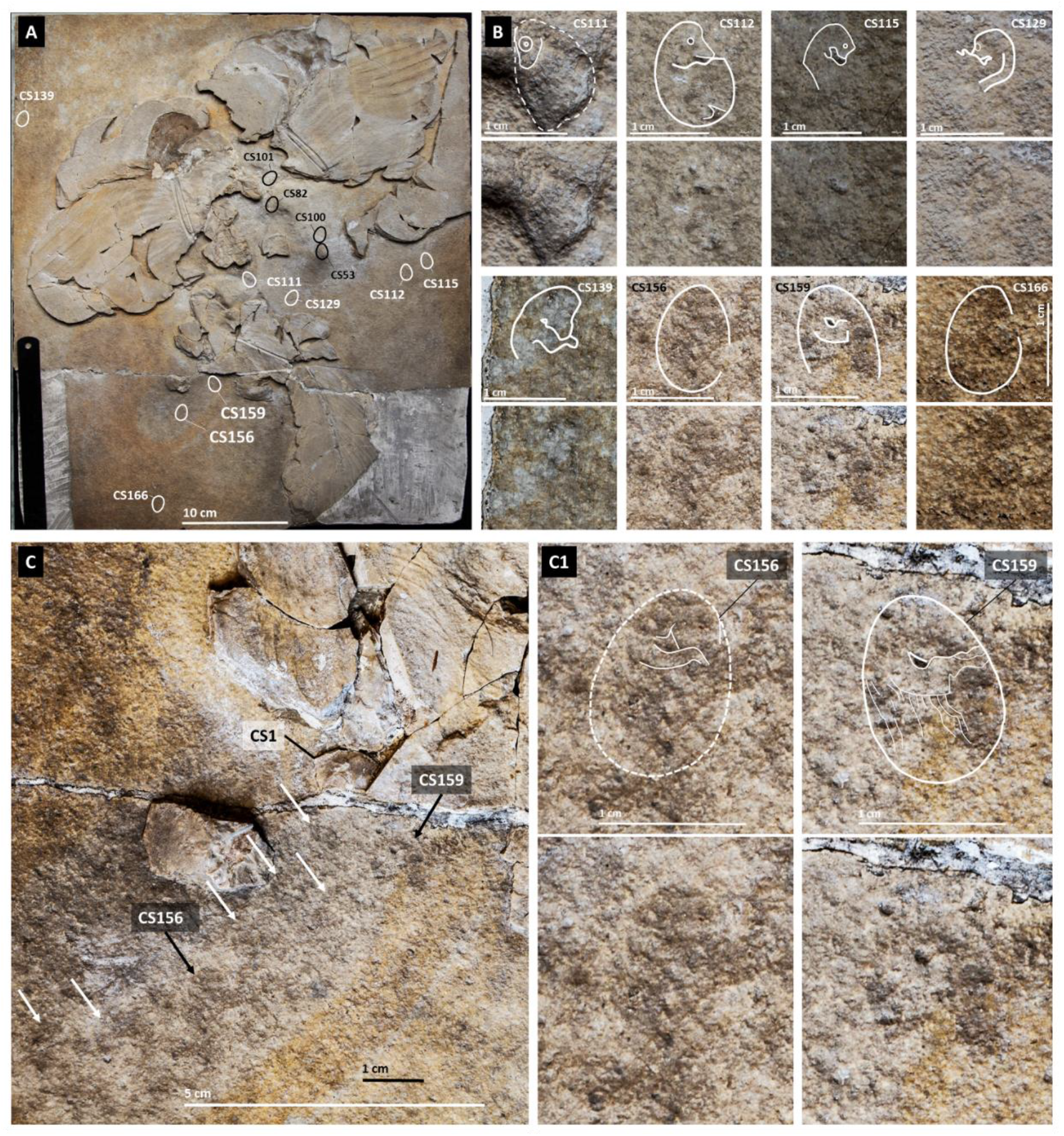
Remnants of egg-enclosed fetuses located outside the body perimeter of the Berlin specimen of *Archaeopteryx* in the counterslab. The location of impressions of egg-enclosed fetuses exposed on the surface of the slab proper of the counterslab and 3D preserved eggs detected via 3D software analysis of CT scan data is respectively indicated with white and black outlines in **A**. The former are magnified in **B**, **C** and **C1**. The latter are in Figure 24. The area in C1 corresponds to the right foot of the Berlin specimen, where CS1 shown in Figures 15 and 16 is located. Note numerous oval impressions indicated by white arrows. Contrary to CS1, which is located in a counterslab fragment, egg vestiges CS156 and CS159 (magnified on the right) are located on the slab proper of the counterslab.

**Figure 24.**
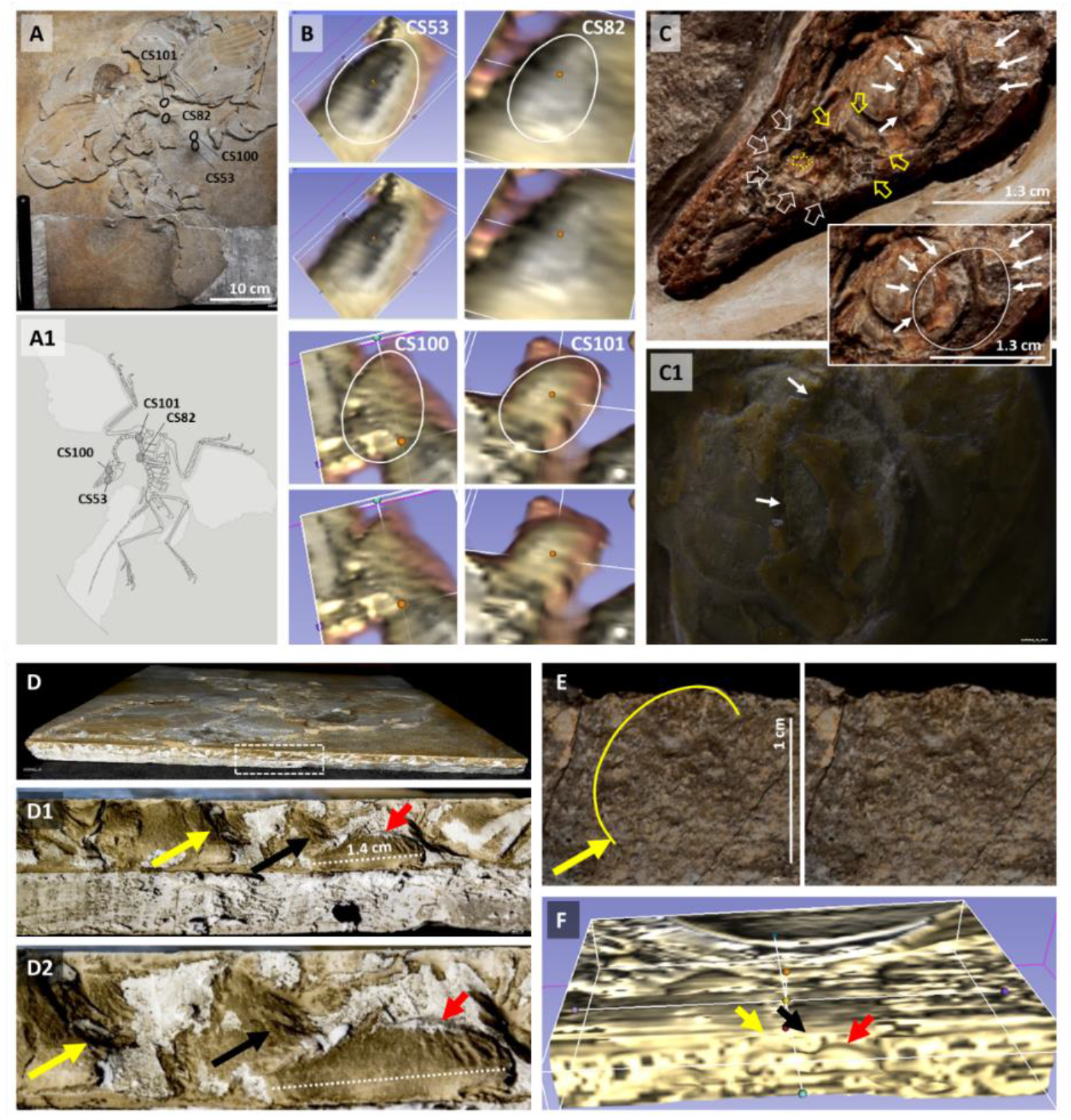
Eggs with 3D preservation are embedded in the slab proper of the counterslab of the Berlin specimen and egg remnants extend to its margins. The location of four eggs embedded in the slab proper of the counterslab detected via 3D software analysis of CT scan data is shown in **A** and in a 3D model in **A1**. They are individually shown in **B**. 3D impressions in the cranium of the Berlin specimen compatible with those caused by eggs are indicated by three sets of arrows in **C**. CS53 appears to have caused the concavity indicated by open yellow arrows, while other eggs are likely to have caused the depression to its left (open white arrows) and the one located on and to the right of the sclerotic ring (white full arrows; outlined in the inset). A magnified view of the sclerotic ring is provided in **C1**. The counterslab is only exposed in very small patches in its upper margin while its other margins are mostly covered in plaster or similar material (data not show). Macrophotography of the top margin of the counterslab (**D**), magnified in **D1** and **D2**, suggestive of abundant egg remnants (arrows). An egg outline is detectable in frontal view of the slab proper (yellow arrow in **E**) that matches a prominence indicated in D2 (yellow arrow) and a 3D structure under 3D software analysis of CT scan data (yellow arrow in **F**). The latter technique suggests a heterogeneous slab composition compatible with an egg amalgamation.

Given the abundance of egg vestiges in the slab proper of the counterslab, its margins were investigated. Egg remnants with some degree of 3D preservation were found to extend to its margins (Figure 24D-F and Movie 1). In turn, the margins of the main slab were not available for study as they remained protected by the permanent exhibition mount even after opening of the sarcophagus-like set up of this mount and exposure of its ventral side (*10*).

### Egg number and distribution in Berlin specimeńs fossil

The areas that represent the body perimeter of the Berlin specimen and the area anterior to its folded wings (Areas 1, 2 and 3) in addition to Area D1 (see Figure 3) were considered to estimate the total number of eggs of the Berlin specimen’s clutch and organized into Groups I-IV (see Materials and Methods).

Several examples of Group I eggs were shown above. Examples of Groups II, III and IV eggs are shown in Figures 25-27, respectively (see Movie 1).

**Figure 25.**
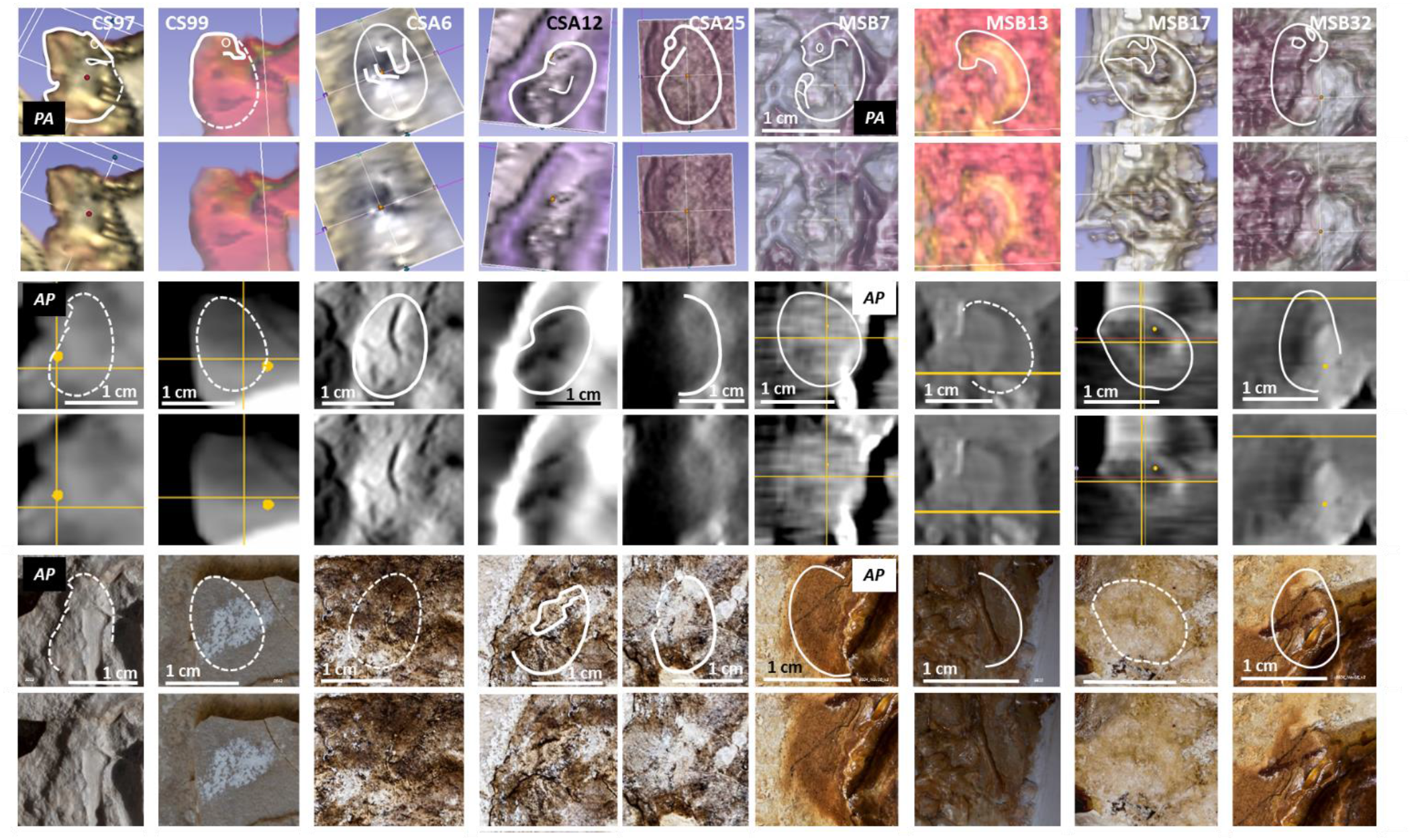
Egg-enclosed fetal remnants with surface morphology preservation detected via 3D software analysis of CT scan data (Group II eggs). Nine of fourteen of these structures are shown; CS1 (Figures 15 and 16), MSH2 (Figure 13) and MSH15 (Figure 20) are other examples; CS83 (mentioned in Figure 17) and MSP12 were not shown. The outline of each fetus with surface morphology preservation detected via 3D software analysis of CT scan data is shown on top, a 2D CT scan section of the same area is shown in the middle and a macrophotography of the corresponding slab surface is shown at the bottom. Dashed outlines are not detectable on the slab surface or via 2D CT scanning. AP – Posterior-anterior view; PA – Postero-anterior view.

**Figure 26.**
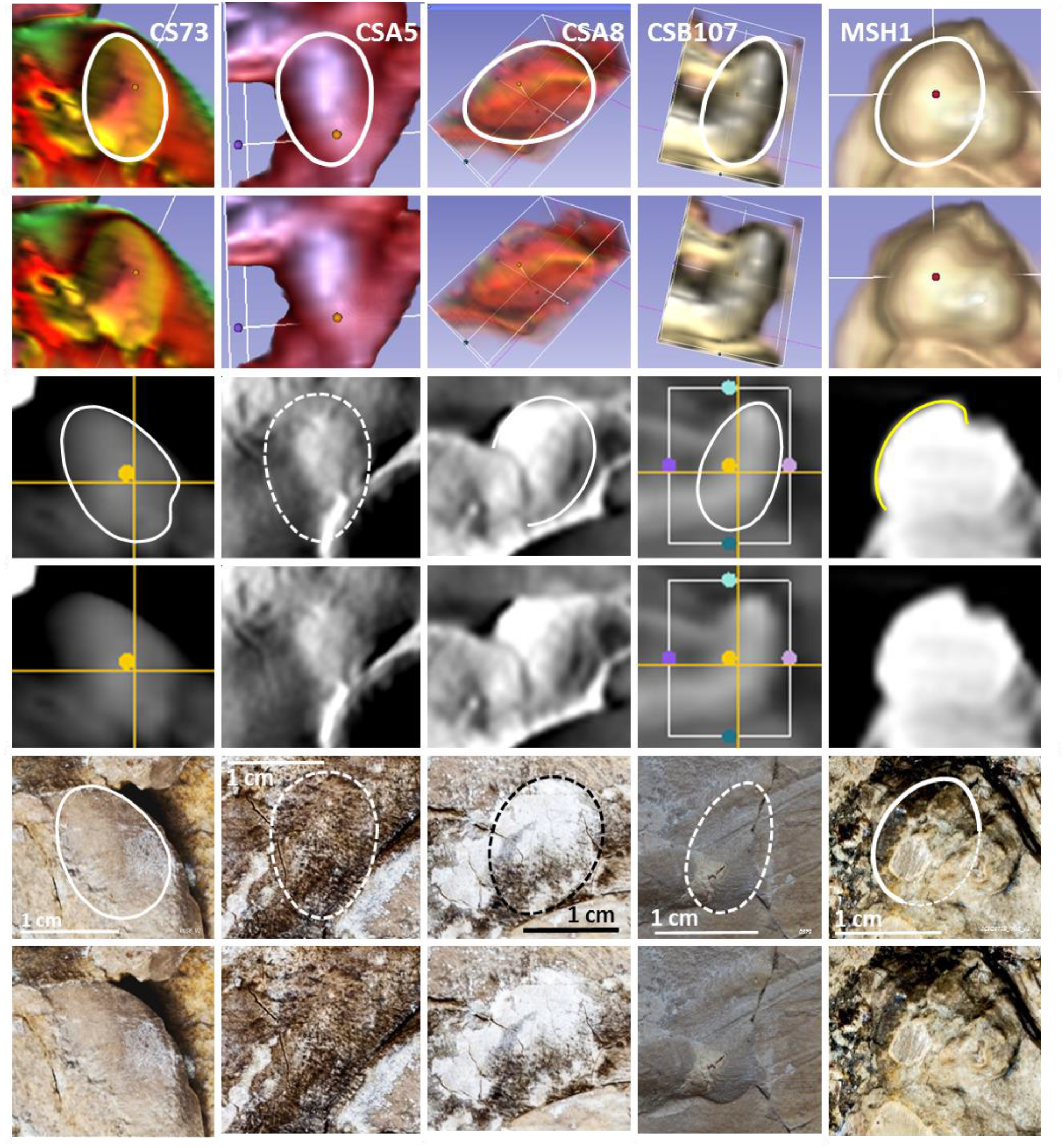
Eggs without a detectable fetus that were identified via 3D software analysis and conventional 2D images of CT scan data (Group III eggs). Five of sixty three structures are shown. The outline of each egg detected via 3D software analysis of CT scan data is shown on top, a 2D CT scan image of the same area is shown in the middle and a macrophotography of the surface of the corresponding slab is shown at the bottom. Dashed outlines are not detectable on the slab surface or via 2D CT scanning. Other examples of Group III eggs are not shown here. All group III eggs and their location are listed in Figure 28.

**Figure 27.**
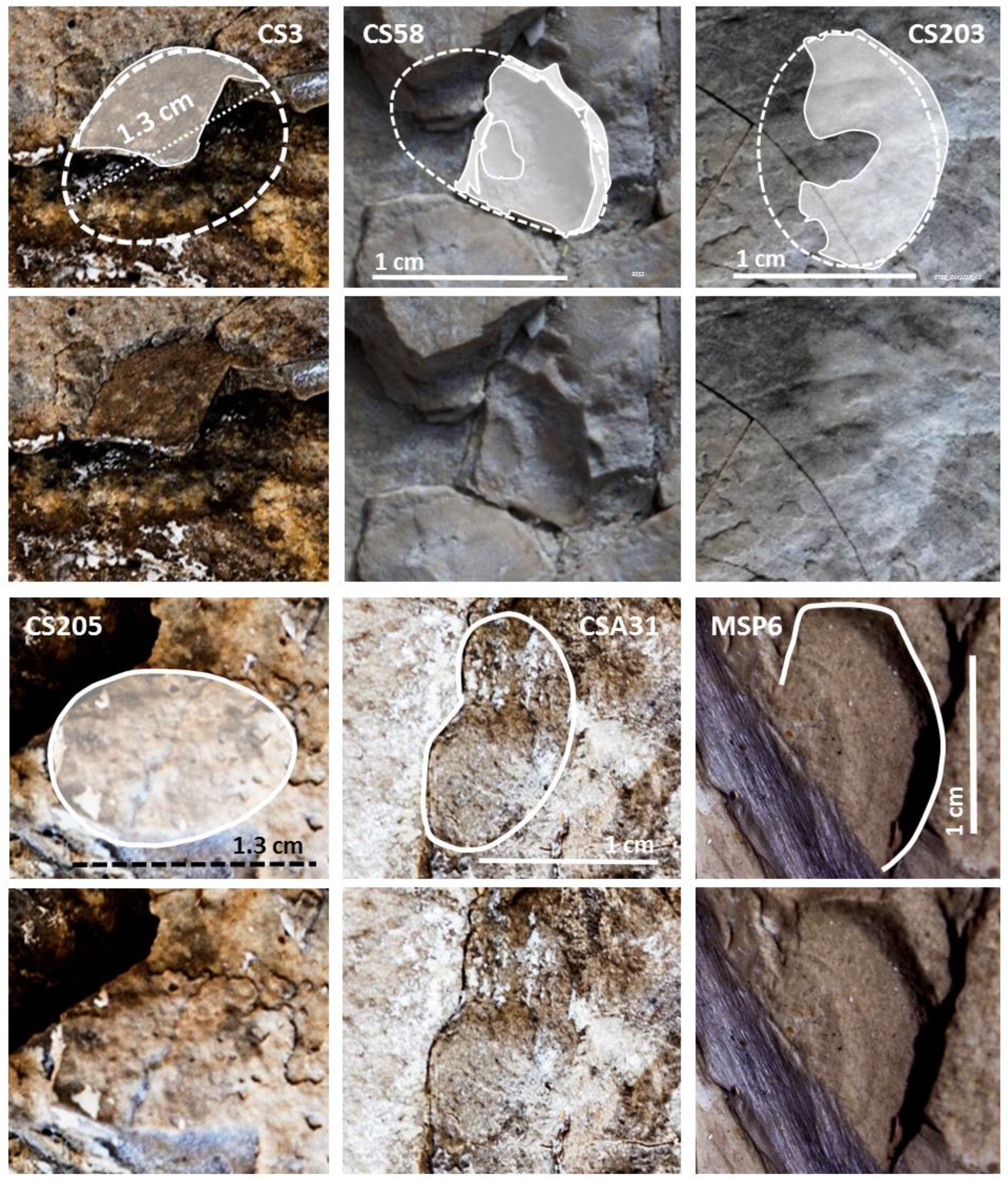
Group IV eggs. Examples of Group IV eggs are shown in the form of a convex fragment (CS3; the dashed outline delimits an impression on the slab surface), a concave fragment (CS58), an egg impression (CS203), a flattened egg remnant with complete outline in coronal view (CS205), a directly visible egg with distorted but complete outline in sagittal view (CSA31), and a 3D egg impression or “egg ghost” (MSP6). All these structures are approximately 1,3 cm long. Other Group IV eggs not shown here and their location are listed in Figure 28.

Except for the MSS14 egg shown in Figure 22 (MSS27 shown in the same figure was classified within Group IV), all other eggs with a directly visible fetus or one detectable via CT scan analysis reside within Berlin specimen’s body perimeter (Areas 1, 3 and D1) or in the adjoined area anterior to its body (Area 2).

Eggs that are anterior to the Berlin specimen are mostly disposed laterally against its folded wings, whereas those within its body perimeter are found alongside its body axis.

In contrast to eggs with different levels of 3D preservation can be observed within the perimeter of Berlin specimen’s body, only egg vestiges were found in Area 4.

105 eggs can be counted if only eggs with a detectable fetus or radiologic evidence located immediately ventral or dorsal to its body, within the perimeter of its wings and tail, or within Area 2 that is anterior to the adult animal are considered (excluding MSS14 from Area 4; Figures 28 and 29). If egg vestiges and impressions undetectable via CT scan analysis (Group IV) are added, the number of eggs within the same areas rises to 135, well above the 28 eggs initially reported (*10*). There are almost as many eggs anteriorly to the animal (in Area 2) as there are ventrally to its body and wings. Importantly, only two eggs were identified dorsally to the specimen (Area D1).

**Figure 28.**
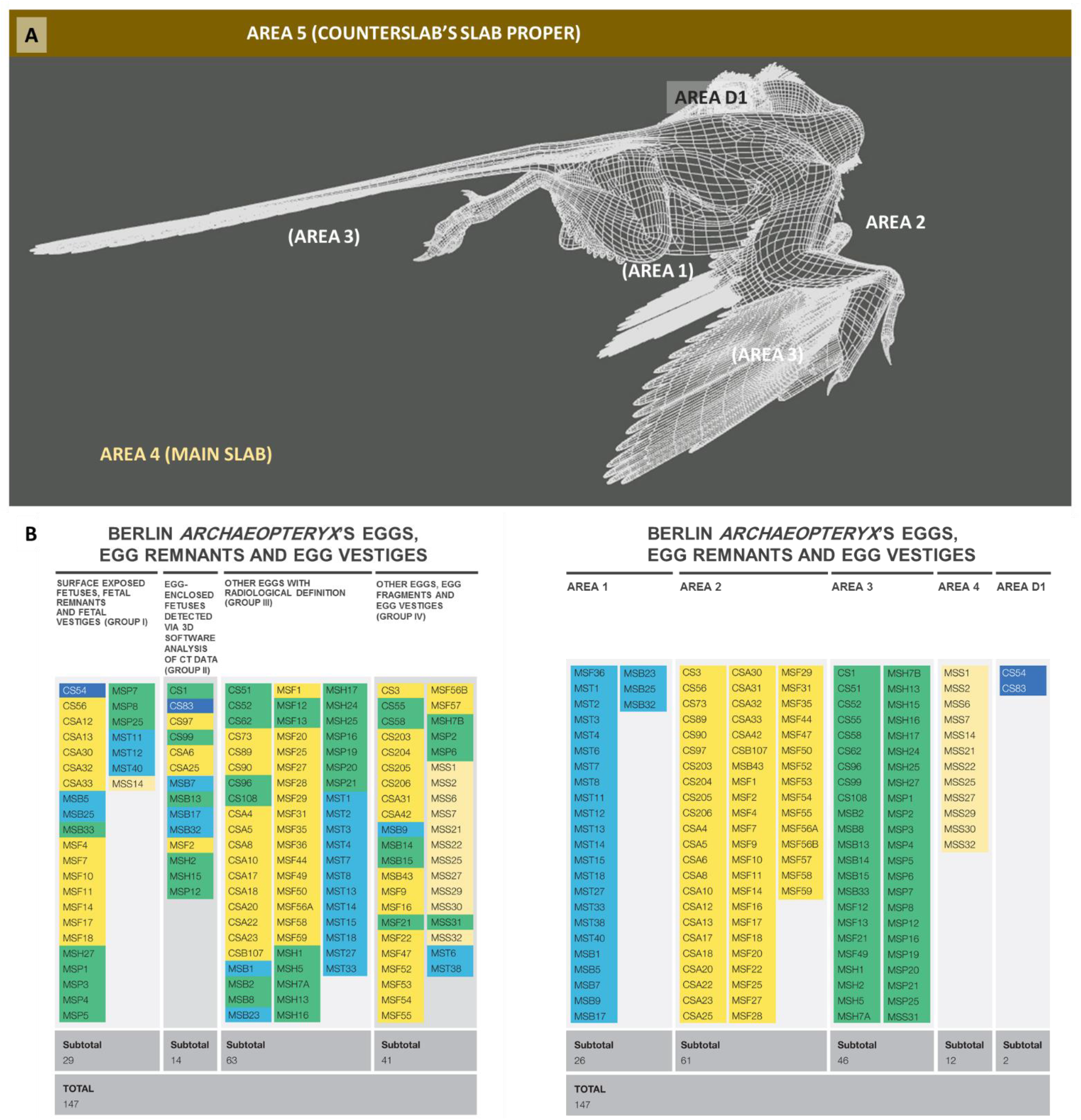
Berlin specimen eggs per type of evidence and fossil location. The fossil areas defined for this study are shown in **A**. Each egg identified in **B** is labeled in a color that corresponds to its location (B, right).

**Figure 29.**
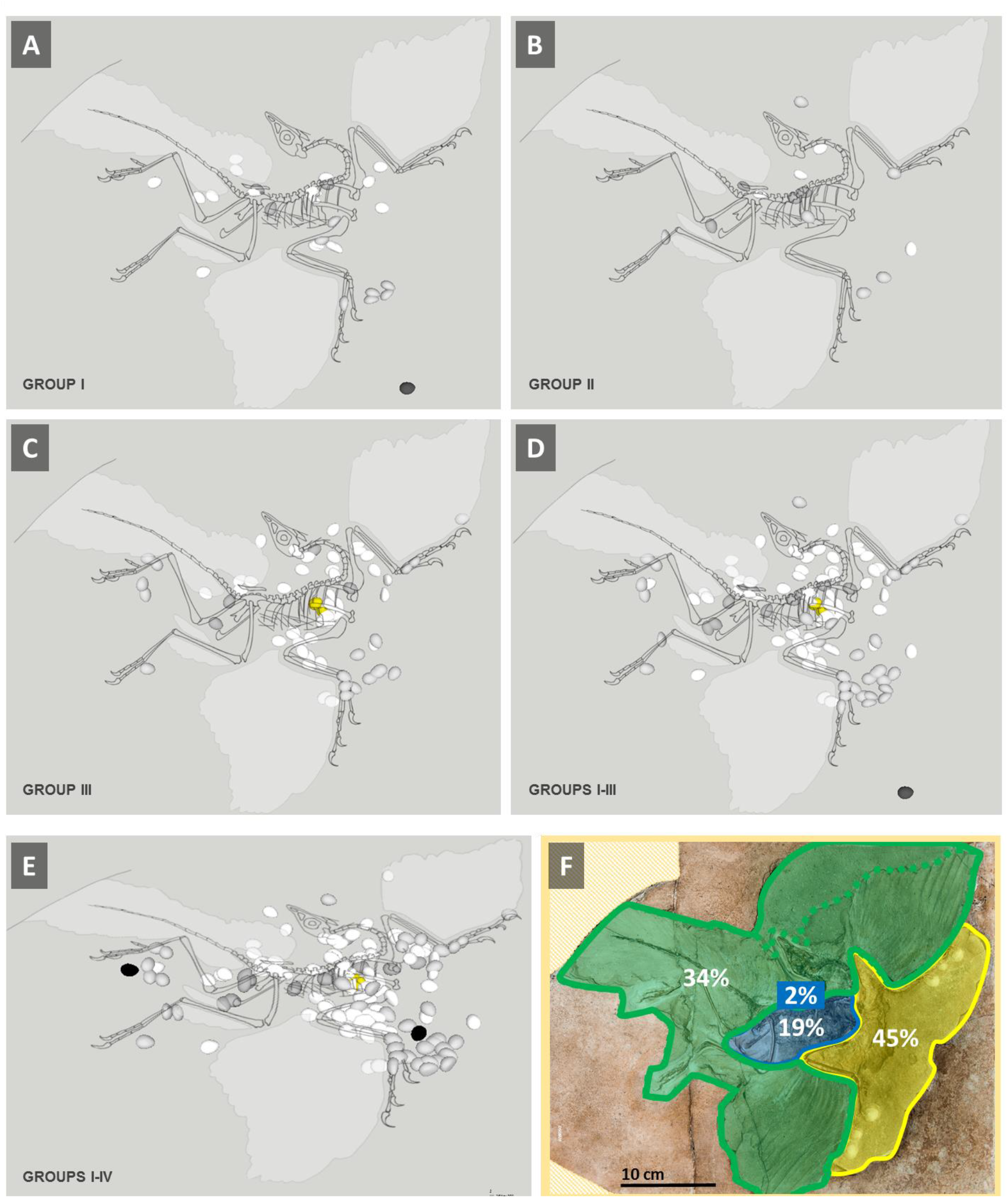
Egg distribution in Berlin specimen’s fossil. Eggs and egg remnants from Groups I (**A**), II (**B**) and III (**C**) from Areas 1, 2, 3, 4 and D1 (Area 5 was not considered). All eggs from Groups I-III are represented in **D** and all eggs from Groups I-IV, including the eggs detected in Area D1, in **E**. The relative egg distribution of all eggs from Groups I-IV is shown in F, with Area 1 in blue, Area 2 in yellow, Area 3 in green and Area D1 over Area 1. Non-original areas of the main slab are shown filled in beige (*10*). MST1 and MST2 eggs are shown in yellow for reference. Eggs in white were identified in the front surface of the main slab, those in light grey in counterslab fragments, the ones in dark grey in the back of the main slab (mostly alongside the vertebral column and therefore difficult to see in these images) and the ones in black are 3D impressions or “ghost eggs” (the one near the left foot resides in the main slab and the one near the right wing in a counterslab fragment). The dark egg in D is MSF14 that lies within Area 4.

### Fossilized posture of all known *Archaeopteryx* specimens

All articulated *Archaeopteryx* specimens fossilized with flexed thighs and knees, abducted and flexed wings at the elbows and wrists, and retroflexed neck. Their pose matches different views of the same nesting posture: dorsolateral in the Berlin specimen, lateral in the Eichstatt, Solnhofen, Teylers and 12^th^ specimens, antero-lateral in the Munich specimen, dorsal in the London specimen, ventral in the Thermopolis specimen and ventro-lateral in the 11^th^ specimen (Figure 30).

**Figure 30.**
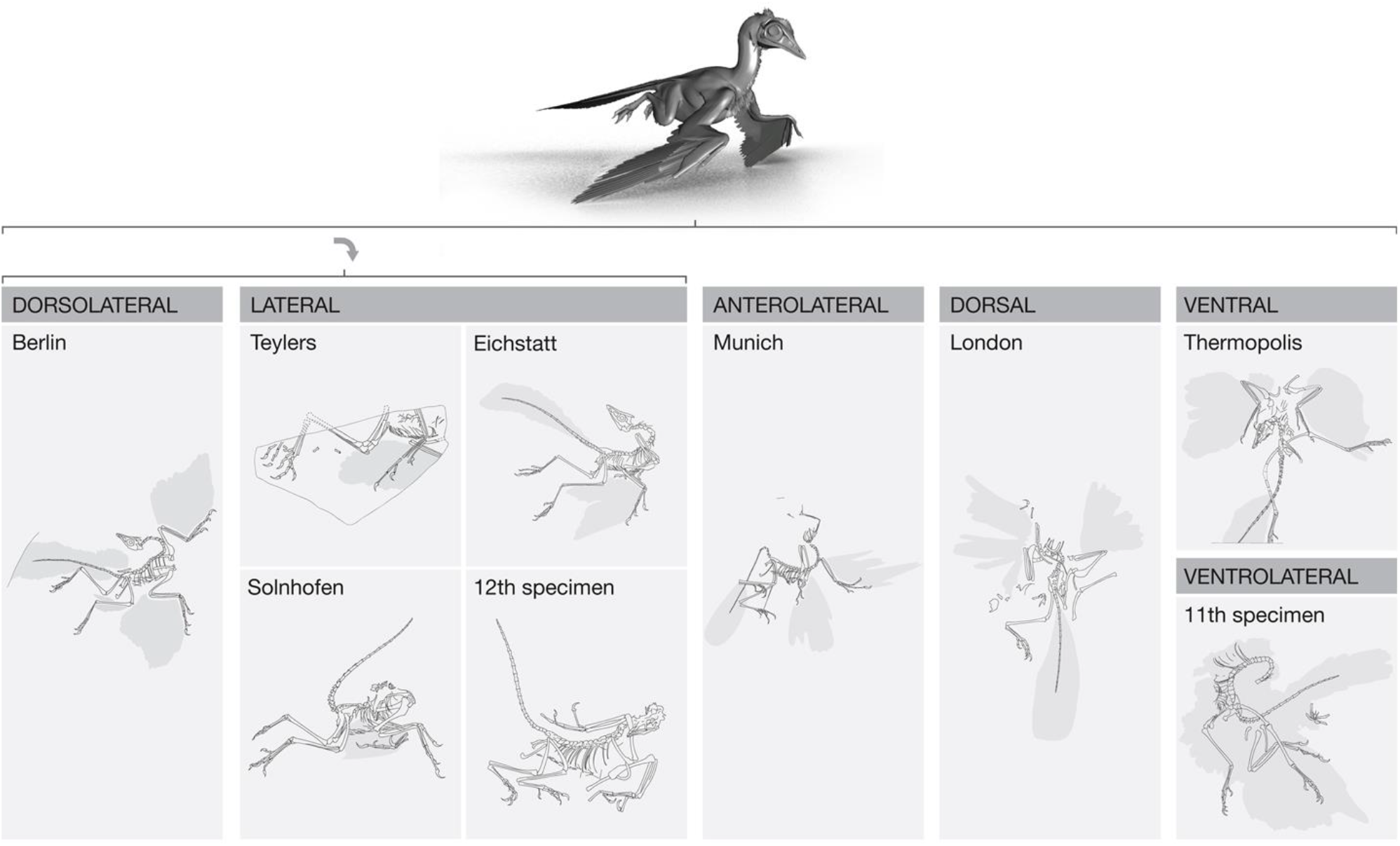
All articulated *Archaeopteryx* specimens fossilized in body positions that match different views of the nesting posture.

### A ventral egg mass, hatchlings and evidence of egg littering the Teylers specimen

A mass is ventral to the thoracoabdominal region of the Teylers specimen of *Archaeopteryx*. CT scan views of this mass show its higher radiodensity and honeycomb appearance. TSS1, measuring 1.3 cm in its longer axis, is the most evident oval convexity at the surface of this mass, among other convexities and concavities of similar characteristics, with a matching impression on the other slab. As seen with eggs from the thoracic egg mass of the Berlin specimen, TSS1 caused distortions on overlying gastralia and encloses a fetus with surface morphology preservation (Figure 31).

**Figure 31.**
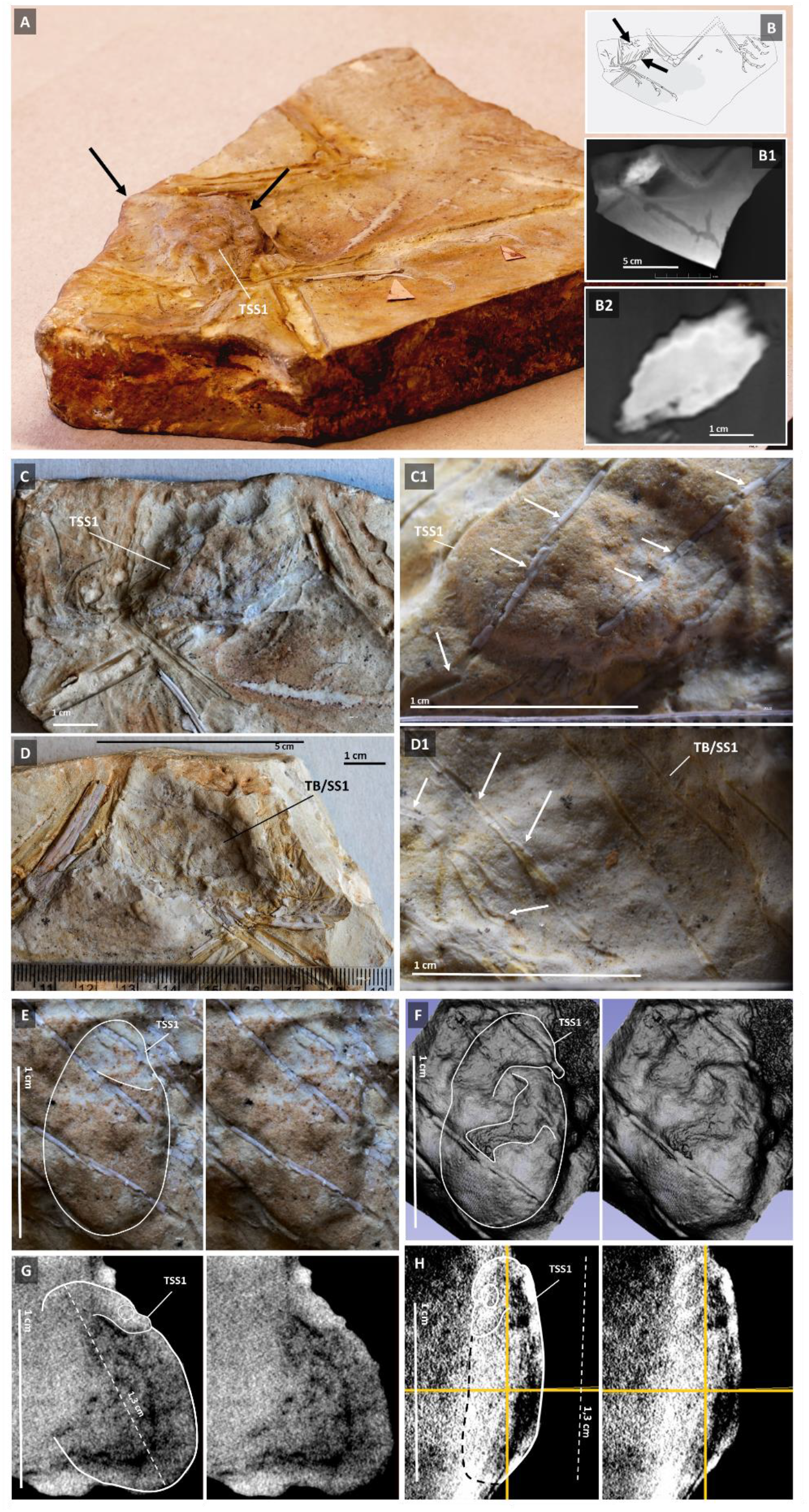
Egg mass with an egg enclosed fetus with surface morphology preservation in the abdominal area of the Teylers specimen of *Archaeopteryx*. **A:** Angled view of the fossil’s smaller slab or counterslab. Black arrows indicate the mass ventral to the specimen. **B**: Diagram with the representation of the specimen’s skeleton in relation to a coronal CT scan view (**B1**), showing the higher density of the egg mass in relation to the surrounding sediment and its magnification (**B2**). **C**: Enlarged view of the egg mass in the counterslab that makes its globular units apparent. **C1**: Outline of the TSS1 egg under macrophotography. **D**: Enlarged view of the impression of the egg mass in the main slab. **D1**: Outline of the TSS1 egg impression in the main slab. TSS1 and the outline of its enclosed fetus under macrophotography (**E**), 3D software analysis of CT scan data at approximately 45 μm resolution (**E**), as well as coronal (**G**) and sagittal (**H**) 2D CT scan views at the same resolution.

Hatchlings with a 3 to 5 mm head were also identified in the Teylers specimen fossil, near the thoracoabdominal egg mass, with the same dark macroscopic appearance suggestive of exceptional fossil preservation seen in DAA from the Berlin specimen, but here located at the margin of the counterslab (Figures 32-34). TH1SS has a beaked head and a maxilla inserted tooth, in addition to the wing finger crossover characteristic of *Archaeopteryx* (Figure 33). TH2SS is a second hatchling from the Teylers specimen fossil that has a beaked head. Like TH1SS, it also appears to have maxilla inserted teeth. Importantly, TH2SS has a preserved forelimb claw, and, like H2MS from the Berlin specimen, tail preservation (Figure 34).

**Figure 32.**
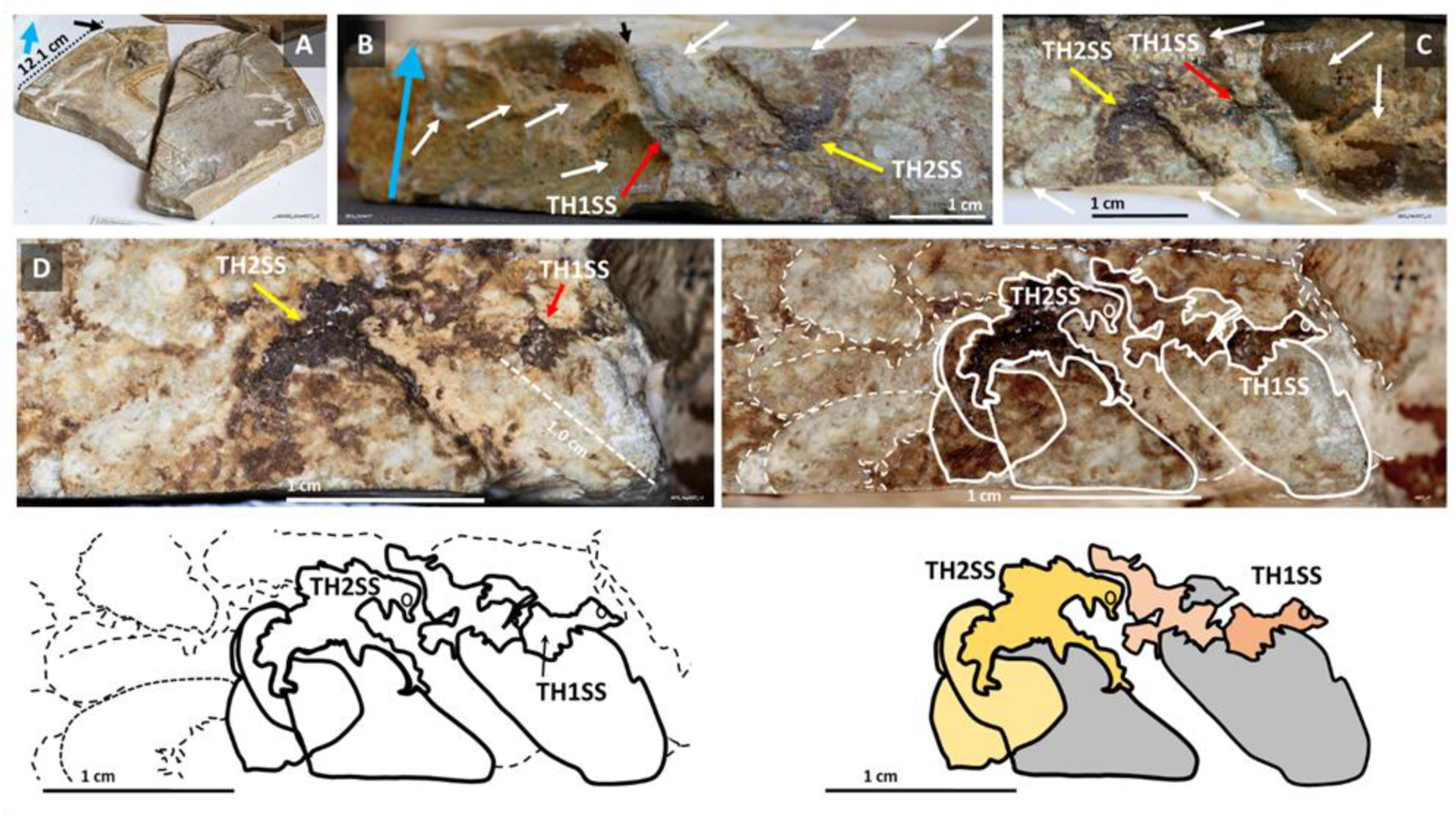
Two hatchlings found in distinctly brown areas of the Teylers specimen fossil. **A:** Area of the counterslab where the hatchlings are located (black arrow). **B**: TH1SS (Teylers Hatchling 1 from the Smaller Slab) and TH2SS (Teylers Hatchling 2 from the Smaller Slab) among egg impressions (white arrows). The blue arrows in A and B are used for orientation. **C**: Reorientation of the image shown in B for easier interpretation. **D**: The body surface remains of THI1SS and TH2SS are distinctly brown and can be outlined among numerous egg impressions.

**Figure 33.**
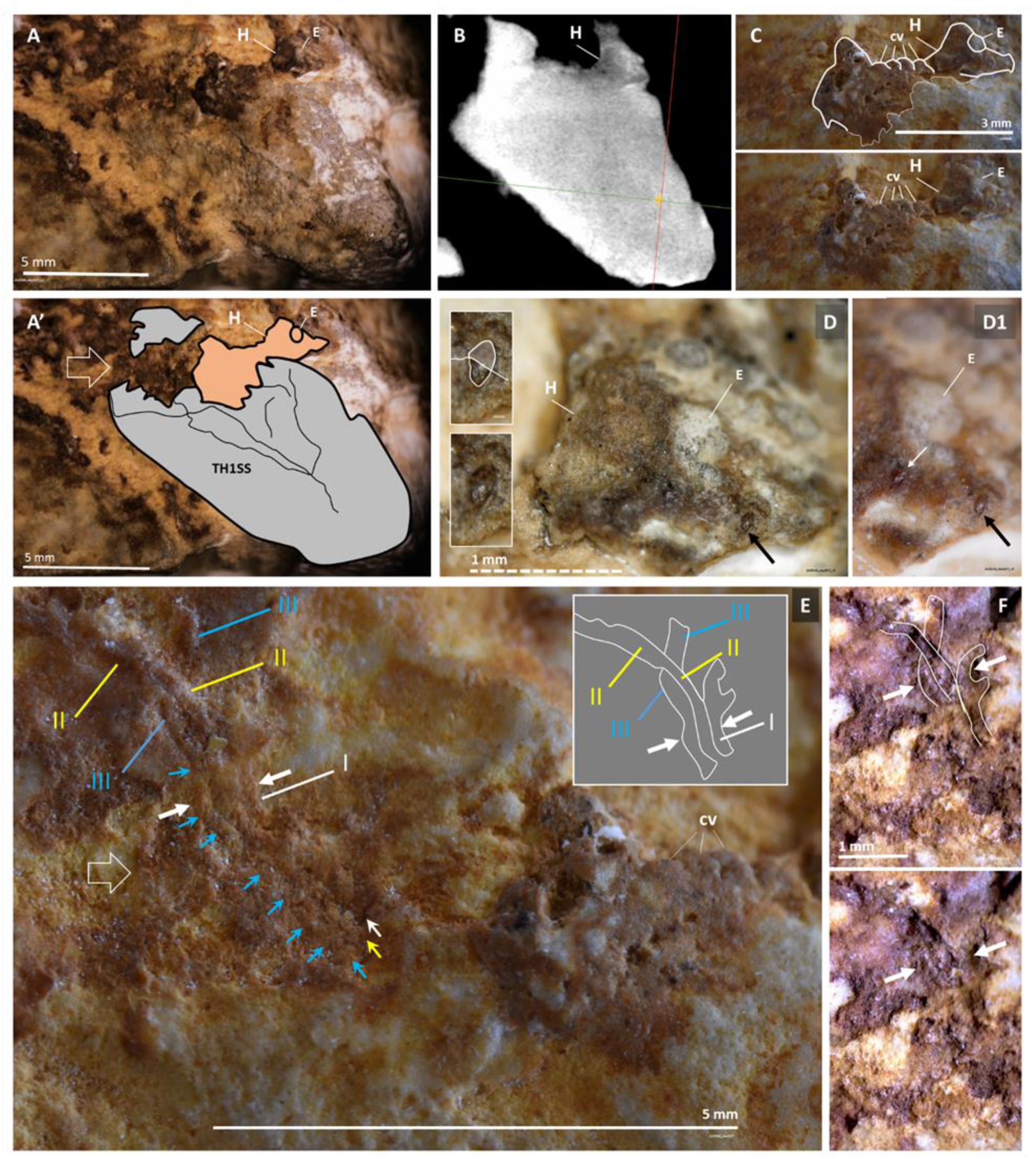
TH1SS, a hatchling from the Teylers specimen, has a beaked head and a maxilla inserted tooth, in addition to the wing finger undercrossing characteristic of *Archaeopteryx*. **A:** Macrophotograph of TH1SS. Open arrow in A’ points at the area where its wing remnant was identified (see below). The illustration denotes that this hatchling seems to be hatching from its egg. **B**: CT scan of TH1SS and associated egg at approximately 45 μm resolution. **C**: upper body of TH1SS. **D**: Head of TH1SS with an inserted tooth in its maxilla (arrow), outlined in the inset. **D1**: Another tooth exposed against TH1SS’s maxilla. **E**: Stereomicroscopy of the wing remnant of TH1SS at the location indicated by the open arrow in A’. The 3^rd^ wing finger (III) crosses under the 2^nd^ (II; see diagram in inset). **F**: The 1^st^ wing finger (I), poorly discernible in E, under a different light incidence. The three fingers of the wing remnant are outlined. Arrows are provided for guidance. cv – Cervical vertebrae; E – Eye; H – Head.

**Figure 34.**
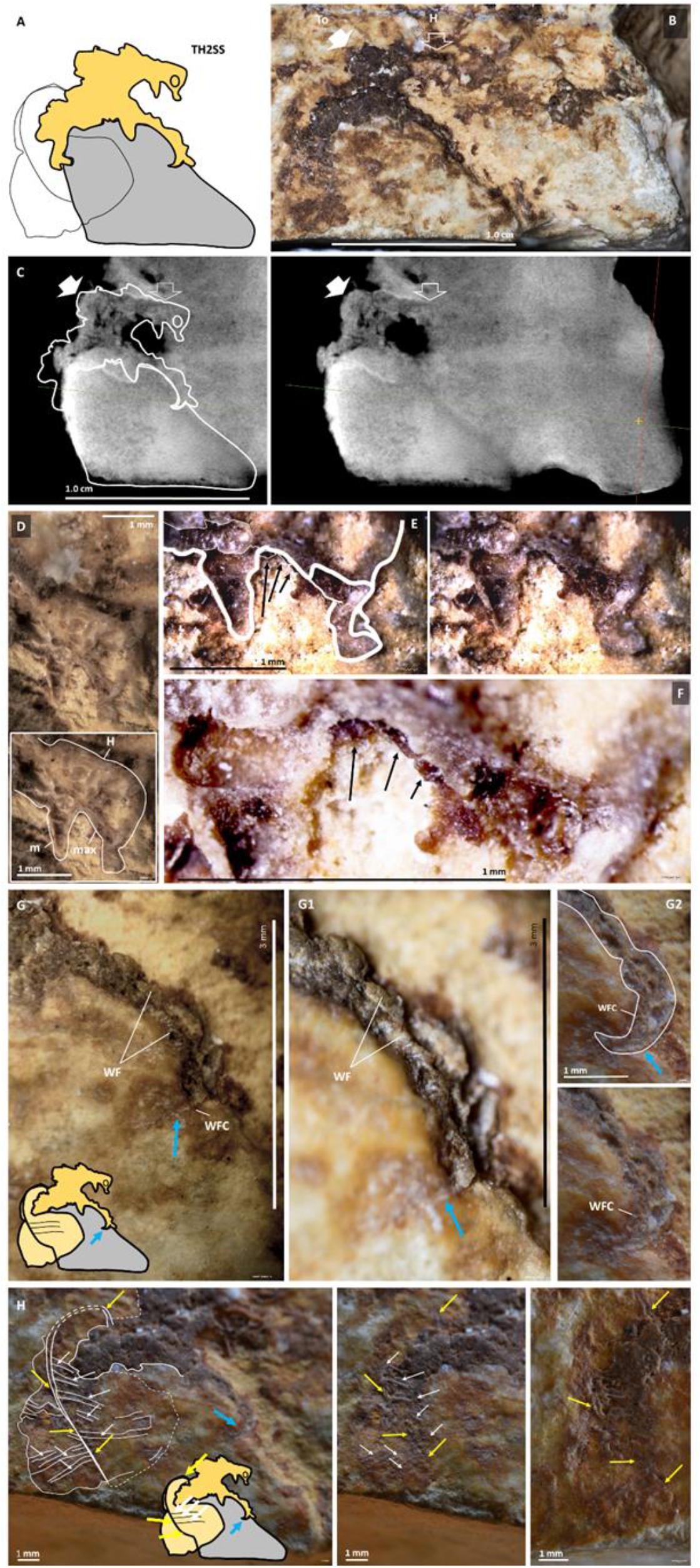
TH2SS is another hatchling from the Teylers specimen fossil, also with a beaked head and maxilla inserted teeth in addition to a forelimb claw and tail preservation. **A:** Illustration of the macrophotograph shown in **B. C:** CT scan at approximately 45 μm resolution and correlation with the illustration shown in A. Arrows are provided for guidance. The hemi-head of TH2SS exposed in sagittal view is shown under stereomicroscopy in **D.** Details of TH2SS’s hemi-head where teeth are indicated and magnified are shown in **E** and **F**. **G:** TH2SS forelimb with indication of the area (blue arrow) where the claw is located. **G1**: Anterior rim of the forelimb. **G2**: Magnified view of the claw indicated by blue arrows in G and G1. **H**: Details of TH2SS’s tail. Yellow and white arrows are provided for guidance to indicate remnants and vestiges of tail feather shafts and of the claw (blue arrow) magnified in G and G1. H – Head; m – mandible; max – maxilla; To – Torso; WF – Wing finger; WFC – Wing finger claw.

Egg remnants were identified near the distal end of Teylers specimen hindlimbs (Figure 35). Also, like in CS1 from the Berlin specimen, an egg was identified in the Teylers specimen fossil that shows linear impressions compatible with those of feather shafts (Figure 35).

**Figure 35.**
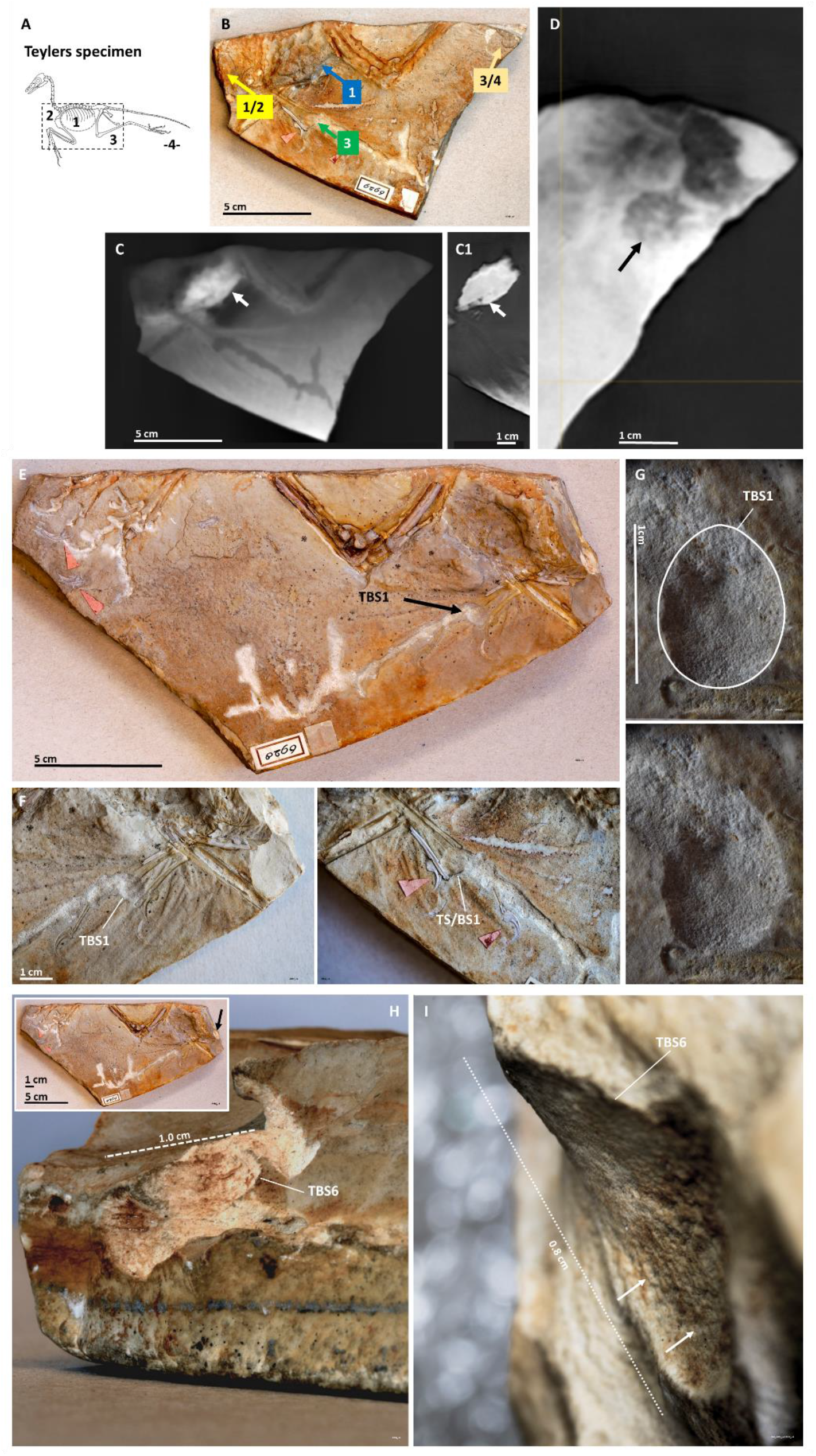
The partial skeleton of the Teylers specimen is associated with an egg mass located in its abdominal region and evidence of egg remnants near the distal end of its hindlimbs. **A:** illustration of the part of the Teylers specimen’s skeleton that was preserved shown here in accordance with the nesting posture hypothesis, together with areas equivalent to those described above for the Berlin specimen. **B**: areas homologous to those from the Berlin specimen. **C** and **C1**: coronal CT scan views of the egg mass (arrow) located in the abdominal region of the Teylers specimen. **D**: Globular structures approximately 1 cm in size (exemplified by arrow) compatible with eggs can be seen in coronal sections at the posterior region of the counterslab near the distal end of the specimen’s hindlimbs. **E**: TBS1 is a 3D egg impression located in the main slab, ventrally to the specimen’s abdominal region. **F**: TBS1 egg impression in the main slab (left) and counterslab (right; where it is designated TS/BS1). **G**: Observation of the TBS1 3D impression from the main slab under stereomicroscopy. **H:** TBS6 is a 3D egg impression located in the anterior margin of the main slab (inset on the left). Under stereomicroscopy (right), TBS6’s concave surface shows linear impressions (arrows) compatible with those of feather shafts (**I**).

The periphery of one egg remnant from the Teylers specimen that is split among the two slabs of this fossil was identified that is also observable in CT scan images (Figure 36; see also below).

**Figure 36.**
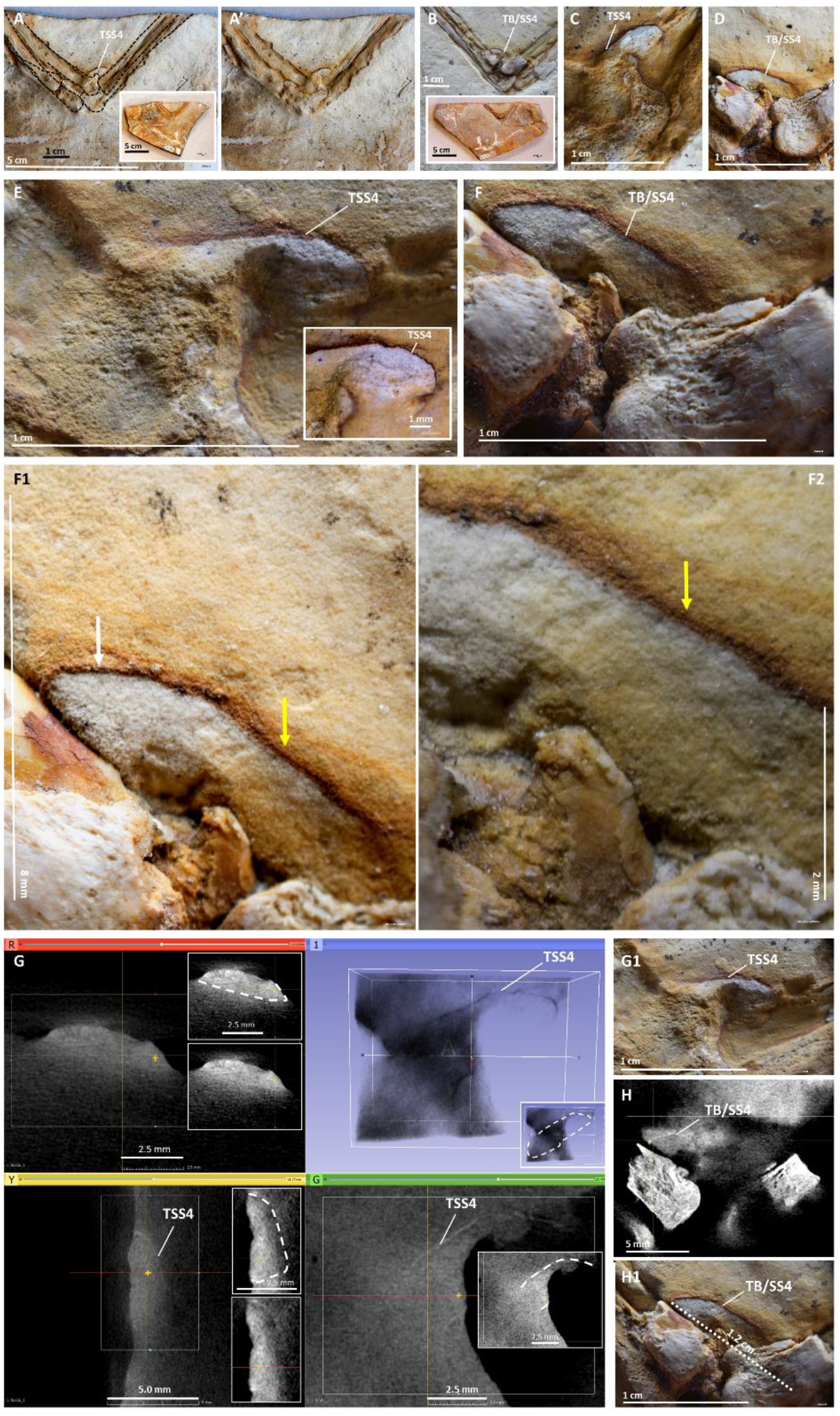
Periphery of the a Teylers specimen egg. TSS4 (<u>T</u>eylers <u>S</u>mall <u>S</u>lab egg <u>4</u>) is located dorsally to the flexed knee joint of the Teylers specimen and is split between the counterslab (shown in **A**, **C** and **E**) and the main slab (shown in **B**, **D**, **F**, **F1** and **F2**.) The part of TSS4 that is visible in the main slab is designated TB/SS4 (<u>T</u>eylers <u>B</u>ig slab counterpart of <u>S</u>mall <u>S</u>lab egg <u>4</u>). Arrows in F1 and F2 are provided for guidance in the observation of TSS4’s putative eggshell, which is shown to be visible and radiodense in the counterslab (**G**) and in the main slab (**H**). G and H were obtained via CT scan, respectively for TSS4 and TB/SS4, at approximately 45 μm resolution.

### Winged hatchlings and egg littering in the isolated *Archaeopteryx* feather fossil

Corroborating evidence of the same wing anatomy observed in eggs and hatchlings from the Berlin and Teylers fossils was observed in the *Archaeopteryx* isolated feather fossil, including a detached small wing remnant with three identifiable fingers and an egg remnant with two sets of parallel lines suggestive of the impressions of wing remiges shafts from an egg-enclosed fetus, which remiges number as twelve in each adult *Archaeopteryx* wing. In addition, convexities and concavities compatible with those of egg remnants were identified in the margins of this fossil and a putative egg vestige was identified in its back surface (Figure 37).

**Figure 37.**
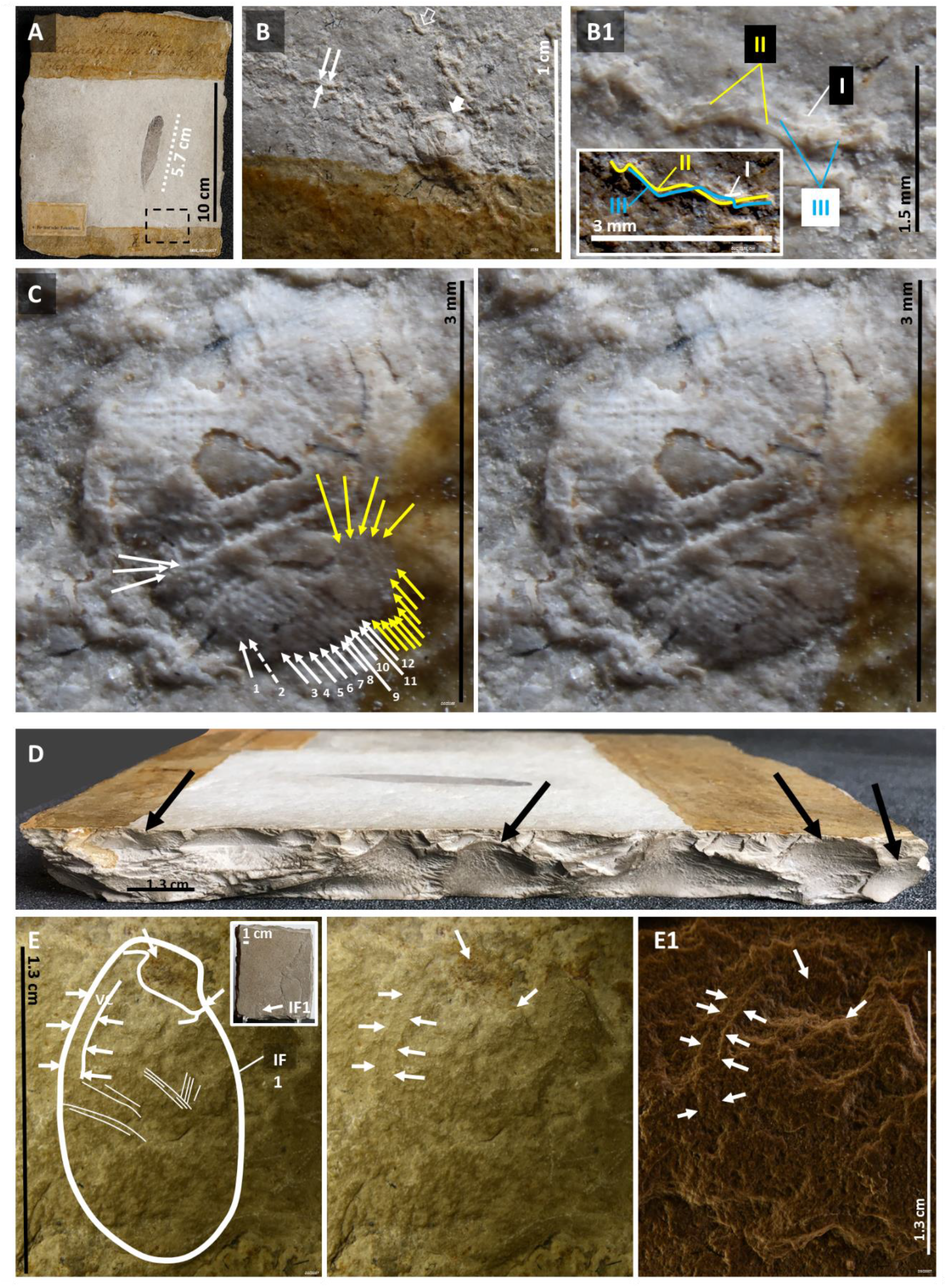
Evidence of wing remnants and egg littering in the isolated *Archaeopteryx* feather fossil. **A**: Isolated *Archaeopteryx* feather fossil. The dashed rectangle area is magnified in **B**: Wing remnants are indicated by thin white arrows (magnified in **B1**) and the open arrow. Three-fingers are identified in one wing remnant and shown in a diagram (inset). The large white arrow in B points to an egg remnant amplified in **C**, where two separate sets of parallel lines are shown and represented in different colors, white and yellow. **D**: The right side margin of the isolated *Archaeopteryx* feather fossil shows one convexity and three concavities approximately 1,3 cm long (arrows). IF1 is a 1.3 cm egg vestige in the back surface of the isolated *Archaeopteryx* feather fossil shown under stereomicroscopy using two light conditions (**E** and **E1**). Arrows are provided as visual references. VC – Vertebral column.

### Evidence of eggs in other *Archaeopteryx* fossils

The Thermopolis specimen is ventrally exposed. According to the nesting hypothesis, it would be in the counterslab, whose location is unknown to science, that eggs and egg remnants would be more likely to be found. Still, egg impressions were found near its tail and evidence of egg vestiges was also found in other areas of the available slab (Figure 38A and data not shown).

**Figure 38.**
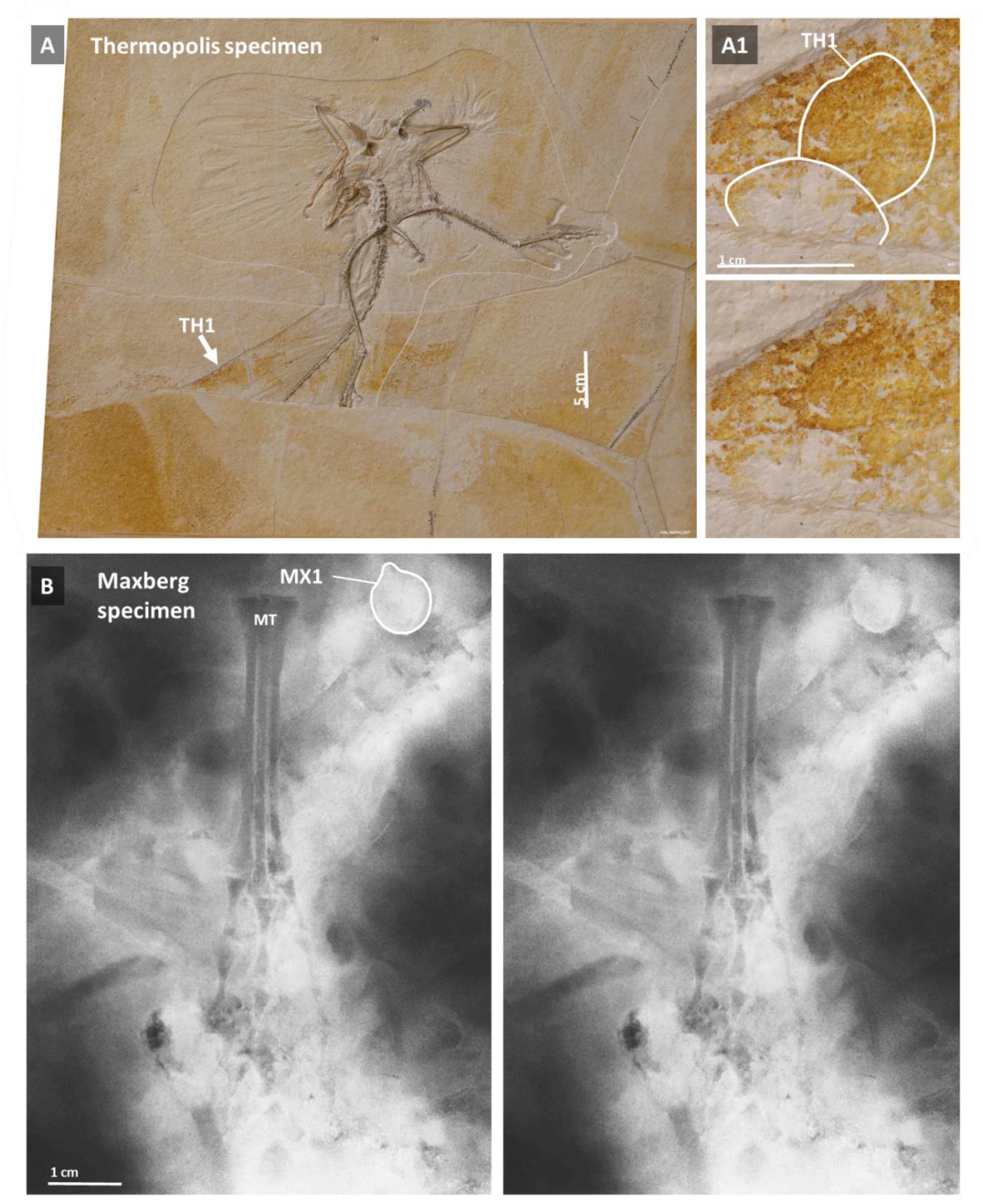
Evidence of eggs in the Thermopolis and Maxberg specimens. **A:** The main slab of the Thermopolis specimen shows impressions that are approximately 1,3 cm long, of which TH1 is an example (**A1**). **B**: Radiologic image compatible with that of an egg in the Maxberg specimen. MT – Metatarsals.

In addition, an X-ray of Maxberg specimen’s fossil shows, near one of its feet, MX1 as a radiodense globular structure approximately 1 cm in size with a papillary pole similar to that of MSB5 and MSB15 from the Berlin specimen (Figure 38B).

### Insights into the cause of death of Archaeopteryx specimens

Granular sediment, different from the very thin and agranular sediment seen everywhere else in both slabs of the Berlin specimen fossil, was observed in between its tail feathers (Figure 39).

**Figure 39.**
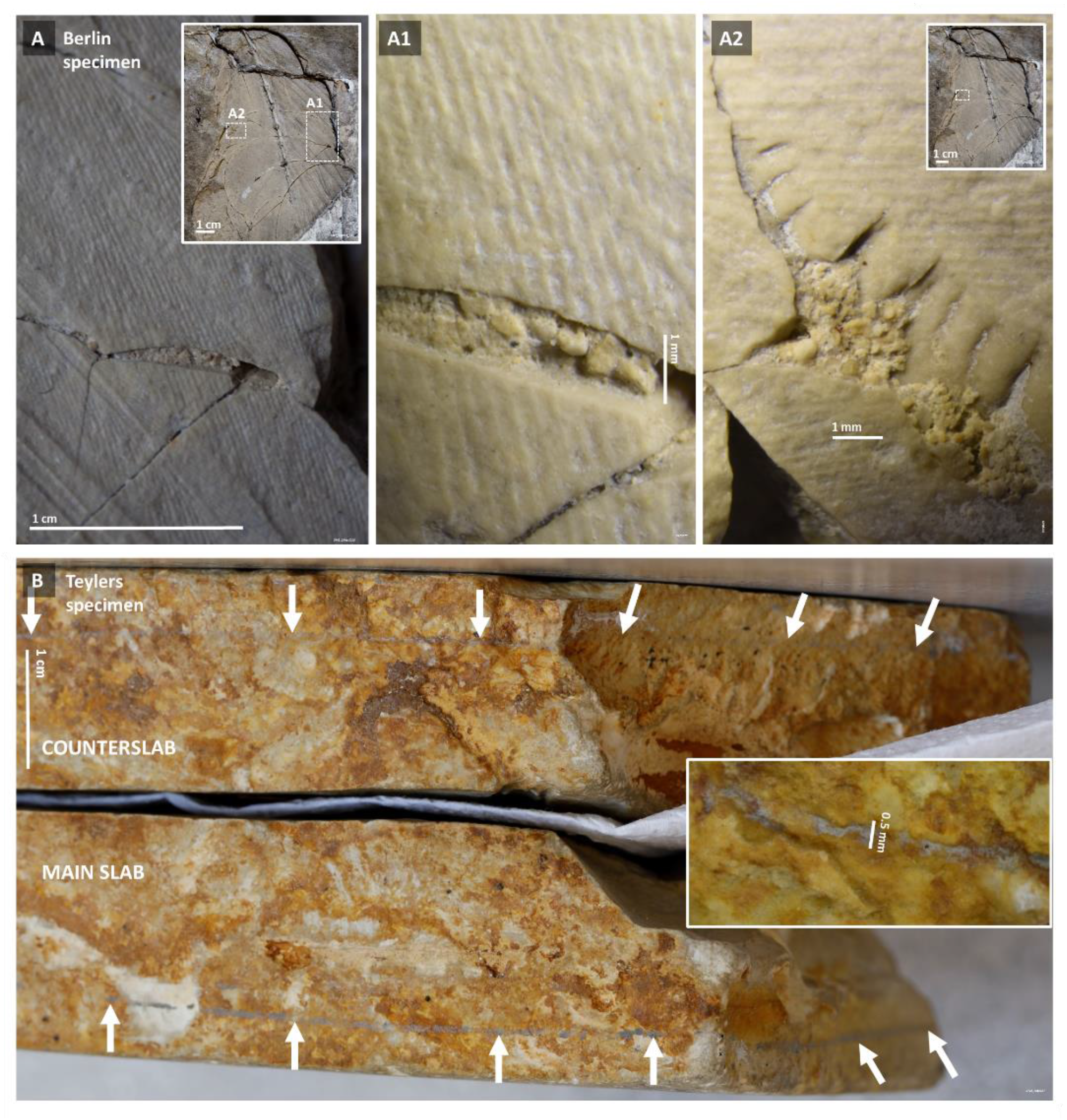
Granular sediment beneath tail feathers of the Berlin specimen and sediment lines that delimit the Teylers specimen fossil bed. **A:** Parts of a counterslab fragment that represents the tail of the Berlin specimen shown and amplified in **A1** and **A2**. **B**: Sediment lines (arrows) above and below the Teylers specimen skeleton. A magnified view of one of these sediment lines is shown in the inset.

In turn, one discrete sediment line was observed above and another line below the Teylers specimen’s skeleton (Figure 39).

### Summary of egg size, morphology and periphery from different *Archaeopteryx* fossils

The evidence of eggs detected in the Berlin, Teylers, Thermopolis, Maxberg and isolated feather fossil of *Archaeopteryx* is consistent with flattened eggs measuring 1.0 - 1.4 x 0.8 - 1.0 x 0.6 - 0.8 mm (Figure 40). Egg outlines are in agreement with the predominant egg view expected in the dorso-laterally exposed Berlin specimen and in the laterally exposed Teylers specimen.

**Figure 40.**
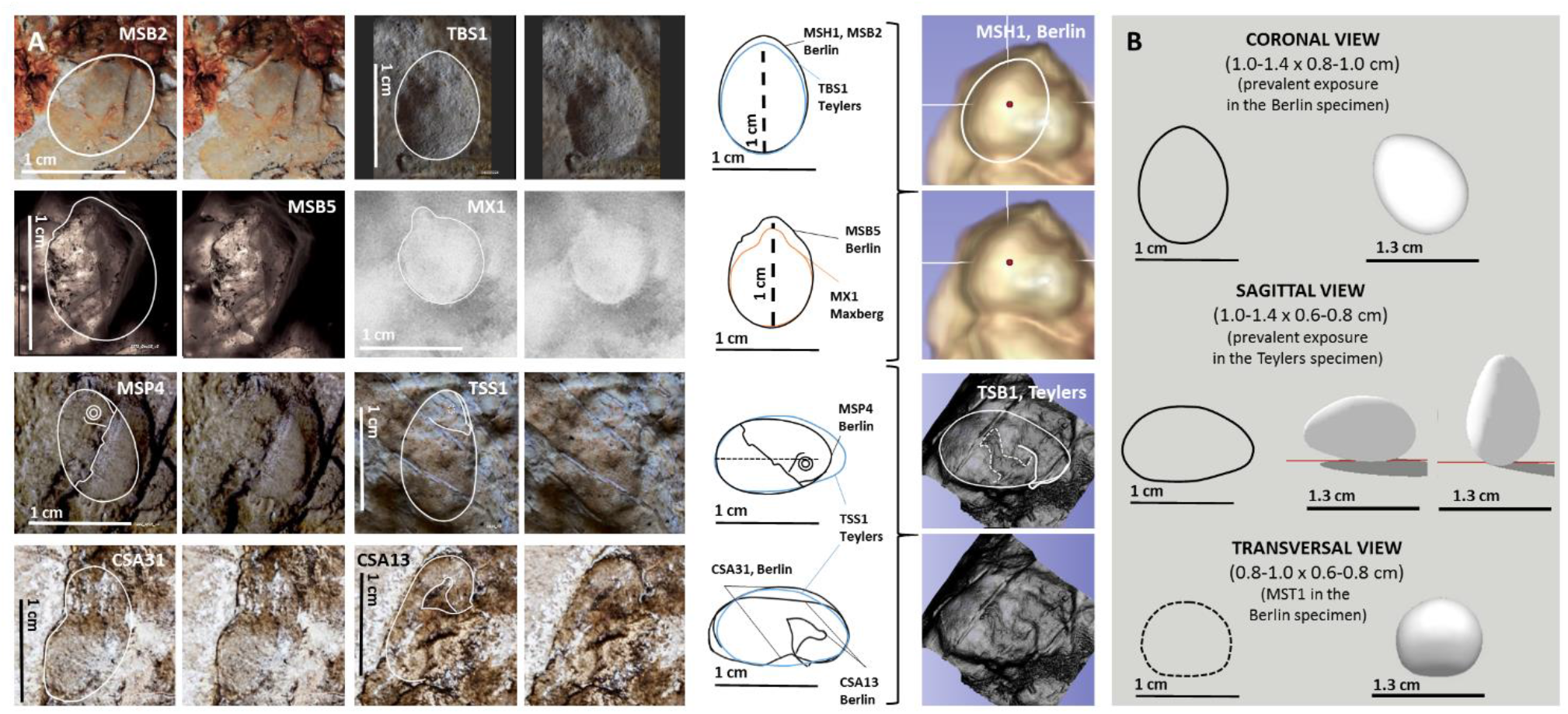
Eggs from different *Archaeopteryx* specimens have the same size and flattened morphology. **A:** Eggs in 2D (left) and 3D (center) views are shown with outlines. The 2D outlines shown in the center are in agreement with images obtained from 3D software analysis of CT scan data (center right). **B**: 2D outlines (coronal, sagittal and transversal) were used to build a computer generated 3D egg model.

The linear periphery of MSB1 and egg fragments in the Berlins specimen, and that of TSS4 from the Teylers specimen, all observable in radial view, and the exposed surface of TH1SS’s egg in the Teylers specimen and of the putative egg remnant in the isolated *Archaeopteryx* feather fossil where linear impressions suggestive of the wing remiges shafts were detected, indicate that the surface of these eggs is not nodular nor ornamented but smooth and appears to be punctuated by pores (Figure 41).

**Figure 41.**
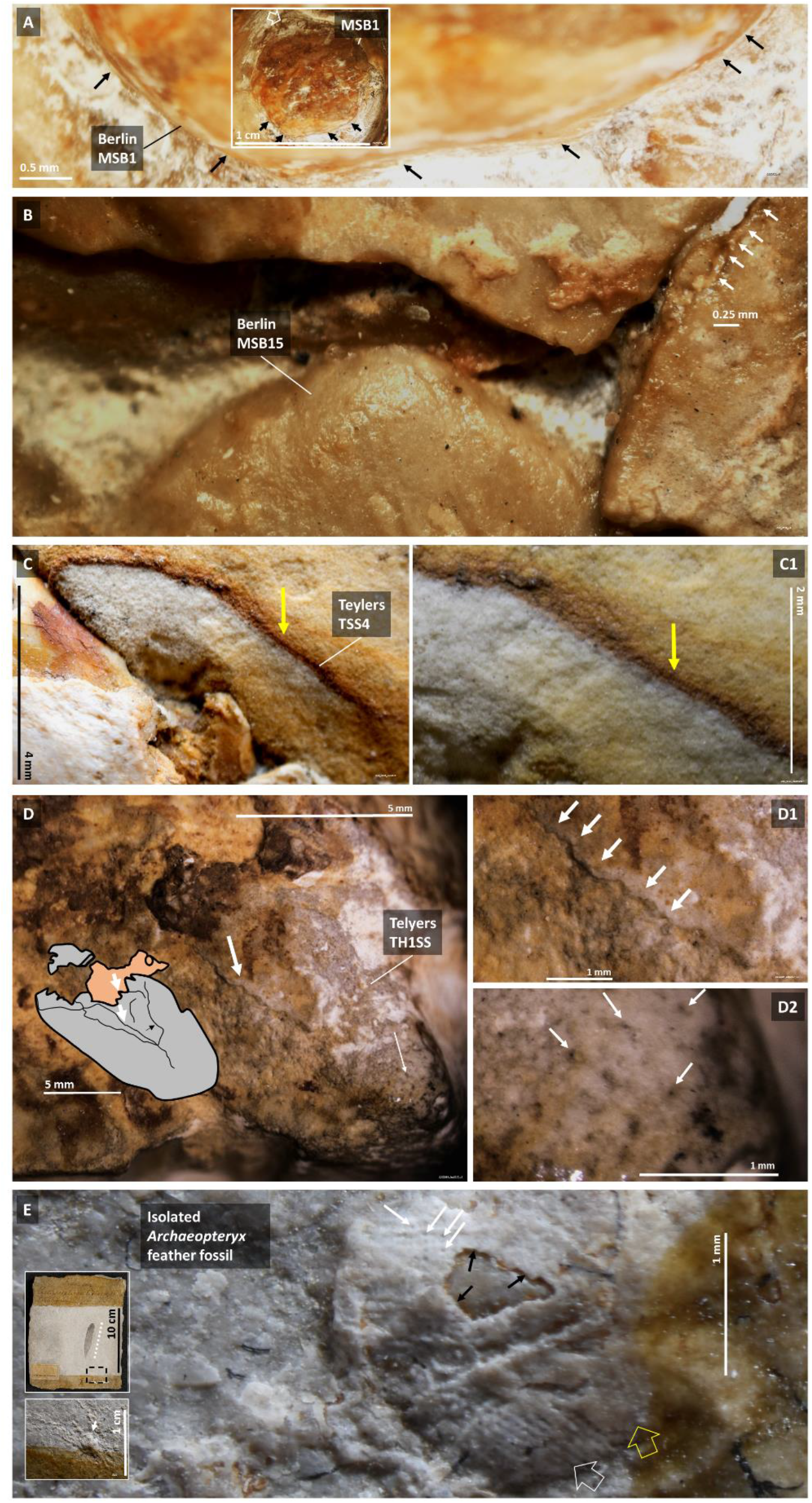
Egg periphery from different *Archaeopteryx* fossils. Egg periphery of MSB1 (**A**) and an egg fragment located in the abdominal region near MSB15 (**B**), both ventral to the Berlin specimen, and Teylers specimen’s TSS4 (**C**). **D**: TH1SS’s egg surface is shown under stereomicroscopic view with interpretation (inset). A putative wrinkle or crack is shown in **D1** and putative pores in **D2**. The surface of an egg remnant with two sets of parallel linear impressions suggestive of remiges shafts (open arrows) is shown in **E**, which shows a loss of substance (black arrows) and is punctuated by apparent pores (white arrows).

### Summary of fetal and hatchling head morphology findings from different *Archaeopteryx* fossils

A fetus with toothed beak was identified in the MSB5 egg from the Berlin specimen. Findings suggestive of beaked hatchlings include H1MS, H2MS and MSH15 in the Berlin specimen and TH1SS and TH2SS in the Teylers specimen. Teylers specimen’s TH1SS and TH2SS both present evidence of a toothed beak (Figure 42).

**Figure 42.**
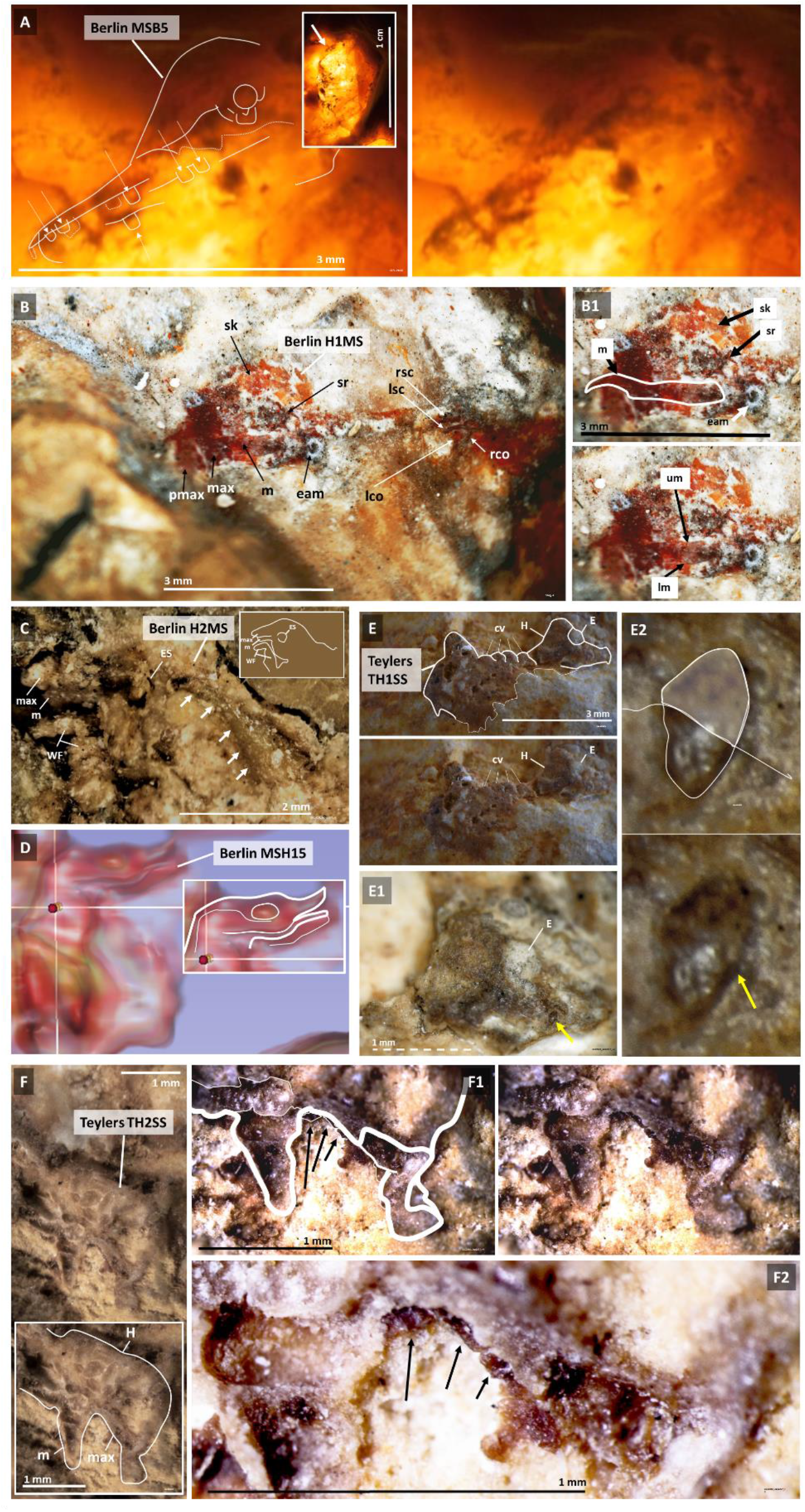
Toothed beak head morphology of *Archaeopteryx* fetuses and hatchlings. **A**: Outline of egg-enclosed MSB5 fetus’s toothed beak. **B**: Head, neck and upper thorax of H1MS from the Berlin specimen, and its magnified and outlined beaked head in **B1**. **C**: Left side and back of hatchling H2MS’s head. The non-segmented structure in the cervical area (arrows) may represent its neural tube. **D**: MSH15’s head outline detected via 3D software analysis of CT scan data. **E**: Upper body of hatchling TH1SS from Teylers specimen’s fossil and detail of its head with a maxilla inserted tooth (arrow) in **E1** and **E2**. **F**: Outline of hatchling TH2SŚs hemi-head from Teylers fossil and its maxilla inserted teeth (arrows) in **F1** and **F2**. cv-cervical vertebrae; E – Eye; eam – external acoustic meatus; ES – Eye socket; lco – left coracoid; lsc – left scapula; m – mandible; max – maxilla; pmax – premaxillary; rco – right coracoid; rsc – right scapula; sk – skull; sr – sclerotic ring; WF – wing finger.

### Summary of fetal and hatchling wing morphology findings from different *Archaeopteryx* fossils

Two egg-enclosed fetuses with evidence of finger claws (CS54 and CS1), one of which with feather shaft impressions (CS1), from the Berlin specimen and two sets of parallel impressions suggestive of wing remige shafts, supposedly one from each wing, in an egg remnant from the *Archaeopteryx* isolated feather fossil were identified (Figure 43).

**Figure 43.**
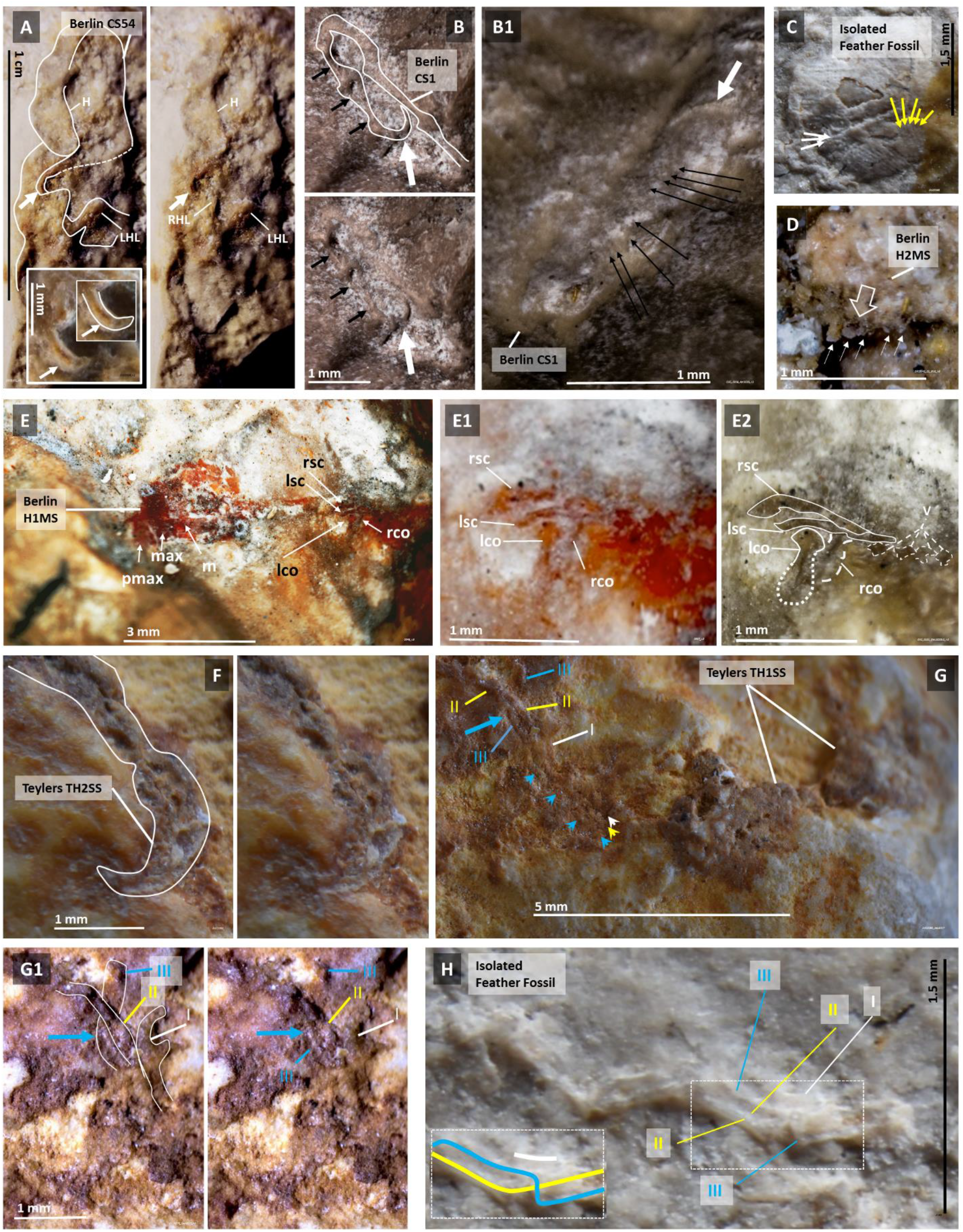
Findings suggestive of fetal feathered wings and wing anatomy in hatchlings from different *Archaeopteryx* fossils. **A:** Wing claw (arrow) in surface exposed egg-enclosed CS54’s fetus from the Berlin specimen. **B**: Claw (bigger arrow) at the distal end of a folded forelimb (smaller arrows) and impression of parallel lines (thin arrows) showing through the surface of Berlin specimen’s CS1 egg. **C**: Two separate sets of diagonal lines on the surface of an egg remnant in the isolated *Archaeopteryx* feather fossil. **D**: Feather shaft stumps (arrows) on the body surface of Berlin specimeńs H2MS hatchling. **E**: Shoulder girdle bone preservation in H1MS from the Berlin specimen. **F**: Forelimb claw from Teylers specimen’s TH2SS hatchling. **G**: Three wing fingers (I, II and III) from Teylers TH1SS hatchling, and close-up of the 3^rd^ finger under-crossing the 2^nd^ (**G1**). **H**: Three-finger wing remnant observable on the surface of the isolated *Archaeopteryx* feather fossil. co – coracoids; eam-external acoustic meatus; H – head; lco – left coracoid; lsc – left scapula; LFL – left forelimb; LHL – left hindlimb; m – mandible; max – maxilla; rco – right coracoid; rsc – right scapula; sc – scapulae.

One hatchling with two hatchet-shaped coracoids and strap-like scapulae (H1MS), and another hatchling with a parallel arrangement of wing feather shaft stumps (H2MS) from the Berlin specimen, a clawed forefinger in one hatchling (TH2SS) and a wing remnant with 3-finger anatomy and undercrossing of the 2^nd^ wing finger by the 3^rd^ in another (TH1SS) in the Teylers fossil, and one detached 3-finger wing remnant in the *Archaeopteryx* isolated feather fossil are also shown in Figure 43.

## Discussion

### The Berlin specimen of *Archaeopteryx* fossilized in its ground nest

The association of an adult bird, egg-enclosed fetuses, hatchlings and extensive egg littering make more likely the interpretation of the Berlin specimen fossil as that of a fossilized animal with its ventral side against its ground nest than that of a dead animal carried away by rain waters that, after drifting, would have sunk with its back against the floor of a Solnhofen lagoon (*1, 10*).

The originality of the slab proper of the counterslab indicates that the entire animal was fully encased, in full articulation, together with its nest environment, within the two fossil labs before fossilization occurred and became part of the Solnhofen marine floor.

The counterslab fragments include surface body parts and sediment that was probably deposited over the body of the animal at the moment of its death or shortly after. In turn, the slab proper of the counterslab detached not only from the main slab but also from counterslab fragments. Therefore, it represents sediment probably deposited some time later than that which integrates the counterslab fragments and the main slab where the Berlin specimen body resides (Figure 44 and Movie 1).

**Figure 44.**
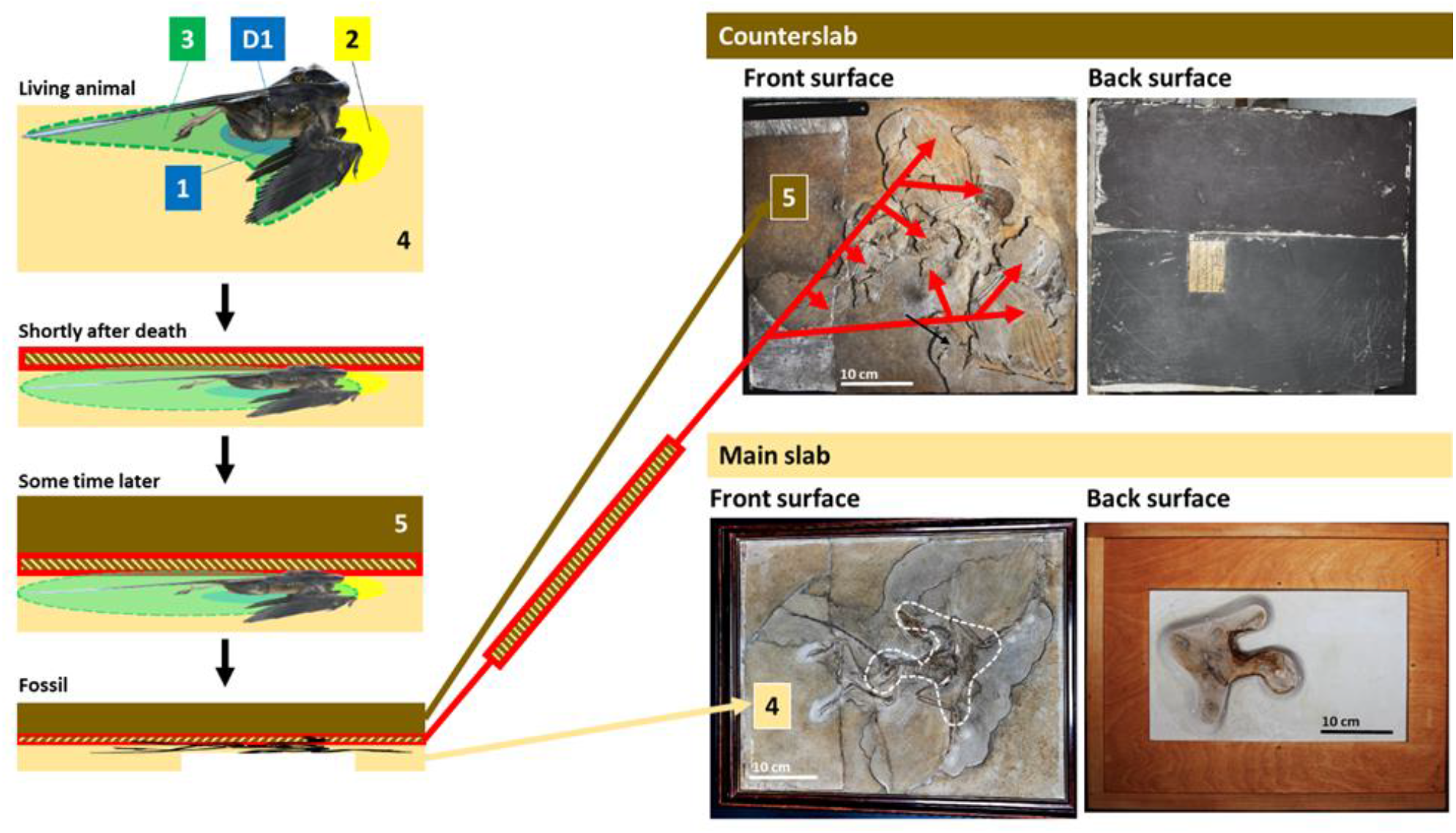
Relative position of the Berlin specimen fossil slabs based on the nesting hypothesis: the counterslab lies atop the main slab and not the other way around. Areas 1 (ventral to the specimen) and 4 (external to the perimeter of the Berlin specimen’s feathered body) are contributed solely by the main slab. In turn, Area 5 represents a layer situated above the Berlin specimen fossil bed that is contributed solely by the slab proper of the counterslab. The red rectangle with diagonal lines on the left represents a layer of sediment that would have been deposited peri-mortem over the dorsal surface of the animal, which layer corresponds to the counterslab fragments (arrows in the front surface of the counterslab). The part of these counterslab fragments that is dorsal to the animal’s torso is called Area D1. Areas 2 and 3 are each contributed by the main slab and counterslab fragments.

Aquatic ecosystems often include surfaced areas within and around them, which, similarly to tidal flats and mudflats, are subject to seasonal flooding. Similarly, Solnhofen’s marine lagoons are unlikely to have been entirely submerged at all times, and may have included permanently or periodically exposed dry areas. The Berlin specimen’s fossil location cannot be safely determined. Therefore, despite the fact that this fossil’s interpretation should take into consideration general data from its environment, this should not supersede data extracted from its own slabs.

Presumably, near water ground nesting of soft eggs resulted in periodic submersion of nesting areas in Solnhofen making difficult the differentiation of nesting areas from unexposed areas, especially in the absence of rigid eggshells and their reproducible 2D outlines, which would have been easier to recognize than the outline variations permitted by non-rigid eggs (*10*).

Reported examples of periaquatic habitats of fossilized animals include a drowned bird colony (*13*), dinosaur nests on tidal flats (*14, 15*) and a periaquatic pterosaur colonial nesting site (*16*). A*rchaeopteryx*’s reproductive behavior was presumably influenced by its dinosaurian origin (*17*), and near-shore bird reproduction of hard shelled eggs is observable in extant birds.

### The ventral mass of soft eggs, evidence of brooding and parental behavior

The lumbar-thoracic curved segment of the specimen’s backbone represents the upwards segment of a curvature that resulted from laying over an egg mass, whose downwards segment became obliterated by the postmortem overextension of the neck of the animal that dislocated thoracic vertebrae from the thoracic grid, an event that could have occurred above or under water and resulted in the animal’s opisthotonic posture (*9, 10*). However, the overextension of the neck may have occurred upon an already retroflexed neck, as seen in today’s resting and nesting birds (*9;* see below).

Only part of the Teylers skeleton is available but an egg mass ventral to its thoraco-abdominal area was identified, which is very similar in size to that of the Berlin specimen (*1, 10–12*).

Both here and in previous reports (*10,12*), fetuses in tucked head pose and with surface morphology preservation were identified within eggs measuring approximately 1.0 to 1.4 cm in the longer axis, both in the Berlin and Teylers specimens. Their size conforms to *Archaeopteryx*’s estimated 1.4 cm wide pelvic opening and fused pubis, a structure the eggs had to pass through when laid (*1,10*). These eggs are in the size range of the extant Goldcrest (*Regulus regulus*) and Azure-crowned Hummingbird (*Amazilla cyanocephala*) eggs (*18*; see below).

Instead of well demarcated hard eggshells with homogeneous 2D and 3D shapes, 2D egg outlines determined by egg orientation and molding to neighboring structures were found in the Berlin, Teylers, Thermopolis, Maxberg and isolated feather fossil of *Archaeopteryx*. *Archaeopteryx*’s eggshells were apparently less preserved than their egg content. Exceptions include MSB1’s and that from an egg fragment ventral to the abdominal area of the Berlin specimen, the periphery of Teylers TSS4 egg and the egg remnant identified in the isolated *Archaeopteryx* feather fossil, all less than 0.1 mm thick. The existence of a putative wrinkle in Teylers TH1SS and the membranous appearance of the egg remnant surface located on the front surface of the isolated feather fossil suggest that the eggs were truly soft and not just pliable, with shape distortions accentuated by compression forces.

Clearly, understanding the microstructure and chemical composition of the eggshells would be relevant but those studies were not authorized in the Berlin and Teylers specimens. The counterslab of the Thermopolis represents the surface that is ventral to the specimen, but this slab is apparently lost to science, like the Maxberg specimen for which only X-rays are available. The London specimen would be most promising for study, including the ventral side of the specimen represented by the back surface of the main slab, but access to the original fossil, was not permitted (*10*). Access to the Munich and Eichstatt specimens’ fossils was not permitted either.

If the presence of calcium suspected from CT images and white powder observed in the carbonaceous DAA and DAB of the counterslab of the Berlin specimen is confirmed, it is possible that shell units exist in the eggshell that allow for relative movement along their suture line, a mechanism previously suggested by others to explain subspherical shapes in dinosaur eggs and egg fragmentation patterns (*19, 20*).

If these were calcareous concretions instead of eggs, they would be more disparately sized and would show signs of radial growth instead of fetal morphology in their interior. They are not egg follicles either, not even if released from a ruptured abdomen following decomposition, because of their organized location underneath the thorax and anteriorly to the flexed wings of the Berlin specimen, inclusively with juxtaposition. Late stage fetal development as seen in CS1 and CS54, and a beaked fetus with teeth are also not compatible with egg follicles but are indicative of eggs. The existence of the same flattened egg outline in several fossils rules out soft eggshells due to stress (*21*) and favors natively deformable soft eggs.

Diagenesis is known to differ within the same clutch (*22*), explaining how a single fetal head with a toothed beak (MSB5 egg) and another case of hind and forelimb fetal preservation (CS54 egg) were so far identified, whereas most other fetuses appear to have only surface morphology preservation.

The intimate association of the upper body and especially both flexed wings with eggs suggests egg arrangement and brooding. In turn, the presence of hatchlings in the retrocurved neck area, a place within reach of the adult animal’s mouth and appropriate for feeding the hatchlings, possibly via food regurgitation, is compatible with parental care. The fossilized pose of H2MS suggests crawling around of the hatchlings in the nest. However, despite this apparent locomotor capability, their toothed beak and possibly even some degree of wing development, given their very small size, it is likely that the hatchlings were close to the altricial end of the spectrum of semi-precocial young (*23*).

### Convergence versus non-convergence in the evolution of hard eggshells

All extant birds are oviparous and lay hard-shelled eggs. They are believed to have derived from a maniraptoran theropod dinosaur, an animal closely related to *Archaeopteryx* (*24*), which in turn may have derived from dinosaur groups known to have laid hard-shelled eggs (*25*), despite an argument for the first dinosaurs to have laid soft eggs (*26*).

Several dinosaur eggs with rigid eggshells have been found, but only a few from pre-Cretaceous times. Soft or pliable eggs are an explanation for this (*10, 27*).

An alternative theory for the origin of birds suggests that they derive from more primitive archosaurs for which no oological data is available (*28*). However, regardless of whether it was dinosaur-bird or an earlier archosaur-bird lineage that brought us to today’s birds, the first hard-shelled egg would have arisen in the presumed common ancestor to crocodiles, dinosaurs and birds because they all produce calcareous rigid eggs, whereas an even earlier precursor would have had soft shelled eggs like some extant reptiles. Therefore, non-rigid eggs in *Archaeopteryx* poses some challenges to the phylogeny of birds unless one accepts that different reproduction strategies existed in dinosaurs and possibly even in birds (*10; manuscript in preparation*).

Both soft and hard eggshells have been described among turtles, geckos, sharks and pterosaurs (*29*). The presence of both types of eggs in different animal groups has been previously attributed to convergence (*26*). However, a more parsimonious mechanism such as heterochrony or a molecular switch acting upon multiple possibilities inherited from a common ancestor was alternatively proposed in these animals and also in birds (*10*), and it was also proposed that the adoption of different egg types may have influenced survival at the end of the Mesozoic (*10*).

### Ossification

Tiny *Archaeopteryx* hatchling bones were identified, not by their radiologic bone density properties but by their morphology via direct observation, suggesting that bone ossification may have mainly occurred after birth.

It is possible that late ossification, diffusion of skeletal elements during demineralization followed by preferential soft tissue fixation of radiodense elements during remineralization or a combination of both contributed to surface morphology of hatchlings and egg-enclosed fetuses.

In any case, faster remineralization of the egg structures than of the adult animal’s skeleton could explain the rib distortions observed atop the egg masses in the Berlin and Teylers specimens.

### Egg size, egg number and clutch size

The fact that the Berlin specimen is encased within its two slabs is most important to the effort of trying to determine its clutch size, an effort not permitted by the partial skeleton of the Teylers specimen or the lack of the other slab of the Thermopolis specimen.

The different areas of the Berlin specimen fossil defined for this study translate the attempt to correlate egg preservation and their proximity to the adult animal with the likelihood that they belonged to this animal. However, clutch size determination is difficult because not all eggs within a clutch are likely to have been preserved, egg dispersal is likely to have occurred and egg littering from a previous seasons is difficult to discern. In addition, it is possible that small counterslab fragments were lost and that the dark areas of the counterslab (DAA and DAB) represent areas of exceptional but yet incomplete fossil preservation that may have originally contained additional nest elements.

The main egg mass of the Berlin specimen is ventrally adjoined to its thorax. The eggs within the area anterior to its flexed wings are also clearly intimately associated with the adult animal. It seems reasonable to consider that at least these two egg subsets belonged to the specimen, whereas egg impressions outside its feathered body contour are more likely to have resulted from other nesting seasons due to their outer location and lower degree of preservation.

The two eggs located over its torso were probably dislodged from the area that is anterior to the animal (Area 2; see also below).

In turn, the eggs embedded in the slab proper of the counterslab must have been laid later by animals that subsequently used the same nesting site.

Excluding a few egg fragments located in the ventral side of the main slab, 135 eggs from Areas 1, 2, 3 and D1 were listed in Figure 28. 105 of these either have a directly visible surface exposed fetus in egg-enclosed position or can be detected by radiological methods (Movie 1). Two thirds of the same 135 eggs are located ventrally to the animal’s torso or just anteriorly to it, between its folded wings. Therefore, at least one hundred eggs or so are highly likely to have belonged to the Berlin specimen’s clutch (Figure 45).

**Figure 45.**
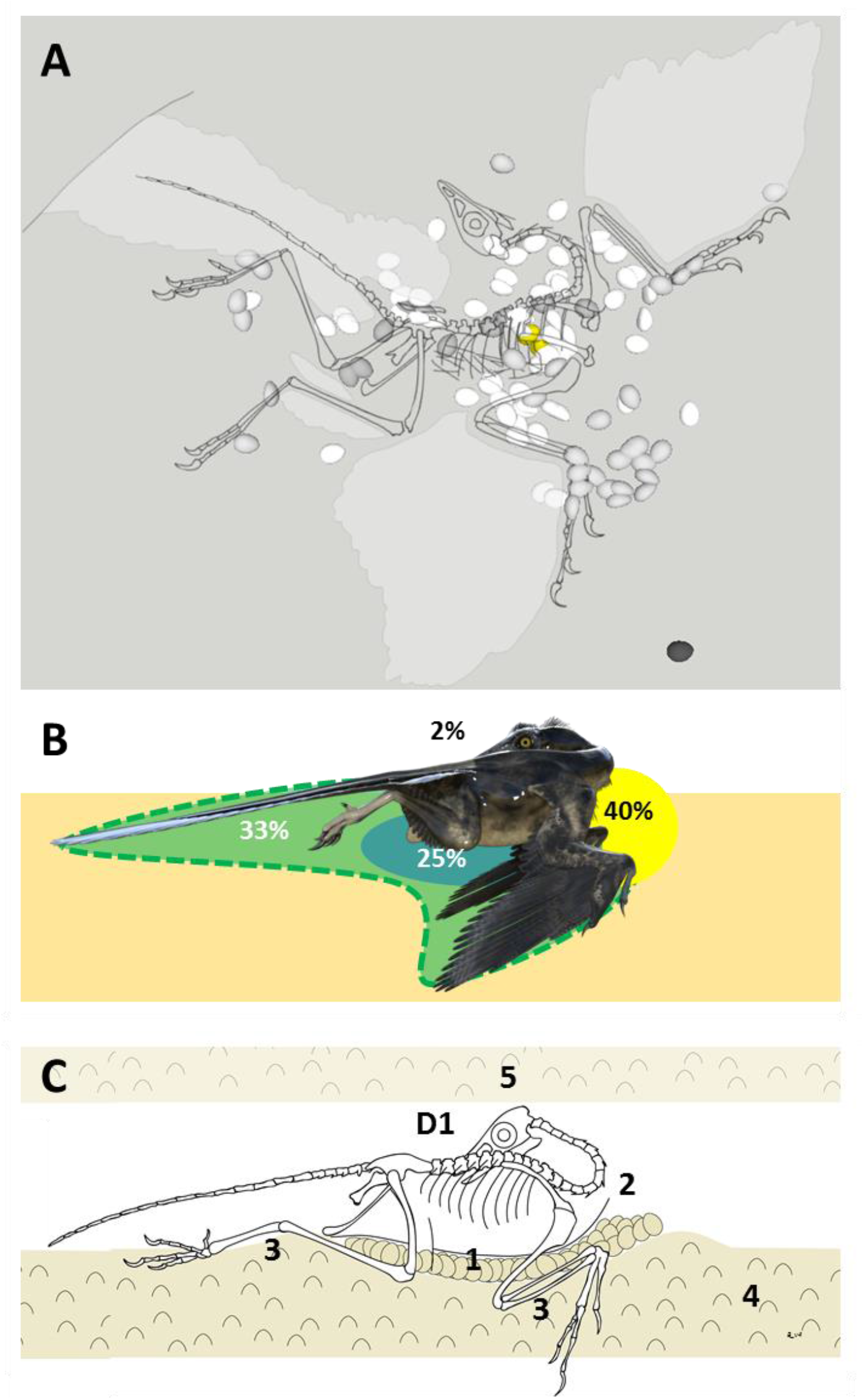
Number of eggs and their distribution in the nest. 135 Groups I, II and III eggs (**A**), their relative distribution among Areas 1, 2, 3 and D1 of the Berlin fossil (**B**) and a diagram that represents an interpretation of the egg arrangment in the living animal (**C**). Egg littering is represented in Areas 4 (main slab outside the Berlin specimen body permiter) and 5 (slab proper of the counterslab).

However numerous, given that each egg, averaging 1.3 x 1.0 x 0.8 cm in size, would weigh below 1 g, the 130 g or so weight of the resulting egg clutch for a Berlin specimen estimated to have weighed 0.8 – 1.0 Kg as a living animal (*1*), would be proportional to a 12 egg clutch from a 3.5 kg chicken with each egg weighing on average 50 g (*30*). Therefore, despite the possibility of communal nesting and the high egg number oddity reported here, which is more comparable to that of extant turtles than to the egg clutches of today’s birds, such a large clutch could be a trade-off of egg size by egg number and is actually in accordance with expected values for bird mass (*31*).

*Archaeopteryx* young were probably semi-precocial at most or altricial, and this type of organism tends to be part of a larger litters with shorter gestation times (*32*). The large number of eggs in the Berlin specimen nest and the presence of a ventral mass in addition to eggs anterior to its flexed wings is compatible with more than one wave of egg brooding and hatching in the same reproduction cycle and brooding may have been a facilitator of egg development for subsets of eggs (*11*).

### Colonial ground nesting

The presence within the Berlin specimens fossil of an egg fetus with characteristics of the adult animals (the toothed beak of MSB5), hatching animals in this fossil (MSH15) and possibly also in the Teylers fossil (TH1SS), a hatchling whose wing fingers show the characteristic crossover seen in *Archaeopteryx* from the Berlin fossil (TH2SS) and hatchlings with a fossilized tail in both fossils (H2MS from the Berlin specimen and TH2SS from Teylers), demonstrate that the association of the these specimens with their nests. In turn, the detection of egg littering demonstrates that these are ground nests.

All known *Archaeopteryx* fossils conform to variations of the same fossilized position, which is compatible with the nesting posture. This is supported by the identification of eggs, hatchlings and egg littering in the Berlin and Teylers specimen, and by evidence of eggs in the Thermopolis and Maxberg specimens. A phylogenetic analysis suggesting the Teylers specimen may be an anchiornithid (*33*) and a challenge to the *Archaeopteryx* isolated feather fossil identity (*34*) were counterweighted here by eggs and hatchlings similar to Berlin specimen’s in those fossils.

The fact that egg littering was demonstrated both in the main slab and also in the slab proper of the Berlin specimen’s counterslab indicates that the same site was used repeatedly. Inclusively, eggs embedded in the slab proper distorted the cranium of the Berlin specimen. Therefore, another animal nested over its body remains, possibly in a crowded nesting area. This type of behavior, added to the findings suggestive of egg littering in the Teylers specimen and the identification of wing and egg remnants in the isolated *Archaeopteryx* feather fossil are highly suggestive of colonial ground nesting. Therefore, all *Archaeopteryx* specimens could have been nesting at the time of their death in surfaced areas within the 60 km long Solnhofen area where they were found (*1*). The fact that multiple specimens show findings compatible with ground colony nests make tree nesting elsewhere unlikely (*10*).

### Nest structure, mechanism of death and fossil preservation

The Berlin and Teylers specimens were nesting at the time of their death. Given the similar fossilized pose of the other specimens, the possibility that they all died nesting needs to be explored. In this regard, the antero-posterior rotation of forelimbs in the Thermopolis and London specimens needs consideration (Figure 46 and Movie 1).

**Figure 46.**
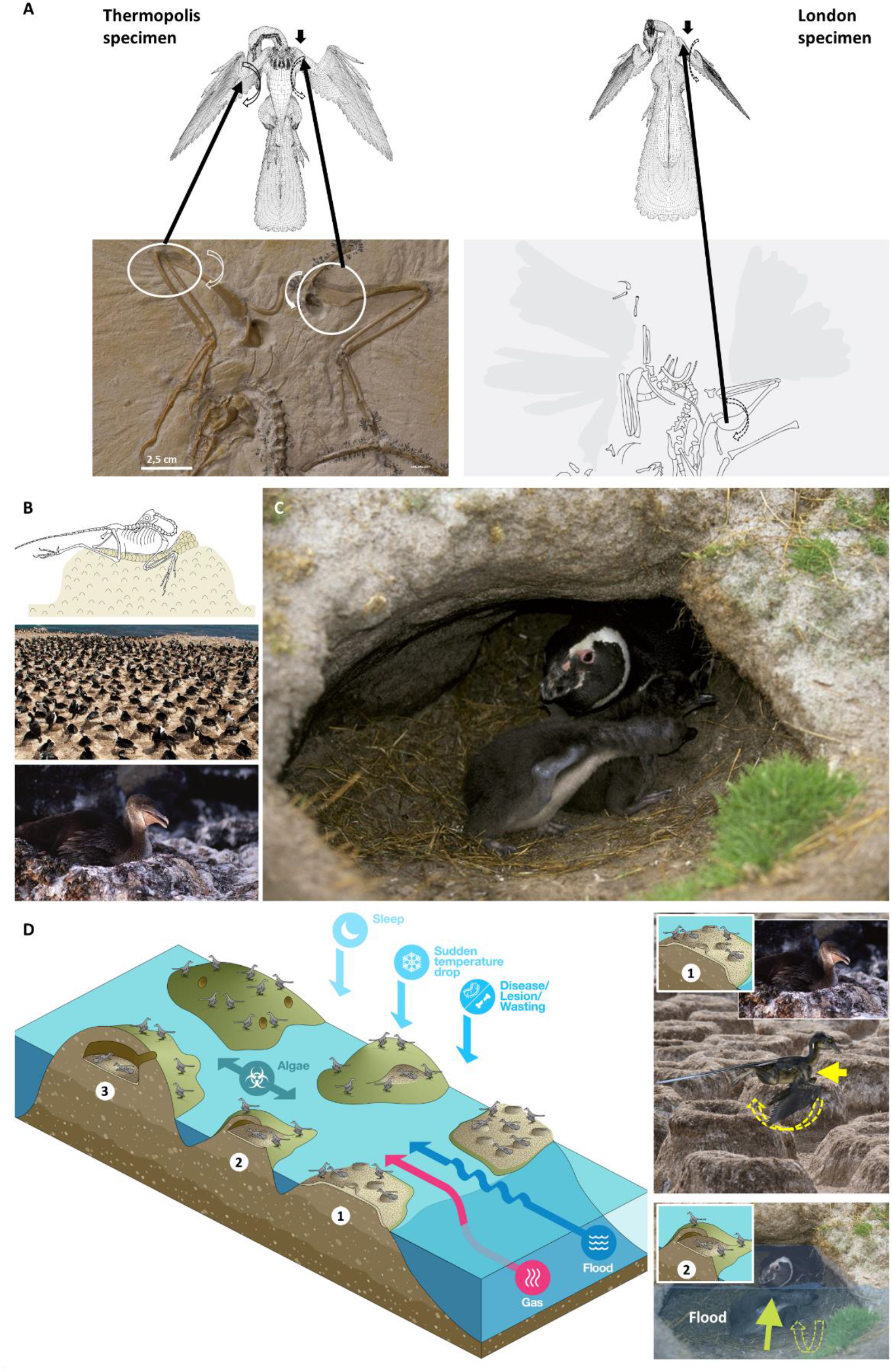
Nest structure and mechanism of death. **A**: Forelimb rotations in the Thermopolis and London specimens of *Archaeopteryx*. Two rotations in ventrally exposed Thermopolis specimen (left) and one rotation in dorsally exposed London specimen (right) are indicated. The direction of each antero-posterior rotation is indicated; those in the Thermopolis specimen occurred ventrally to the specimen and involved the entire portion of the limb distal to each rotation point; that of the London specimen occurred dorsally to the body and involved only the right humerus and not the right forearm. **B**: An elevated ground nest as one of the working hypothesis for *Archaeopteryx*’s ground nest. Diagram of a lateral view (top) and photographs of nesting colonies from extant Patagonian Cormorants (*Phalacrocorax atriceps*). The photograph on the bottom left was used with permission from Dr. Giacomo Dell’Omo and Flavio Quintana. Those on the right were licensed from Getty Images (GettyImages-10014079_medium and GettyImages-E001566_medium, respectively). **C**: Nest in a reentrance by extant penguins. **D:** Nest architecture scenarios in patches of land or sand banks near Solnhofen’s lagoons: elevated nest (1), nest in a reentrance (2) and burrow (3). Limb rotations could also have occurred in an elevated nest (right), but eggs and subsequent nesting over the Berlin specimen’s body are better explained by a flooded nest in a reentrance (bottom right). Collapse of the latter also explains the fossilized pose of hatchlings. Other possible causes of death beyond a likely flood are also represented.

Rotations can be seen in both forelimbs of the ventrally exposed Thermopolis specimen and in one of the forelimbs in the dorsally exposed London specimen. These are antero-posterior rotations that preserved the flexion of the wings at the elbow, but whereas those in the Thermopolis specimen occurred ventrally to the specimen and involved the entire portion of the limb that is distal to each rotation point, that of the London specimen occurred dorsally to the body, involved only the right humerus and was not accompanied by the rotation of the right forearm that is distal to it.

Limb rotations would be unlikely on a superficial ground nest that fossilized with specimen-egg association without complete disturbance of the nest environment, but could have resulted from water currents acting upon a cadaver located in a confined environment such as an elevated nest or a nest located in a reentrance or less likely perhaps in a burrow because limb rotations are more difficult to explain in the latter (see below). Incidentally, no reptile or known altricial bird chicks live in open nests (*35*).

On the other hand, only spatial entrapment during a flood in a reentrance would allow not only for forelimb rotations by water currents after flooding, as observed in the Thermopolis and London specimens, but also for eggs to be located over the torso of a dead animal, as observed in the Berlin specimen (Movie 1). To explain all these findings and even the fossilized clinging poses of hatchlings seen in the Berlin and Teylers specimens, the most likely explanation is the subsequent collapse of a flooded reentrance nest similar to those seen in extant penguin nearshore colonies.

The adult animal’s death could have been caused by flooding or by an event that preceded it, such as gas released from lagoons, toxic algae or other causes acting upon colony animals that, differently from most, were asleep, diseased, wasted by brooding, hampered by a temperature drop or a combination of these. Alternatively, several animals may have died but fossilization may have been rare. It is also possible that the collapse of reentrances occurred from time to time, causing the death of some nesting specimens, in areas that were subsequently flooded.

Regardless of whether flooding was the actual cause of death, it is likely to have dislodged some Berlin eggs and to have caused the limb rotations of the London and Thermopolis specimens before their nests collapsed. Granular sediment was found underneath Berlin specimen’s tail feathers and sediment lines overlie the Teylers specimen’s fossil (Figure 40), suggesting flood sediment burying, contributing to exceptional fossil preservation probably aided by phosphorus rich guano characteristic of nesting sites (*36–41*).

In addition to that of bone and teeth, fossilization of keratin-containing tissues such as feathers and skin is known to occur (*1*). The beaks of several hatchlings were partially preserved, and apparently also the palate of TH2SS and possibly even its trachea, similarly to the preservation of the syrinx in a Mesozoic bird (*11, 36*). H2MS’s body distinctly orange color warrants additional studies to determine if melanin, tetrapyrroles and other pigments can be detected (*11,37*).

As mentioned above, the fossilized clinging pose of H2MS could be explained by the collapse of the nest, favoring the possibility that the neck was already retroflexed at the time of the Berlin specimen’s death. The postmortem opisthotonic posture of the adult animal, which includes detachment of part of the vertebral column and overextension of the tail, implies that the animal could still be moved in relation to the sediment floor but either it did not disturb H2MS or the death of the adult animal and of the hatchlings occurred at different times and the nest eventually collapsed causing the still-lives described here, including H2MS’s functional contracture posture (*11, 38, 41*).

### Why these findings were missed earlier

These findings escaped earlier for approximately 140 years because: (1) *Archaeopteryx* was presumed to have sunk in a lagoon after drifting dead on the surface and the nesting hypothesis explored here was not considered earlier (*10-12*); (2) The Solnhofen environment was assumed to be entirely marine and *Archaeopteryx*’s life was believed by many to have been based on trees; (3) *Archaeopteryx* fossils have been extensively studied in relation to the evolution of birds and flights, mainly for their feathers and anatomical details but not much attention was probably given to their sediment; (4) Fossil orientation and light conditions were taken into consideration for visualization and interpretation; (5) The ventral surface of the Berlin specimen main slab, where the nest environment is represented, and the counterslab were rarely studied; (6) Transillumination was not used previously to study these fossils; (7) Instead the 2D and 3D outline of rigid eggshells, the 2D outline of a soft egg varies as a function of the way the egg is oriented to the viewer, which in turn is influenced by whether the fossilized nesting animal is exposed in ventral, dorsal or lateral view (*42*, Movie 1); (8) The process of identifying one egg that could be confidently outlined, followed by the attempt to identify eggs of similar size that could be displayed in other orientations was key to the interpretation of these fossils; (9) Despite the 1.4 cm pelvic opening of *Archaeopteryx*, the eggs were expected to be significantly longer (*1*), instead of averaging 1.3 cm in their longer axis and producing 2 cm hatchlings; (10) Careful analysis of the sediment, which is usually overlooked in detriment of bones, feathers and claws, using techniques that go far beyond mere macroscopic examination was necessary.

### Ground nesting by *Archaeopteryx* and its implications for the evolution of flight

Berlin specimen’s ground nest supports the hypothesis that wing remiges developed in association with reproduction on the ground and not primarily with flight (*10, 43*). Wings would preserve moisture necessary for the incubation of soft eggs and would also keep together, provide cover and camouflage extensive progeny. The appearance of the first feathers is probably a different issue, but it may have been in association with nesting that wings derived from forelimbs with dissociated fingers, that wing feathers elongated from filaments or relatively simple feathers and that a planar feathered wing structure developed. In accordance with this hypothesis, flight would have been a secondary capacity from feathered wings that developed from the ground up (*44*) and tree bird nests would have followed ground nests (*10, 11*).

### Other implications of this discovery

It is possible that some descriptions of microvertebrates and small dinosaur material in the absence of eggshells may correspond to soft egg remnants (*10, 22*), not only in Solnhofen but also in other Formations, and it now seems likely that Solnhofen may have been a breeding site for egg-associated Compsognathus (*7*), pterosaurs, which are more commonly found in this Formation than *Archaeopteryx*, and possibly also other animals.

The sediment needs to be reevaluated in fossil handling, preparation and study, and new techniques and methodologies are needed to determine how much is preserved in a fossil beyond bones, feathers and claws (*36–41*).

A new display position is suggested for some *Archaeopteryx* fossils to reflect the nesting hypothesis and the resulting paradigm shift in the understanding of this iconic animal (*10*).

## Acknowledgments

To Daniela Schwarz, Johannes Vogel and Thomas Schossleitner for Berlin specimen access and support, Pedro Sá for illustrations and help with 3D software modelling, Paulo Matos for help with movie production, the author’s family, ALERT’s staff, Pascal Godefroit, Dorothy Yen and Alice Wong for support, Richard Deng, Qiang Sun, Bárbara Guimarães, Ricardo Ribeiro, Ricardo Couto and Helena Gonçalves for collaboration, Helmut Tischlinger for UV photographs, Thomas Hildebrandt and Guido Fritsch for DICOM CT scan files, Peter Wellnhofer for his *Archaeopteryx* book and, with Friedrich Pfeil, Maxberg specimen’s X-rays, Herman Voogd for Teylers specimen access and support, and Aurora Peixoto for manuscript reviews.

## Data and materials availability

The Berlin specimen and the isolated *Archaeopteryx* feather fossils are kept at Naturkundemuseum Berlin, in Germany, the Teylers specimen is kept the Teylers Museum in Haarlem, Netherlands, and the Thermopolis specimen is kept at the Dinosaur Center, in Thermopolis, Wyoming, USA. Data that supports this study is available from the author upon reasonable request.

## Supplementary Materials

- Movie 1 is submitted in attachment to this manuscript and is available for download at: https://oc.alert-online.com/index.php/s/uAY9QHhgfBS7SAz
- An exploration tool of outlined fossil findings was made available at: www.alert-online.com/research/#explore/

